# Genome-wide mapping of cyclic AMP receptor protein binding in Enteroaggregative *Escherichia coli* reveals targeting of virulence-associated genes

**DOI:** 10.1101/2025.05.09.652842

**Authors:** Munirah M. Alhammadi, Joanne Hothersall, Georgina S. Lloyd, Sophie V. Titman, Thomas Guest, Douglas F. Browning, David C. Grainger, Stephen J.W. Busby, James R.J. Haycocks

## Abstract

Bacterial pathogens use a wide array of virulence factors to colonise and subsequently elicit disease in their host. These factors are often subject to extensive regulation at the transcriptional level, to ensure that their expression is timely. Although many pathogens use bespoke transcription factors that primarily target virulence genes, global transcription factors also sometimes play a role in controlling these genes. Enteroaggregative *Escherichia coli* (EAEC) is a significant cause of watery and mucoid diarrhoea globally. The organism colonises the small intestine before producing toxins that elicit disease, using a multitude of virulence factors that are encoded both chromosomally and on virulence plasmids. In this work, we have studied the cAMP Receptor Protein (CRP), a well-characterised bacterial global transcription factor, focusing on its role in pathogenicity of the prototype EAEC strain 042. We show that, although most functional CRP binding sites on the chromosome are conserved between *E. coli* K-12 and 042, CRP has been co-opted to couple the expression of some virulence genes to the nutritional state of the cell. We report novel mechanisms for CRP-dependent regulation of genes, whose products contribute to adhesion, production of a bacterial antibiotic, and export of a polysaccharide capsule.

## INTRODUCTION

Enteroaggregative *Escherichia coli* (EAEC) is a common cause of diarrhoea in adults and children in both industrialised and developing countries (1–3). The organism binds to the brush border of the small intestine and forms a distinctive “stacked brick” structure (4). Various proteins, including adhesins and toxins, have been identified as key virulence factors, though it is clear that there is enormous variation from one EAEC strain to another (5,6). These factors are encoded by chromosomal segments that are absent from harmless laboratory strains of *Escherichia coli*, such as the MG1655 K-12 strain, which many researchers use as a reference strain (7).

The expression of EAEC virulence genes is tightly regulated and many of these genes are located on large virulence plasmids. Hence the prototype EAEC 042 strain, that is often adopted as a paradigm (7), contains the pAA2 plasmid that encodes a transcription activator, AggR, which has been found to be the ‘master’ regulator of virulence (8,9). Transcript initiation at many promoters that control expression of 042 virulence genes, both on the pAA2 plasmid and on the 042 chromosome, is dependent on AggR, which is a member of the well-studied AraC/-XylS family of transcription activators (9,10). In previous papers, we reported the organisation of several AggR-dependent-promoters, and we suggested that AggR activity is regulated by a stochastically triggered feed-forward loop (10,11). In parallel research, we studied the expression of the EAEC 042 plasmid-borne *pet* gene that encodes a secreted toxin (known as Pet: plasmid-encoded toxin) that is important for successful EAEC infections and is harmful to human hosts. We were surprised to find that *pet* gene expression is independent of AggR, but, rather, depends on the cyclic AMP receptor protein (CRP) (12,13). Recall that *Escherichia coli* and related bacterial species exploit hundreds of different transcription factors to couple gene expression to changes in their environment. Whilst some of these, such as the lactose operon repressor, bind to a small number of DNA targets, others, such as CRP bind at hundreds of targets, and are global regulators of gene transcription (14,15). The activity of CRP is controlled by the level of 3’5’ cyclic adenosine monophosphate (cAMP) which, in general, in enteric bacteria, rises in response to certain stresses, such as starvation (16,17).

In more recent work, we identified the *pic* gene, located on the EAEC 042 chromosome, as a second virulence determinant whose transcription depends on CRP rather than AggR (18). This gene encodes a secreted mucinase, that was originally labelled as ‘<u>p</u>rotein involved in <u>c</u>olonisation’ (19). Encouraged by these results, we have now used chromatin immunoprecipitation in combination with high throughput DNA sequencing (ChIP-seq) to identify the full complement of DNA sites for CRP in EAEC 042. As expected, we found hundreds of targets, and most are common with the MG1655 laboratory strain. However, after triage, we identified a small number of targets that are not found in MG1655. Amongst these, we found three new targets where CRP binding directly affects transcript initiation, and these are reported here. We suggest that this argues for a role for CRP during EAEC colonisation and infection.

## METHODS AND MATERIALS

### Strains, plasmids and oligonucleotides

Strains, plasmids and oligonucleotide primers used are listed in Table S1. ChIP-seq was exploited to identify gene regulatory regions in EAEC 042 that are targets for CRP. Using PCR and primers listed in Table S1, these regions were amplified and flanked with *Eco*RI and *Hind*III to facilitate cloning into the *lac* expression vectors, pRW224 or pRW50, which were then used to measure promoter activities in strains M182 and M182Δ*crp*, exactly as in our previous studies (12,13). Differences found in M182Δ*crp* were complemented by plasmid pDCRP and derivatives. For *in vitro* studies, *EcoR*I-*Hind*III fragments carrying different gene regulatory regions were cloned into the pSR ‘holding’ plasmid. For the purposes of plasmid maintenance, strains harbouring pSR and pDCRP were grown in media supplemented with 100 μg/ml ampicillin, for strains harbouring pRW224 and pRW50 media contained 15 μg/ml tetracycline.

### DNA constructs

DNA fragments used for EMSA experiments were generated by PCR using *Escherichia coli* 042 gDNA as template using primers listed in Table S1. The p*mchA* construct was generated by cloning the 619 bp p*mchA*_600 construct *Eco*RI/*Hind*III into pRW50, which served as an intermediate template for generation of the shorter p*mchA* construct by site-directed mutagenesis (SDM), using primers p*mchA*_200 SDM F/R. For cloning *Eco*RI/*Hind*III into pSR, primers p*mchA*_200 F and p*mchA* R were used to amplify the 213 bp p*mchA* fragment. The p*mchA* p37G, p11G and p11A mutations were created by SDM, using the p*mchA* construct cloned into either pRW50 or pSR as a template. Promoter fragments sub-cloned between pRW50 and pSR were amplified with primers p*mchA* F/p*mchA* R and cloned *Eco*RI/*Hind*III. To make the p*mchA* 14A mutant, two separate “arms” were generated (using p*mchA* SOE 14A left arm F/R and p*mchA* SOE 14A right arm F/R primers), which were spliced together by overlap extension PCR to generate the final product. This was then cloned *Eco*RI/*Hind*III into pRW50.

Mutations to the 042 p0536 promoter construct were introduced using SDM, except for the p0536 38T 36A construct, which was generated by PCR using overlapping oligos, then cloned *Eco*RI/*Hind*III into pRW50. The p*kpsM*_201 DNA fragment was generated using p*kpsM*_201 F/R primers. p*kpsM*_440 was constructed using primers p*kpsMII*_440 F/R, with mutations being introduced using p*kpsM* P1-F and p*kpsM* P2-R.

All plasmid constructs were confirmed by Sanger DNA base sequencing.

### ChIP-seq

Chromatin immunoprecipitation experiments were carried out in duplicate, using the method of Middlemiss *et al.*(20) with modifications. EAEC 042 overnight cultures were sub-cultured into 40 ml of fresh LB media and grown to mid-log phase (OD_600_= 0.4-0.5), and were crosslinked in 1 % *v/v* formaldehyde for 20 minutes. The reaction was quenched by adding glycine to 0.5 M, and cells were washed in an equal volume of cold 1x TBS. Lysis was carried out by re-suspending cell pellets in 1 ml immunoprecipitation buffer (consisting of 50 mM HEPES-KOH pH 7.5, 150 mM NaCl, 1 mM EDTA, 1 % Triton-X 100, 0.1 % sodium deoxycholate, with 1x EDTA-free protease inhibitor cocktail (Roche) added per 50 ml) containing 2 mg mL^-1^ lysozyme at 37°C for 30 minutes. Samples were chilled on ice, prior to sonication using a Bioruptor plus instrument (using the high intensity setting, for 30 cycles, 30 secs on, 30 secs off). Lysates were clarified by centrifugation and immunoprecipitation was carried out by adding anti-CRP antibody (clone 1D8D9, purchased from Biolegend), with overnight incubation at 4°C on a rotating wheel. Immunocomplexes were immobilised by adding 15 μl each of washed Protein A and Protein G Dynabeads (Invitrogen) for 1 hour at 4°C on a rotating wheel. Samples were then washed 6 times with immunoprecipitation buffer, once with high salt immunoprecipitation buffer (containing 500 mM NaCl), once with ChIP wash buffer (20 mM Tris HCl pH 8.0, 250 mM LiCl, 0.5 % IGEPAL CA-630, 0.5 % sodium deoxycholate and 1 mM EDTA), and once with TE buffer. DNA was eluted in buffer containing 50 mM Tris-HCl pH 7.5, 10 mM EDTA, and 1 % SDS. To decrosslink, 5 μl of 10 mg mL^-1^ proteinase K (Sigma Aldrich) was added to the eluted DNA, and samples were incubated at 42°C for 1 hour, followed by incubation at 65°C for 5 hours. DNA was purified using a Qiagen PC purification kit, and libraries were prepared using an NEBnext Ultra II DNA library preparation kit. Library size distributions were checked using an Agilent Tapestation instrument (using High-sensitivity DNA Screentapes), and were quantified by qPCR using an NEBnext library quantification kit. Libraries were pooled, and sequenced by Azenta Life Sciences (Next Generation Sequencing Services) on a MiSeq instrument (paired end, 2 x 150 bp configuration). Sequencing data have been submitted to ArrayExpress and is available under accession code E-MTAB-14748.

### Bioinformatics

Reads were aligned to the EAEC 042 chromosome and pAA2 plasmid (references genome accession numbers NC_017626.1 and NC_017627.1 respectively) using Bowtie2 (21). Following alignment of Illumina reads to the reference genomes, MACS2 software (22) was used to identify peaks enriched in immunoprecipitated samples, compared to the mock controls (a cut-off value of q= 0.05 was used), and called peaks were visually inspected using the Artemis genome browser. The R package ChIPseeker was used to identify the nearest open reading frame to peak centres (23). To identify a binding motif, 150 bp of the DNA sequence flanking each peak centre was extracted and submitted to MEME suite using the MEME tool (24).

### β-galactosidase assays

β-galactosidase assays were done using the method of Miller (25). Overnight cultures of strain M182 (and the Δ*crp* derivative) were sub-cultured into fresh LB (with appropriate antibiotics) and grown to mid log phase (OD_600_= 0.3-0.6) aerobically with shaking at 37°C.

### Electrophoretic mobility shift assays (EMSAs)

DNA fragments were generated by excision from a verified plasmid construct, using *Eco*RI/*AatII* and *Hind*III restriction enzymes. Each fragment was end-labelled using ^32^P-γ-ATP (purchased from Hartmann Analytic GmbH) and T4 polynucleotide kinase (NEB). EMSA reactions were carried out as previously described (18). For each reaction, 0.5 ng of the ^32^P-radiolabelled DNA fragment was incubated with or without CRP in buffer containing 20 mM 4-(2-hydroxyethyl)-1-piperazine ethane sulphonic acid (HEPES) (pH 8.0), 5 mM MgCl2, 50 mM potassium glutamate, 1 mM dithiothreitol (DTT), 0.2 mM cAMP, 5 % (v/v) glycerol, 25 μg mL^-1^ herring-sperm DNA and 0.5 mg mL^-1^ bovine serum albumin. Reactions were incubated at 37°C for 30 mins before being run on a 6 % polyacrylamide gel that was visualised using a Bio-Rad PMI Personal Molecular Imager.

### Potassium permanganate footprinting

Potassium permanganate footprinting was carried out as previously described (18). DNA carrying the promoter was purified from *Hind*III/*Aat*II digested pSR, containing the cloned *Eco*RI/*Hind*III fragment, and labelled at the *Hind*III end with γ^32^P-ATP. Each reaction (20 µL) contained c. 3 nM labelled DNA fragment in 20 mM HEPES (pH 8.0), 5 mM MgCl_2_, 50 mM potassium glutamate, 1 mM dithiothreitol (DTT) and 0.5 mg mL^-1^ bovine serum albumin, and 0.2 mM cyclic AMP, 100 nM CRP and 50 nM RNA polymerase as required. Footprinting reactions were incubated at 37°C for 30 min, and then treated with 200 mM potassium permanganate for 4 min. Reactions were then stopped with 50 µL of stop solution (3 M ammonium acetate, 0.1 M EDTA pH 8.0, 1.5 M β-mercaptoethanol). Following phenol– chloroform extraction and ethanol precipitation, samples were resuspended in 40 µl 1 M piperidine and incubated for 30 min at 90°C. Samples were again purified by phenol–chloroform extraction and ethanol precipitation then resuspended in 8 µl loading buffer (95% formamide, 20 mM EDTA pH 8.0, 0.025 % bromophenol blue, 0.025 % xylene cyanol FF). Samples were analysed by 6% denaturing gel electrophoresis. Gels were calibrated with a Maxam–Gilbert ‘GA’ sequencing reaction of the labelled fragment and visualised with an Amersham Typhoon 5 biomolecular imager.

### In vitro transcription

*In vitro* transcription reactions were carried out using the method of Kolb et al.(26). Template DNA (used at a concentration of 16 μg mL^-1^) was incubated with or without CRP in buffer containing 40 mM Tris-Acetate pH 7.9, 10 mM MgCl_2_, 1 mM DTT, 100 mM KCl, 100 μg mL^-1^ BSA, 200 μM ATP/CTP/GTP, 10 μM and 5 μCi α^32^P-UTP for ten minutes at 37°C. Reactions containing CRP were supplemented with 0.2 mM cAMP. To start the reaction, σ^70^-containing RNA polymerase holoenzyme was added at a concentration of 50 nM for ten minutes at 37°C. Reactions were stopped by the addition of stop solution (95 % formamide, 10 mM EDTA pH 8.0, 0.025 % bromophenol blue and 0.025 % xylene cyanol FF). Reactions products were separated on a 6 % denaturing acrylamide gel, and were visualised using a Bio-Rad PMI Personal Molecular Imager.

### Proteins

CRP was purified using the Ghosaini method (27). RNA polymerase holoenzyme was purchased from NEB.

## RESULTS AND DISCUSSION

### ChIP analysis of CRP binding targets in EAEC 042

RegulonDB, the leading transcription regulation database for *Escherichia coli* lists over 300 transcription units where CRP is involved in regulation (28). In most cases, CRP binds to one or more specific target sequences at the regulatory region of the unit, and, alone, or in combination with other transcription factors, either activates or represses transcript initiation. In previous work, working with laboratory strains of *E. coli*, ChIP was applied directly to measure CRP binding, independent of any regulatory function (14). These studies confirmed hundreds of targets but, surprisingly, also identified targets outside of gene regulatory regions where CRP bound, apparently, with no ensuing effects on transcription (29). Prompted, by our observation that the promoters of the EAEC *pet* and *pic* genes are dependent on CRP for transcript initiation, we applied ChIP to map sites at which CRP is bound in EAEC 042 growing in mid-log phase in LB media. Figure 1a illustrates the distribution of the 322 sites on the EAEC 042 chromosome and the 10 sites on the virulence plasmid: these are listed in Table S2, along with their location with respect to the nearest coding region. We used MEME to deduce a CRP-binding logo from DNA base sequences centred on each peak (Figure 1b). As expected, this motif consists of two copies of the 5-base pair element 5’-TGTGA-3’, organised as an inverted repeat, separated by 6 base pairs. Note that this corresponds with previously derived logos for CRP (30). Inspection of the location of these targets showed that, as in previous work, nearly 1 in 5 of the DNA sites for CRP is at an intragenic location and is unlikely directly to be associated with any transcriptional regulation (Figure 1c)(14,31) .

**Fig. 1.**
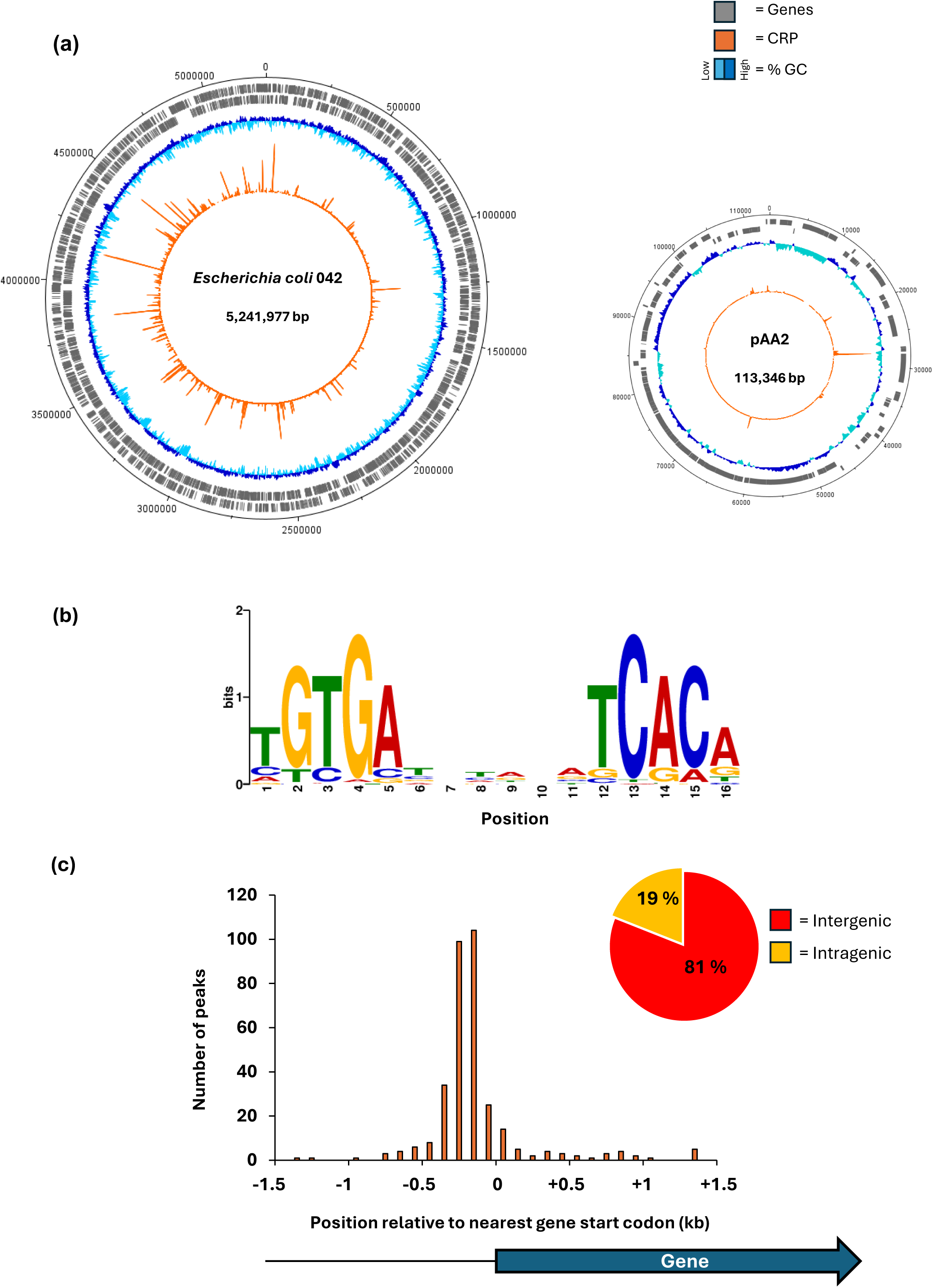
Genome-wide distribution of CRP as determined by ChIP-seq. (a) The circular plots show binding of CRP across the EAEC 042 chromosome and plasmid pAA2. Genes are shown as grey blocks, the GC-content is shown as a dark blue and cyan plot, and CRP binding is shown as an orange plot. (b) Binding motif obtained from 332 CRP peak sequences using MEME suite (24). (c) Histogram showing positions of CRP binding motifs relative to gene start codons. Four peaks were excluded that fall outside the range of the x-axis. The pie chart shows the proportion of CRP binding sites that are located within genes, or in intergenic regions.

In order to minimise false positives, we applied a cut-off to our data for targets assigned a score (during peak calling) of 100 or more: this reduced the number of targets to 150 on the EAEC chromosome and 5 on the plasmid. Inspection of these revealed that 9 are located in chromosome segments that are absent from the reference MG1655 genome, whilst 117 located to regulatory regions that have been annotated in MG1655, with CRP identified as a regulatory factor. For each of these 117 targets, we compared the MG1655 base sequence from position -80 to +20 with respect to the transcript start (according to RegulonDB) with the EAEC 042 sequence (see Table S3). Identical sequences were found at 55 targets and only 3 targets showed more than 3 base differences. We found single base insertions, upstream of the promoter -35 region in the *serC* and *pckA* regulatory regions.

### CRP interactions at uncommon EAEC 042 targets

Table 1 lists the 9 EAEC 042 chromosomal targets not found in MG1655, together with the 5 targets on the plasmid. To validate these targets, we used electromobility shift assays (EMSA) using purified CRP and ^32^P-end labelled fragments covering each regulatory region (generated by PCR, see Table S1). We did not test the 2670 and *set1A* targets as they appeared to be outside of any regulatory region. Similarly, with the plasmid-borne targets, we ignored RS26580 and the two *repA* targets as they appear to be connected with core plasmid functions, and the DNA site for CRP in the *pet* regulatory region has been already exhaustively characterised (13). Figure 2 shows the bandshift experiment with the remaining 8 targets. The data show a strong CRP interaction with 6 targets, but not with 3975 or *virK.* Inspection of the chromosomal context of each of the strong binding sites suggested that the EAEC 042 *kpsM*, *mchA*, 0536 and 0225 targets were most likely to be associated with a promoter-active regulatory region. Hence DNA fragments carrying each region were cloned into plasmid pRW50, a low copy number *lac* expression vector, each recombinant plasmid was used to transform the *Escherichia coli* K-12 Δ*lac* strains, M182 and M182Δ*crp*, and β-galactosidase activities were measured. These activities are taken as a measure of promoter activity and comparison of levels in the *crp*^+^ and Δ*crp* strains give an indication of any CRP-dependent regulation. As a control, an assay with pRW50 carrying the semi-synthetic CC(-41.5) promoter, whose activity is tightly coupled to CRP (32), is included. The data, illustrated in Figure 3, show that the p*kpsM*_201, p*mchA*, p0536 fragments carry at least one promoter. The observed activity with *mchA* is clearly CRP-dependent, whilst, with the p*kpsM*_201 and p0536 fragments, activity is higher in the Δ*crp* strain, suggesting repression by CRP. These effects were investigated further and our experiments are described in the next sections. Whilst there is no evidence for any promoter activity associated with the 0225 target, we were concerned that CRP might be regulating expression of the divergent gene, 0224 and so the cloned fragment was inverted in pRW50. However, measured expression levels were very low (data not shown), and we conclude that the intergenic region between 0224 and 0225 may be bereft of promoter activity (at least in our experimental conditions).

**Fig. 2.**
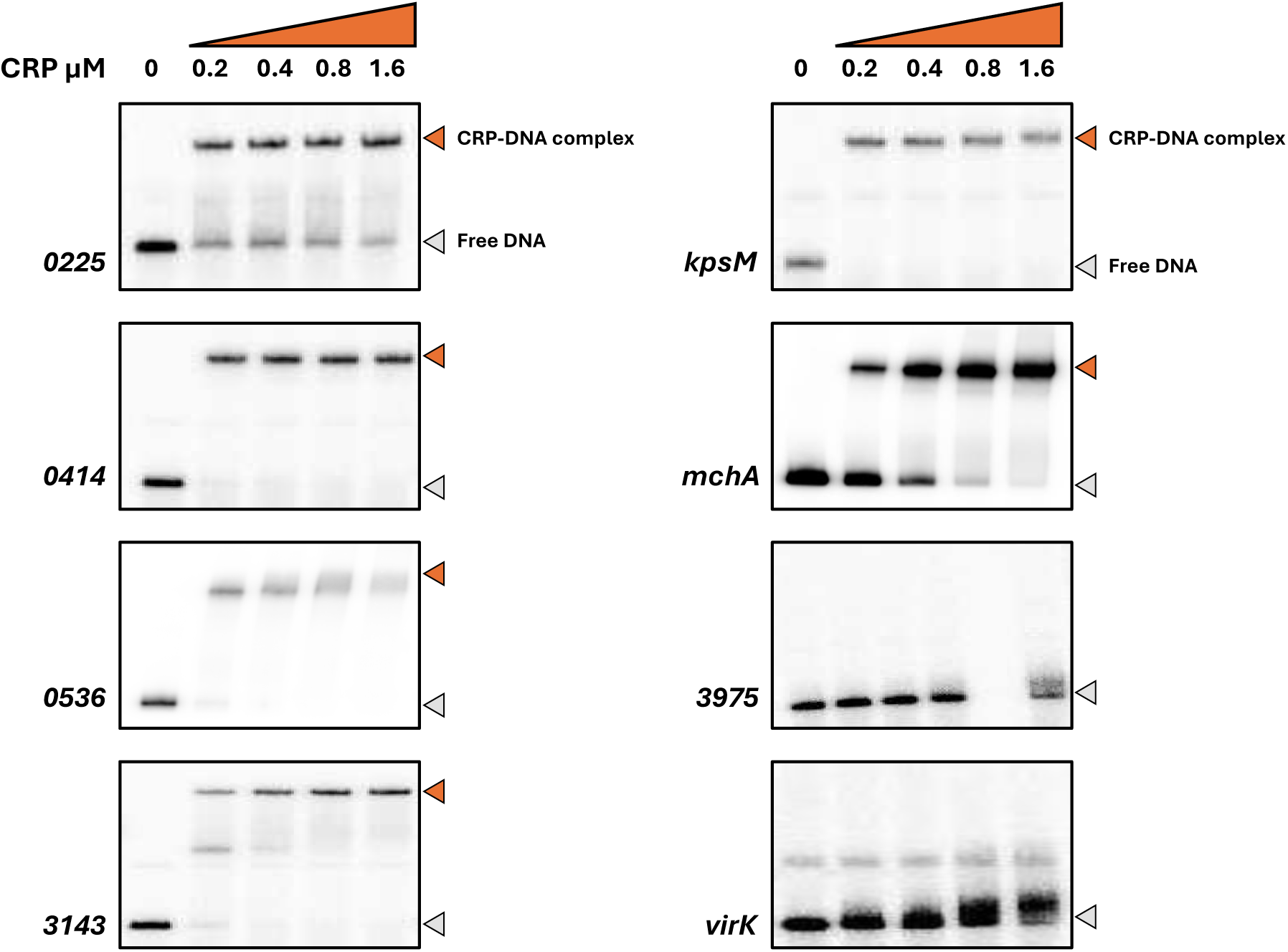
Electrophoretic mobility shift assays of CRP binding to EAEC 042 targets. Eight 042 targets, each approximately 200 bp in length, were excised from pSR as *Hind*III/*Eco*RI or *Hind*III/*Aat*III fragments and radiolabelled with γ32-P ATP. These were incubated, in the presence of cAMP, with increasing concentrations of CRP (0-1.6 µM) and then separated by electrophoresis on a 6% polyacrylamide gel. In each case the band in the first lane shows the mobility of free DNA, and is indicated by a grey arrow. In subsequent lanes, a reduced mobility band indicates a DNA fragment complexed with CRP, marked with an orange arrow.

**Fig. 3.**
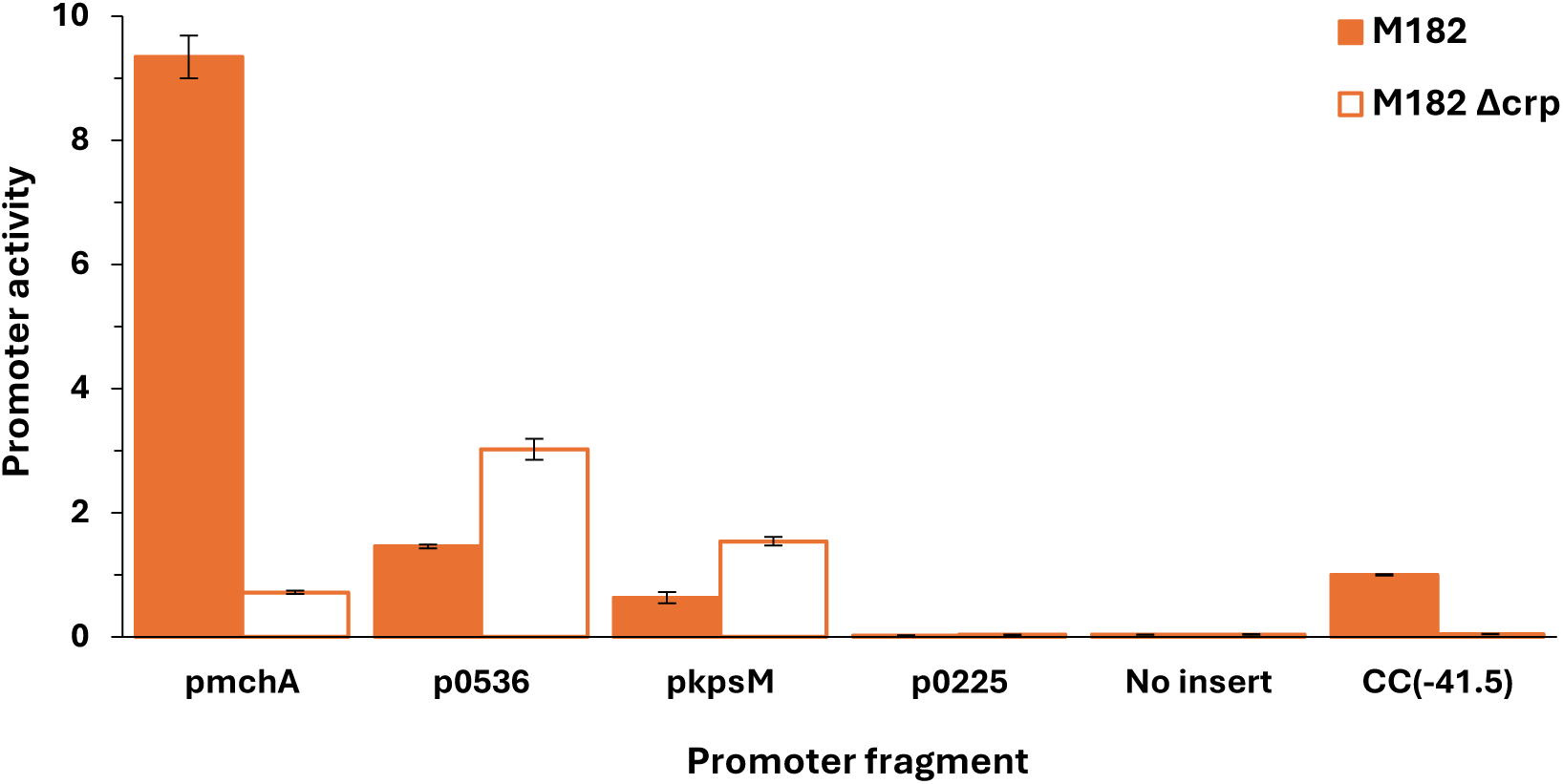
CRP regulation at EAEC 042 promoters. The figure shows β-galactosidase activity measured in *E. coli* K12 Δ*lac* M182 and M182 Δ*crp* strains carrying a *lacZ* reporter plasmid pRW50 containing a target promoter region: p*mchA*; p0536; p*kpsM*_201; or p0225. Activities are expressed as fold activity compared to the activity of pRW50 carrying the promoter, CC(-41.5) measured in M182 *crp*^+^ cells. pRW50 with no insert was included as a negative control. Data is a representative of duplicate independent experiments, averaged from three biological replicates, with standard deviations shown.

**Table 1:**
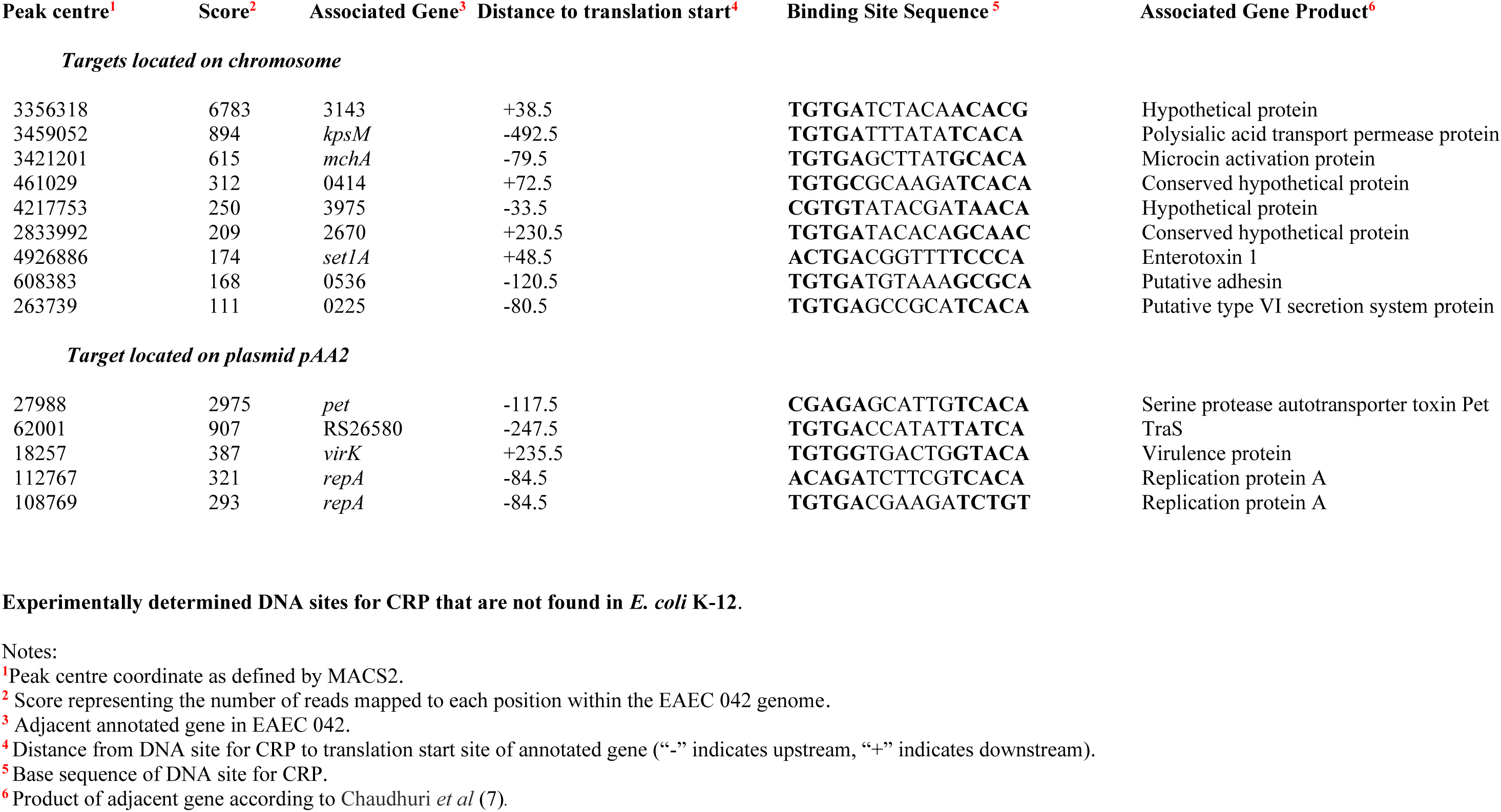
Uncommon targets for CRP in EAEC strain 042.

### CRP-dependent activation at the mchA promoter

The EAEC 042 *mchA* gene encodes an enzyme that transfers a glucose residue to microcin H47, a well-characterised bacterial antibiotic (33–36). The significance of the glucose residue is that it serves as a bridge to link a siderophore molecule, thereby creating a siderophore-toxin which can play an important role in bacterial colonisation and, hence, virulence (37). Data in Figure 3 suggest that *mchA* expression may be due to a strong promoter that is dependent on CRP. Recall that most bacterial promoters carry crucial sequence elements located ∼10, 14 and 35 bases upstream of the transcript start site (known as the -10, extended -10 and -35 elements, respectively), and that one or more of these is usually defective at promoters dependent on an activator such as CRP (38,39). Moreover CRP-activated promoters fall into two major classes: at Class I promoters the DNA site for CRP is located over 15 bases upstream from the promoter -35 element whilst, for Class II promoters, it overlaps the -35 element (40). Inspection of the base sequence adjacent the proposed DNA site for CRP at the *mchA* regulatory region suggested the existence of a Class II CRP-dependent promoter (Figure 4a). On this basis we created the 37G and 14A mutations that would be expected to inactivate the DNA site for CRP and the extended - 10 element respectively. DNA fragments carrying these base substitutions were cloned into pRW50: data in Figure 4b show that CRP-dependent activation is completely stopped by either mutation. Furthermore, an 11G mutation within the predicted -10 hexamer reduced activity and an 11A mutation, to make the base at the -11 position consensus, increased activity (Figure 4b). To confirm the existence of a Class II CRP-activated promoter, we complemented the M182Δ*crp* strain with plasmid-borne *crp* carrying substitutions in either or both of the major activating regions (known as AR1 and AR2). Data in Figure 4c show that both AR1 and AR2 are needed for optimal complementation, with AR2 making the larger contribution to activation. Since AR2 is known to play little or no role at Class I CRP-activated promoters (40), taken together, the evidence suggests that EAEC 042 *mchA* expression depends on a single Class II CRP-dependent promoter, and this was confirmed by an *in vitro* transcription assay (Figure 4d).

**Fig. 4.**
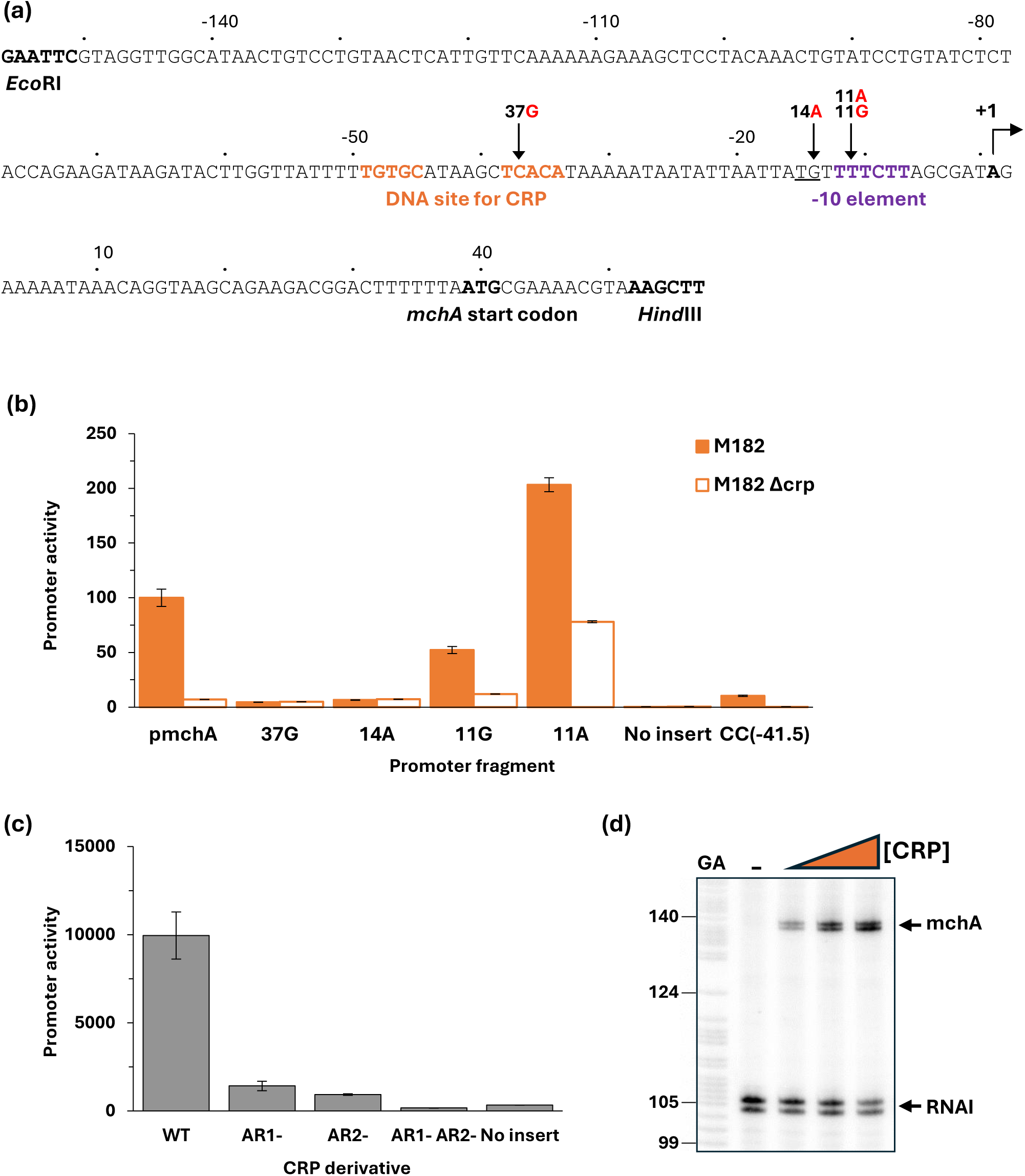
CRP-dependent activation of the EAEC 042 *mchA* promoter. (a) Sequence of the p*mchA* promoter fragment, amplified to incorporate *Eco*RI and *Hind*III restriction sites. Numbering is relative to the predicted transcript start site noted by +1. Locations of the -10 element and CRP binding site are shown in bold purple and orange respectively; the extended -10 is underlined; and substitutions labelled accordingly, for example 11G indicates the base at position -11 is changed to a G. (b) This panel shows mutational analysis of the *mchA* promoter. β-galactosidase activity was measured in *E. coli* K12 Δ*lac* strains M182 and M182 Δ*crp* carrying a reporter plasmid pRW50 containing either the starting *mchA* promoter (p*mchA*) cloned upstream of *lacZ* or the *mchA* promoter with a substitution in the CRP binding site (37G), in the extended -10 (14A) or the -10 element (11G or 11A). Promoter activities are expressed as a percentage relative to the starting *mchA* promoter activity in M182 *crp*^+^ cells. pRW50 with no insert was included as a negative control. Data is from duplicate independent experiments, averaged from three biological replicates, with standard deviations shown. (c) Complementation with CRP derivatives demonstrated Class II activation of the *mchA* promoter. β-galactosidase activity was measured in M182 Δ*crp* cells containing the wild type *mchA* promoter *lacZ* fusion (in pRW50) with derivatives of CRP expressed from pDCRP: wild type CRP (WT); CRP HL159 mutated in AR1 (AR1-); KE101 in AR2 (AR2-); and HL159 KE101 in both AR1 and AR2 (AR1- AR2-). pD with no CRP insert was included as a negative control. β-Galactosidase activity is expressed as Miller Units (nmol of ONPG hydrolysed min^-1^ mg^-1^ dry cell mass). Data is from duplicate independent experiments, averaged from three biological replicates, with standard deviations shown. (d) A CRP-dependent transcript was detected by multi round *in vitro* transcription from the *mchA* promoter, cloned in pSR plasmid, with *E. coli* holo-RNA polymerase (50 nM). The 110 nt RNAI transcript is an internal positive control. Both the *mchA* and RNAI transcripts are indicated by arrows. A Maxim-Gilbert GA sequencing reaction was used to calibrate the transcripts. CRP concentrations: lane 1 (0 µM), lane 2 (25 nM), and lane 3 (100 nM).

### CRP-dependent repression at the 0536 promoter

The EAEC 042 0536 gene appears likely to encode an adhesin (7), data in Figure 3 suggest that the 0536 gene regulatory region contains one or more promoters, and that CRP could down-regulate expression from them. Inspection of the base sequence upstream of the 0536 translation start suggested 4 possible promoter -10 elements. Hence, starting with the cloned p0536 regulatory region fragment, we constructed single base changes that would be expected to inactivate each of the putative -10 elements denoted 11G, 1G, 17G and 25G (Figure 5a). Assays of the promoter activity of each mutant fragment showed that the 11G mutation alone caused complete loss of activity (Figure 5b), suggesting that the 0536 regulatory region contains a single major promoter. This was corroborated by a permanganate footprint and an *in vitro* transcription assay (Figure 5c & 5d). The transcription assay shows an RNA that initiates 7 base pairs downstream from the suggested 5’-TAGATT-3’ -10 element, and the permanganate shows the expected open complex unwinding starting at the upstream end of the -10 element. Both the footprint and the transcript are suppressed by the inclusion of CRP in the assay (Figure 5c & 5d). Taken together, the data argue that the EAEC 042 0536 gene regulatory region contains a single promoter whose activity is down-regulated by CRP. Inspection of the promoter base sequence shows that the most likely -35 element is 5’-TTTACA-3’, located 17 base pairs upstream of the -10 element, which corresponds to the optimal spacing. This is sandwiched between the two flanking 5-base pair elements that comprise the DNA site for CRP and this suggests a simple mechanism for the repressive action of CRP. To confirm this, the 38T 36A double base substitution was constructed in the cloned p0536 regulatory region fragment (Figure 5a). These substitutions, which create a consensus upstream 5-base pair element in the DNA site for CRP, result in stronger repression of the 0536 promoter by CRP (Figure 5e).

**Fig. 5.**
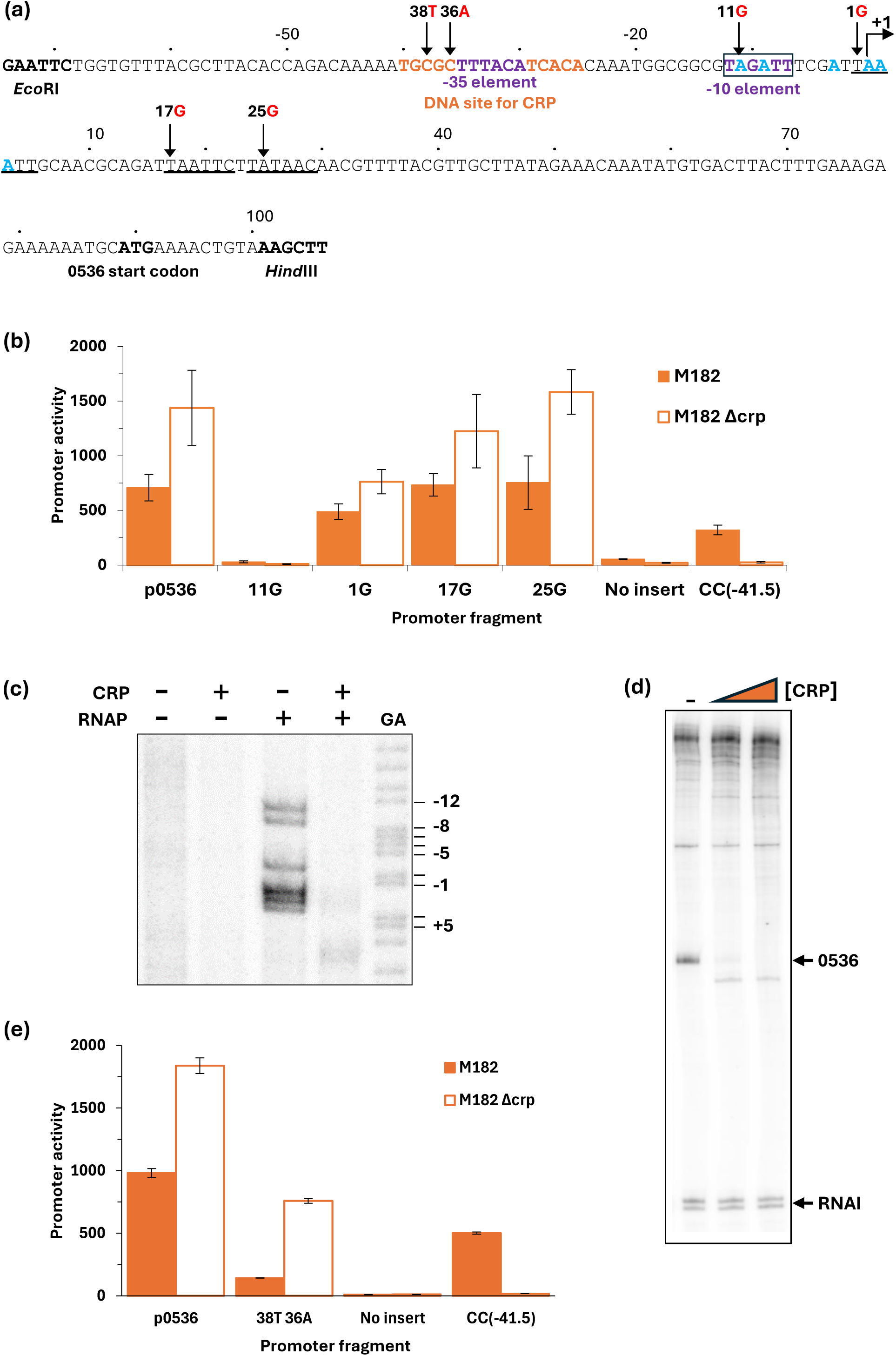
CRP-dependent repression of the EAEC 042 0536 promoter. (a) Sequence of the p0536*_*100 promoter fragment, amplified to incorporate *Eco*RI and *Hind*III restriction sites. Numbering is relative to the predicted transcript start site denoted by +1. Locations of the -10 element and predicted -35 element are shown in purple (with a box around the -10), the CRP binding site is shown in orange. Other potential -10 elements are underlined; and substitutions labelled accordingly, for example 11G indicates the base at position -11 is changed to a G. (b) Mutational analysis was used to locate the 0536 promoter -10 element. β-galactosidase activity was measured in *E. coli* K12 Δ*lac* strains M182 and M182 Δ*crp* carrying a reporter plasmid pRW50 containing either the wild type 0536 promoter (p0536_100) cloned upstream of *lacZ* or the 0536 promoter with a mutation in one of four possible -10 elements: 11G (p0536_105); 1G (p0536_106); +17G (p0536_103); or +25G (p0536_104). pRW50 with the CC(-41.5) promoter was included as a positive control. β-galactosidase activity is expressed as Miller Units (nmol of ONPG hydrolysed min^-1^ mg^-1^ dry cell mass). Data is from duplicate independent experiments, averaged from three biological replicates, with standard deviations shown. (c) Unwinding of the open complex at the -10 element was confirmed with potassium permanganate footprinting of an end-labelled *Aat*II/*Hin*dIII 0536 promoter fragment excised from pSR incubated, in the presence of cAMP (200 µM), with (+) and without (-) CRP (100 nM) and RNA polymerase (50 nM). A Maxim-Gilbert GA sequencing reaction was used to calibrate the gel. The locations of cleavage sites are shown in blue in panel (a). (d) A transcript, produced in the absence of CRP, was detected by multi round *in vitro* transcription from the p0536_100 promoter, cloned in pSR plasmid, with *E. coli* holo-RNA polymerase (50 nM). The 110 nt RNAI transcript is an internal positive control. Both the 0536 and RNAI transcripts are indicated by black arrows. A Maxim-Gilbert GA sequencing reaction was used to calibrate the transcripts. CRP concentrations: lane 1 (0 µM), lane 2 (25 nM), and lane 3 (100 nM). (e) Substitutions to create a consensus CRP binding site resulted in stronger CRP repression of the p0536 promoter. β-galactosidase activity was measured in *E. coli* K12 Δ*lac* M182 and M182 Δ*crp* strains carrying a reporter plasmid pRW50 containing either the wild type 0536 promoter (p0536_100) cloned upstream of *lacZ* or the 0536 promoter with double base substitution 38T 36A (p0536_107) in the CRP binding site to make it consensus. pRW50 with the CC(-41.5) promoter was included as a positive control. β-galactosidase activity is expressed as Miller Units (nmol of ONPG hydrolysed min^-1^ mg^-1^ dry cell mass). Data is from duplicate independent experiments, averaged from three biological replicates, with standard deviations shown.

### CRP-dependent repression at the kpsM regulatory region

The *E. coli kps* gene products enable the production of a form of capsule known as K antigen (41–43). Although absent in MG1655, the *kps* genes are found in many *E. coli* pathovars, and their expression has been studied in some detail (44,45). The *kpsM* gene is unusual in that its promoter is located ∼750 base pairs upstream from its start codon. Inspection of the regulatory region showed that the DNA site for CRP is ∼250 base pairs downstream from this promoter (Figure 6a, denoted here as P1), which is consistent with the observed repression by CRP (Figure 3). However, we noted that the two consensus halves of this DNA site for CRP flank a near perfect promoter -10 sequence (5’-TATAAA-3’), suggesting the possible existence of a second promoter (denoted here as P2). To investigate this, we cloned a 440 bp DNA fragment covering both the CRP binding site and the previously identified promoter region (p*kpsM*_440, Figure 6a). We made single base changes in the p*kpsM*_440 fragment, in which we corrupted the upstream promoter -10 element (P1^-^), the downstream -10 element (P2^-^), or both (P1^-^P2^-^), and we measured the net promoter activity in our *crp*^+^ and Δ*crp* strains as in Figure 3. The data, illustrated in Figure 6b show that *kpsM* promoter activity is largely suppressed by the two base changes together. Crucially, the fragment carrying the P1^-^ mutation shows significant activity in the Δ*crp* genetic background, but this activity is greatly suppressed in the *crp*^+^ strain. These data are consistent with a model in which the activity of the dominant upstream promoter (P1) is moderated by CRP, but a second weaker downstream promoter (P2) is sharply repressed by CRP because its -10 hexamer element is embedded in the DNA site for CRP. This was corroborated by an *in vitro* transcription assay (Figure 6c).

**Fig. 6.**
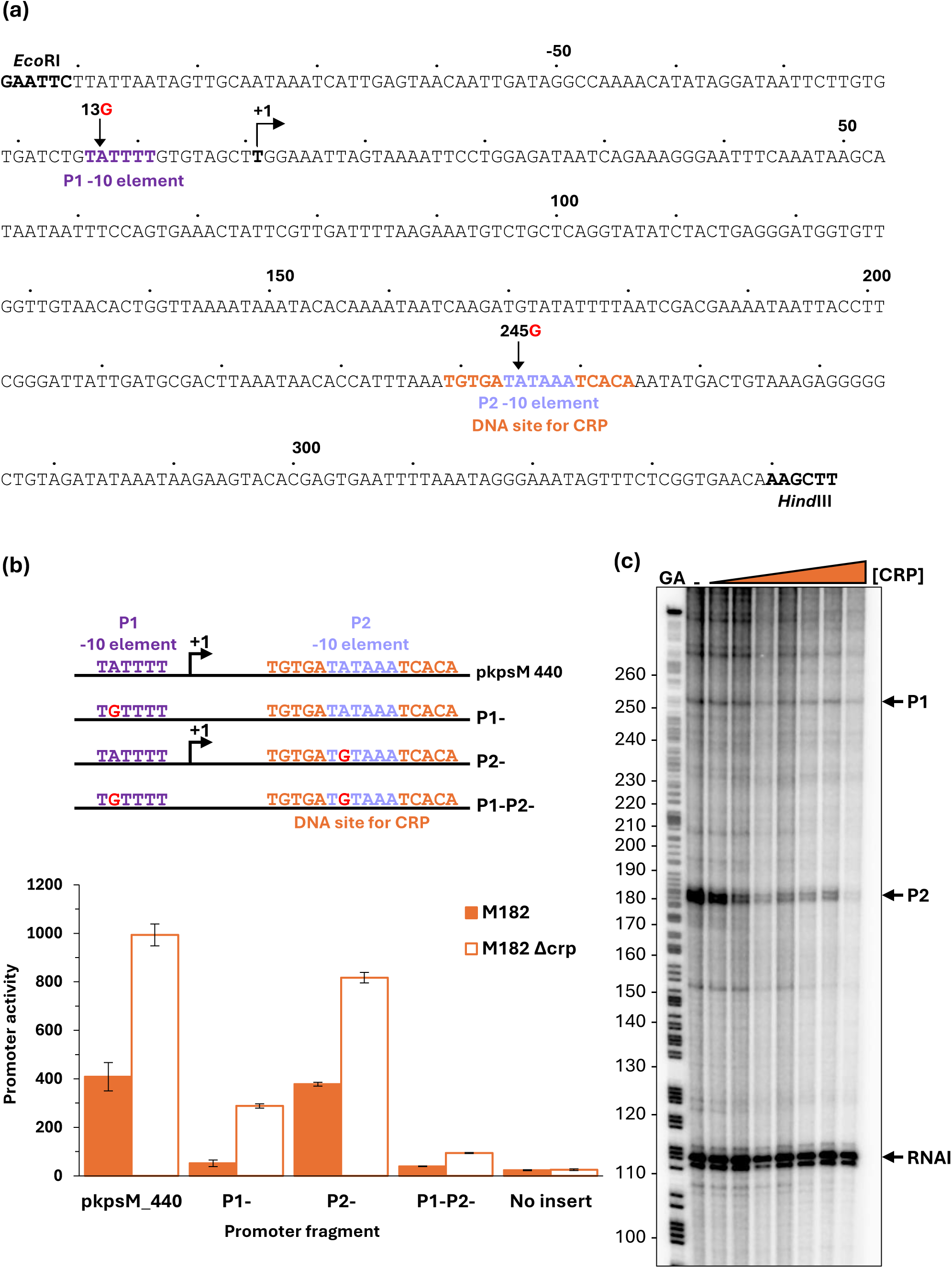
CRP-dependent repression of the EAEC 042 *kpsM* promoter. (a) Sequence of the p*kpsM*_440 promoter fragment, amplified to incorporate *Eco*RI and *Hind*III restriction sites. Numbering is relative to the predicted transcript start site from the P1 promoter noted by +1. Locations of the P1 and P2 promoter -10 elements are coloured purple and blue respectively; the CRP binding site is shown in orange; and substitutions labelled accordingly, 13G indicates the base at position -13 is changed to a G. (b) Mutational analysis confirmed the presence of two *kpsM* promoters. β-galactosidase activity was measured in *E. coli* K12 Δ*lac* M182 and M182 Δ*crp* strains carrying a reporter plasmid pRW224 containing either the starting *kpsM* fragment (p*kpsM*_440) cloned upstream of *lacZ*, or the p*kpsM* fragment with a 13G mutation in the P1 -10 element (P1-), or 245G in P2 (P2-), or 13G 245G in both (P1-P2-). pRW224 with no insert was included as a negative control. β-galactosidase activity is expressed as Miller Units (nmol of ONPG hydrolysed min^-1^ mg^-1^ dry cell mass). Data is from duplicate independent experiments, averaged from three biological replicates, with standard deviations shown. (c) *kpsM* P1 and P2 transcripts were detected by multi round *in vitro* transcription from the p*kpsM*_440 DNA construct, cloned into the pSR plasmid, with *E. coli* holo-RNA polymerase (123 nM). The 110 nt RNAI transcript is an internal positive control. *kpsM* P1, P2, and RNAI transcripts are indicated by black arrows. A Maxim-Gilbert GA sequencing reaction was used to calibrate the transcripts. CRP concentrations: lane 1 (0 µM), lane 2 (0.1 μM), lane 3 (0.2 μM), lane 4 (0.4 μM), lane 5 (0.6 μM), lane 6 (0.8 μM), lane 7 (1 μM), and lane 8 (2 μM).

## CONCLUSIONS

The starting point of this work was a ChIP-seq experiment, designed to identify the different DNA sites for CRP on the chromosome and virulence plasmid of enteroaggregative *E. coli* strain 042. Arguably, CRP is the most widely studied bacterial transcription factor. It was discovered originally as the essential activator of *E. coli* lactose operon expression (46) and its activity was found to be triggered by the binding of cyclic AMP (47–50). Subsequently it was shown to regulate 100s of *E. coli* transcription units and its biological role appears to be to reprogramme gene transcription in response to nutritional stresses (30,51–53). Our research, reported here, was prompted by the observation that transcription of two key EAEC 042 virulence genes (*pet* and *pic*) directly depends on CRP (13,18), rather than AggR, which is regarded as the master regulator of virulence in EAEC (9) . We wanted to discover if the transcription of any other EAEC virulence genes was likewise activated by CRP. Our experiments showed that, as expected, CRP binds to hundreds of gene regulatory regions and that the vast majority of binding targets are common with the MG1655 *E. coli* reference strain. However, after triage, we found CRP-dependent regulation at three genes that are not found in MG1655. In one case, *mchA*, CRP is an activator, whilst in the other two cases, 0536 and *kpsM*, CRP downregulates gene expression. We propose that, whilst EAEC virulence is primarily controlled by the activity of AggR, CRP activity, which increases in response to stress (17), drives expression of a mucinase, the plasmid-encoded toxin and the microcin-siderophore, all of which are useful during the colonisation stage of infection, and, subsequently, at the exit stage of infection. In addition, CRP suppresses expression of an adhesin and production of capsule. By exploiting CRP, EAEC aligns the infection with nutrient availability. From a mechanistic point of view, the two CRP-suppressed promoters are remarkable. Most bacterial repressors act by blocking the access of RNA polymerase to the target promoter. It is significant that, at the EAEC 0536 and *kpsM* P2 promoters, evolution has driven the DNA site for CRP to include the promoter -35 and -10 hexamer elements respectively. Recall that, at most bacterial promoters, these are the two most important elements for promoter function (30), and yet this arrangement is uncommon at CRP-repressed promoters.

Although most textbooks consider the *E. coli* CRP protein to be a gene regulatory transcription factor, at many targets, its binding has little or no direct consequence for gene expression (54). This was first discovered following the application of chromatin immunoprecipitation that reports factor binding independent of any function, and this study with EAEC strain 042 tells the same story as the original chromatin immunoprecipitation study on *E. coli* K-12 (14,29). A plausible explanation for this is that the original function of ancestral CRP had little or nothing to do with transcription regulation but, rather was concerned with management of bacterial chromosomes (54). It has been argued that gene regulatory functions evolved later, and this may explain the variety of architectures found at different CRP-regulated promoters (54,55). Similar observations have been made with other bacterial transcription factors (56–58) and it is possible to argue that these apparently function-less bound factors can, given certain constraints, evolve to participate in functional gene regulatory regions. Given this flexibility, it is easy to imagine that incoming DNA segments, which encode genetic determinants that are ‘useful’ to a bacterial host, could become subject to regulation by CRP. Our data suggest that whilst CRP-dependent function may have resulted at some loci, there are still others where CRP binding has no functional outcome. However, we can discern how CRP imposes a layer of regulation on the expression of virulence determinants and this may, in future, prompt new strategies for interfering with virulence gene expression.

## ACKNOWLEDGEMENTS

This work was supported by the Biotechnology and Biological Sciences Research Council [BB/R017689/1 and BB/W00285X/1] and Princess Nourah bint Abdulrahman University Researchers Supporting Project number (PNURSP2025R898), Princess Nourah bint Abdulrahman University, Riyadh, Saudi Arabia. DCG is grateful for funding from a Wellcome Trust Investigator Award (212193/Z/18/Z).

