## Supplementary information for "Genome-wide mapping of cyclic AMP receptor protein binding in Enteroaggregative *Escherichia coli* reveals targeting of virulence-associated genes"

### Table S1: Strains, plasmids and oligonucleotides used

#### *Strains*

| **Strains** | **Characterisations** | **Source** |
|---|---|---|
| EAEC 042 | Wild type, prototype strain, $Sm^R$, $Tet^R$, $Cm^R$, Diarrhoeagenic in volunteers, expresses AAF/II, biofilm positive, harbours pAA2 plasmid. | (1) |
| *E. coli* M182 | Δ*(codB-lacI)3, galK16, galE15(GalS),* $\lambda^-$*, e14-, relA1, rpsL150(strR), spoT1.* | (2) |
| *E. coli* M182 **Δ***crp* | Δ*(codB-lacI)3, galK16, galE15(GalS),* $\lambda^-$*, e14-, relA1, rpsL150(strR), spoT1 and* Δ*crp*. | (3) |

#### *Plasmids*

| **Plasmids** | **Description** | **Source** |
|---|---|---|
| pRW50 | Low-copy- number *lacZ* expression vector, allows promoter fragments to be cloned using *Eco*RI and *Hin*dIII sites as fusions to *lacZ* transcription. Carries tetracycline resistance gene. | (4) |
| pRW224 | A pRW50 derivative which allows promoter fragments to be cloned using *Eco*RI and *Hin*dIII sites as fusions to *lacZ* transcription, carries tetracycline resistance gene. | (5) |
| pSR | Supercoiled small plasmid with 2.6 kb with *Eco*RI and *Hin*dIII sites for inserting the desired promoter fragment which will be located upstream of a λ*oop* terminator site. Carries ampicillin resistance gene. | (6) |
| pD | pBR322 derivative. Carries ampicillin resistance gene. | (7) |
| pDCRP | pD derivative carrying the *crp* gene. Carries ampicillin resistance gene. | (7) |
| pDCRP $AR1^-$ | A pDCRP derivative carrying a substitution in CRP activating region AR1 (HL159). | (7) |
| pDCRP $AR2^-$ | A pDCRP derivative carrying a substitution in CRP activating region AR2 (KE101). | (7) |
| pDCRP $AR1^-$ & $2^-$ | A pDCRP derivative carrying substitutions in CRP activating regions AR1 (HL159) and AR2 (KE101). | (7) |

#### *Oligonucleotide primers*

| Oligo name | Sequence (5`-3`)** | Use |
|---|---|---|
| *Primers used to sequence pRW50/pRW224 constructs* | | |
| pRW50 F | CCCTGCGGTGCCCCTCAAG | Primer for sequencing in the forward direction, binding upstream of the *Eco*RI site of pRW50/ pRW224. |
| pRW224 R | GGCGATTAAGTTGGGTAACGCCAGGG | Primer for sequencing in the reverse direction, binding downstream of *Hin*dIII site in pRW224. |
| pRW50 R | GCAGGTCGTTGAACTGAGCCTGAAATTCAG | Primer for sequencing in the reverse direction, binding downstream of *Hin*dIII site in pRW50. |
| *Primers used to generate fragments used for EMSA experiments* | | |
| EC042_0225 F | GGGGGGAATTCAACTCGAATAAAGAAAAGGGTGTG | Forward primer to amplify EC042_0225 fragment. |
| EC042_0225 R | GGGGGAAGCTTATGGGGTTGGCATTTATG | Reverse primer to amplify EC042_0225 fragment. |
| EC042_0414 F | GGGGGGAATTCATGTTGCAATCTTCTGCTGACAAAGC | Forward primer to amplify EC042_0414 fragment. |
| EC042_0414 R | GGGGGAAGCTTTTAACTTATAATTAAGAGAAAAAAC | Reverse primer to amplify EC042_0414 fragment. |
| p0536_100 F | GGGGGAATTCTGGTGTTTACGCTTACACCAGACA | Forward primer to amplify p0536 fragment and it’s derivatives. |
| p0536_100 R | GGGGAAGCTTTACAGTTTTCATGCATTTTTTCTC | Reverse primer to amplify p0536 fragment and it’s derivatives. |

| | | |
|---|---|---|
| EC042_3143 F | GGGGGGAATTCATGTCCGTTTGCGGACAAGCAATAG | Forward primer to amplify EC042_3143 fragment. |
| EC042_3143 R | GGGGGAAGCTTGGGAAACCGGTGTTTTGAAAACAGT | Reverse primer to amplify EC042_3143 fragment. |
| EC042_3975 F | GGGGGGAATTCAGATTCGGTTTTTCAGACCCCCATC | Forward primer to amplify EC042_3975 fragment. |
| EC042_3975 R | GGGGGAAGCTTATAAGAGTTATTCAAAATATTTTGT | Reverse primer to amplify EC042_3975 fragment. |
| p*kpsM*_201 F | GGGGGGAATTCTAATTACCTTCGGGATTATTGATG | Forward primer to amplify p*kpsM*_201 fragment. |
| p*kpsM*_201 R | GGGGGAAGCTTCTGTTTCAAACCAGAACGCACGCG | Reverse primer to amplify p*kpsM*_201 fragment. |
| p*mchA*_600 F: | GGAATTCCGCTGGTATTGCGGAAATCAG | Forward primer used to amplify a 619 bp “p*mchA*_600” used for generating the p*mchA* promoter construct. |
| p*mchA* R | GGGAAGCTTTACGTTTTCGCATTAAAAAAGTCCG | Reverse primer used to amplify a 619 bp “p*mchA* intermediate” used for generating the p*mchA* promoter construct. |
| p*mchA*_200 SDM F: | GTAGGTTGGCATAACTGTC | Forward primer used to generate the pmchA promoter construct by site directed mutagenesis (SDM) using the “p*mchA* intermediate” as template. |
| p*mchA*_200 SDM R: | GAATTCAGATCTAGCGTG | Reverse primer used to generate the pmchA promoter construct by (SDM) using the “p*mchA* intermediate” template. |
| p*mchA*_200 F: | GGAATTCGTAGGTTGGCATAACTGTCCTG | Forward primer used to amplify p*mchA* promoter construct, used with p*mchA* R to generate 213 bp fragment. |
| *virK* F | GGGGGGAATTCTCATGTTTTCCGGCAATTGAGATAC | Forward primer to amplify *virK* fragment. |

| | | |
|---|---|---|
| *virK* R | GGGGGAAGCTTTTTCGGTACTCAGAGCGTTTTTTAC | Reverse primer to amplify *virK* fragment. |
| *Primers used to generate* mchA *promoter derivatives* | | |
| p*mchA* 37G F | TGCATAAGCT**g**ACATAAAAATAATATTAATTATG | Forward primer to introduce 37G mutation in the p*mchA* promoter construct by SDM. |
| p*mchA* 37G R | CAAAAATAACCAAGTATCTTATC | Reverse primer used to introduce 37G mutation into the p*mchA* promoter construct by SDM. |
| p*mchA* p11G F | TAATTATGTT**g**TCTTAGCGATAGAAAAATAAAC | Forward primer to introduce 11G mutation in the p*mchA* promoter construct by SDM |
| p*mchA* p11G R | ATATTATTTTTATGTGAGCTTATGC | Reverse primer used to introduce 11G mutation into the p*mchA* promoter construct by SDM. |
| p*mchA* p11A F | TAATTATGTT**a**TCTTAGCGATAGAAAAATAAAC | Forward primer to introduce 11A mutation in the p*mchA* promoter construct by SDM |
| p*mchA* p11A R | ATATTATTTTTATGTGAGCTTATG | Reverse primer used to introduce 11A mutation into the p*mchA* promoter construct by SDM |
| p*mchA* SOE 14A left arm F: | GGAATTCGTAGGTTGGCATAACTGTCCTGTAACTCATTGTTCAAAAAAGAAAGCTCCTACAAACTGTATCCTGTATCTC | Forward primer used to generate left arm of p*mchA* 14A promoter variant. |
| p*mchA* SOE 14A left arm R: | GCTTATGCACAAAAATAACCAAGTATCTTATCTTCTGGTAGAGATACAGGATACAGTTTGTAGGAG | Reverse primer used to generate left arm of p*mchA* 14A promoter variant. |
| p*mchA* SOE 14A right arm F: | GATACTTGGTTATTTTTGTGCATAAGCTCACATAAAAATAATATTAATTAT**a**TTTTCTTAGCGATAGAAAAATAAACAGGTAAGCAG | Forward primer used to generate right arm of p*mchA* 14A promoter variant. |

| | | |
|---|---|---|
| p*mchA* SOE 14A right arm R | GGG<u>AAGCTT</u>TACGTTTTCGCATTAAAAAAGTCCGTCTTCTGCTTACCTGTTTATTTTTCTATCGCTAAG | Reverse primer used to generate right arm of p*mchA* 14A promoter variant. |
| *Primers used to generate* 0536 *promoter derivatives* | | |
| p0536_103 17G F | GCAACGCAGAT**g**AATTCTTATAAC | Forward primer to introduce 17G mutation in the p0536 promoter construct by SDM. |
| p0536_103 17G R | AATTTAATCGAAATCTACGC | Reverse primer to introduce the 17G mutation in the p0536 promoter construct by SDM |
| p0536_104 25G F | ATTAATTCTT**g**TAACAACGTTTTACG | Forward primer to introduce 25G mutation in the p0536 promoter construct by SDM. |
| p0536_104 25G R | CTGCGTTGCAATTTAATC | Reverse primer to introduce 25G mutation in the p0536 promoter construct by SDM. |
| p0536_105 11G F | ATGGCGGCGT**g**GATTTCGATT | Forward primer to introduce 11G mutation in the p0536 promoter construct by SDM. |
| p0536_105 R 11G | TTGTGTGATGTAAAGCGC | Reverse primer to introduce 11G mutation in the p0536 promoter construct by SDM |
| p0536_106 1G F | AGATTTCGAT**g**AAATTGCAACGCAGATTAATTC | Forward primer to introduce 1G mutation in the p0536 promoter construct by SDM. |
| p0536_106 1G R | ACGCCGCCATTTGTGTGA | Reverse primer to introduce 1G mutation in the p0536 promoter construct by SDM |
| p0536_107 38T 36A F | GATC<u>GAATTC</u>TGGTGTTTACGCTTACACCAGACAAAAATG**t**G**a**TTTACATCACACAAATGGCGGCGTAGATTTCGATTAAATTGCAACGCAGATTAATTCTTATAACAACG | Forward primer to introduce 38T 36A mutations in the p0536 promoter construct by PCR. |

| | | |
|---|---|---|
| p0536_107 38T 36A R | GATCAAGCTTTACAGTTTTCATGCATTTTTTCTCTTTCAAAGTAAGTCACATATTTGTTTCTATAAGCAACGTAAAACGTTGTTATAAGAATTAATCTGCG | Reverse primer to introduce 38T 36A mutations in the p0536 promoter construct by PCR. |
| *Primers used to generate* kpsM *promoter derivatives* | | |
| p*kpsM*_440 F | GGGGGGAATTCTTATTAATAGTTGCAATAAATCA | Forward primer to amplify p*kpsM*_440 and P2- fragments. |
| p*kpsM*_440 R | GGGGGAAGCTTTGTTCACCGAGAAACTATTT | Reverse primer to amplify p*kpsM*_440 and P1- fragments. |
| p*kpsM* P1- F | GGGGGGAATTCTTATTAATAGTTGCAATAAATCATTGAGTAACAATTGATAGGCCAAAACATATAGGATAATTCTTGTGTGATCTGT**g**TTTTGTGTAGC | Forward primer to introduce 13G mutation in p*kpsM*_440 P1- and P1-P2- fragments. |
| p*kpsM* P2- R | GGGGGAAGCTTTGTTCACCGAGAAACTATTTCCCTATTTAAAATTCACTCGTGTACTTCTTATTTATATCTACAGCCCCCTCTTTACAGTCATATTTGTGATTTA**c**ATCACATTTA | Reverse primer to introduce 245G mutation p*kpsM*_440 P2- and P1-P2- fragments. |

** Underlined bases represent the site for restriction enzymes *Eco*RI or *Hind*III. Base changes introduced by primers are in lower case and in bold.

### Table S2: List of all CRP sites in *Escherichia coli* 042 as determined by ChIP-seq

| Peak centre [a] | Score [b] | Annotated gene [c] | *E. coli* homologues [d] | Position relative to TSS [e] | Motif identified [f] | P-value [g] | Regulated by CRP? [h] | Gene product [i] |
|---|---|---|---|---|---|---|---|---|
| *Targets on the 042 chromosome* | | | | | | | | |
| 4488945 | 19348 | EC042_4210 | *udp* | -81.5 | GGTGATGGGTATCACG | 4.54E-05 | Y | Uridine phosphorylase |
| 4149109 | 17137 | EC042_3905 | *mtlA* | -224.5 | TGTGATTCAGATCACA | 4.16E-08 | Y | Mannitol-specific PTS system EIICBA component |
| 48719 | 16242 | *fixA* | *fixA* | -204.5 | GGTGATCTATAAAACA | 1.59E-03 | Y | Probable electron transfer flavoprotein subunit of oxidoreductase |
| 2531800 | 12889 | EC042_2405 | *mtlD* | -101.5 | ATTGATCGCCCTCACA | 1.14E-04 | Y | Mannitol dehydrogenase family protein |
| 3838726 | 12183 | *tsgA* | *ppiA* | -129.5 | CGTGATGAGAATCACT | 4.95E-05 | Y | Peptidyl-prolyl cis-trans isomerase |
| 2993024 | 8657 | *raiA* | *raiA* | -70.5 | TGAGATTTCCATCACA | 1.74E-05 | Y | Ribosome-associated inhibitor A |
| 2784492 | 7180 | *ptsH* | *ptsH* | -326.5 | TGTGGCCTGCTTCAAA | 4.95E-05 | Y | Phosphocarrier protein Hpr |
| 1250181 | 7061 | EC042_1182 | *ptsG* | -76.5 | TGTGATCCAGATCACA | 1.40E-09 | Y | PTS glucose transporter subunit IIBC |
| 3356318 | 6783 | EC042_3143 | Uncommon | 38.5 | TGTGATCTACAACACG | 1.50E-05 | - | Hypothetical protein |
| 3350600 | 5944 | *epd* | *epd* | -202.5 | TCTGACTCACATCACA | 9.63E-06 | Y | D-erythrose 4-phosphate dehydrogenase |
| 2529223 | 4498 | *fruB* | *fruB* | -167.5 | TTTGATGTGCTGCACA | 3.81E-05 | N | Multiphosphoryl transfer protein |
| 4560899 | 4155 | *fdhD* | *fdhD* | 39.5 | TGTGACAAATGTCACA | 1.64E-06 | N | Putative formate dehydrogenase accessory protein |
| 5218617 | 4089 | EC042_4877 | *deoC* | -119.5 | TTTGAACCAGATCGCA | 1.23E-05 | Y | Deoxyribose-phosphate aldolase |
| 2151336 | 4034 | *ftnB* | *flhD* | -40.5 | TTTGATTTATCTTGCA | 1.69E-03 | Y | Transcriptional activator FlhD |
| 2497609 | 3901 | *cdd* | *cdd* | -67.5 | TGAGATTCAGATCACA | 9.11E-06 | Y | Cytidine deaminase |
| 4954732 | 3900 | *fxsA* | *aspA/fxsA* | -145.5 | CGTAATCTGGATCACT | 2.93E-04 | Y | Aspartate ammonia-lyase/ membrane protein FxsA |
| 4590079 | 3873 | *glpF* | *glpF* | 244.5 | TGCGCTTGTCGAAACA | 4.23E-03 | Y | Glycerol uptake facilitator protein |

| | | | | | | | | |
|---|---|---|---|---|---|---|---|---|
| 3377662 | 3657 | *ansB* | *ansB* | -114.5 | TTAGAGGCAGGTAACA | 1.85E-03 | Y | L-asparaginase 2 |
| 2535535 | 3525 | *spr* | *mepS* | -119.5 | TGTGCGTTAGTCCACA | 4.95E-05 | N | Lipoprotein |
| 3190075 | 3516 | *sdaC* | *sdaC* | -168.5 | TGGGATCAAGATCACT | 7.25E-05 | N | Serine transporter |
| 3719403 | 3355 | *elbB* | *elbB* | -58.5 | AATGCTACGCATCACA | 1.50E-03 | N | Enhancing lycopene biosynthesis protein 2 (sigma cross-reacting protein 27A) |
| 2073958 | 3353 | EC042_1989 | *manX* | 125.5 | CTCGATCAGACGCACA | 5.52E-04 | Y | Mannose-specific PTS system EIIAB component |
| 5094768 | 3314 | EC042_4748 | In K-12 | -67.5 | TGCGACAATCATCACA | 5.12E-06 | - | PTS system EIIA component |
| 4398875 | 2967 | *rbsD* | *rbsD* | -91.5 | TTCGAGGTTGATCACA | 2.10E-05 | Y | Putative ribose transport (metabolism protein) |
| 2034392 | 2844 | EC042_1961 | *gapA* | -87.5 | TGTGACGAGCATCACG | 1.36E-06 | Y | Glyceraldehyde 3-phosphate dehydrogenase A |
| 3382409 | 2794 | *nupG* | *nupG* | 40.5 | TGTGAGGAAATTAACA | 1.18E-04 | Y | Nucleoside permease NupG |
| 2430107 | 2264 | EC042_2320A | *yegQ* | 834.5 | TTAGAAACCGATCACA | 4.49E-04 | N | tRNA 5-hydroxyuridine modification protein YegQ |
| 4682235 | 2044 | *purH* | *purH* | -352.5 | TGTGAATCACTTCACA | 3.28E-06 | N | Bifunctional purine biosynthesis protein |
| 2795558 | 1964 | *ucpA* | *ucpA* | -82.5 | TGCGGATCAGCTCACT | 4.98E-04 | N | SDR family oxidoreductase UcpA |
| 2773567 | 1953 | EC042_2614 | *yfeC* | -88.5 | AGTTATTCATGTCACG | 9.67E-04 | N | Putative DNA-binding transcriptional regulator |
| 494392 | 1853 | *tsx* | *tsx* | -151.5 | ATCGATTGCGTTCACG | 1.55E-03 | Y | Nucleoside-specific channel-forming protein Tsx |
| 2975867 | 1701 | EC042_2790 | *patZ* | -63.5 | TCTGGGTAGCATCACA | 2.27E-04 | Y | Protein lysine acetyltransferase |
| 1190486 | 1700 | *rne* | *ycdZ* | -241.5 | TGCAACCCGCAGCCCG | 5.37E-03 | Y | DUF1097 domain-containing protein |
| 719805 | 1585 | *rnk* | *rnk* | -129.5 | AGTGATTTGCGTCACA | 7.43E-06 | N | Nucleoside diphosphate kinase regulator |
| 4801045 | 1563 | *proP* | *proP* | -216.5 | TGTGAAGTTGATCACA | 2.69E-07 | Y | Proline (betaine transporter) |
| 3615779 | 1543 | *uxaC* | *uxaC* | -99.5 | CGTGAGATAGATCAAT | 3.04E-04 | Y | Glucuronate isomerase |
| 2877210 | 1487 | *xseA* | *xseA* | -7.5 | TTTGATCTCGCTCACA | 2.63E-06 | Y | Exodeoxyribonuclease VII large subunit |
| 1336252 | 1427 | EC042_1262 | *dhaR* | -94.5 | TGTGATCACGCCCGCA | 1.30E-05 | N | PTS-dependent dihydroxyacetone kinase operon regulator (sigma-54 dependent transcriptional regulator) |

| | | | | | | | | |
|---|---|---|---|---|---|---|---|---|
| 1866407 | 1300 | *malX* | *uidR* | -83.5 | TGTGATTTATGCCTCA | 1.81E-04 | N | *uid* operon repressor (TetR-family transcriptional regulator) |
| 459356 | 1136 | EC042_0411 | *yaiZ* | 71.5 | TGTTGTCTACGCCACA | 1.88E-04 | N | Putative membrane protein |
| 3951363 | 1108 | *rpoH* | *rpoH* | -107.5 | CGTGATTTTATCCACA | 9.67E-05 | Y | RNA polymerase sigma-32 factor RpoH |
| 3152883 | 1050 | EC042_2961 | In K-12 | -33.5 | TGTGTCGGCGGTCAAT | 1.21E-03 | - | Putative FAD-dependent oxidoreductase |
| 732988 | 1014 | *cspE* | *cspE* | -103.5 | CGCGACTTTTATCACT | 2.53E-04 | Y | Cold shock-like protein |
| 3293513 | 1002 | EC042_3089 | *ygfK* | -271.5 | AGTTCTCTTTATCATA | 9.61E-03 | N | Pyridine nucleotide-disulfide oxidoreductase |
| 2480548 | 911 | EC042_2359 | *btsS* | -95.5 | CGTGAGTACGATCACT | 8.56E-05 | N | Two-component system sensor kinase |
| 3459052 | 894 | *kpsM* | Uncommon | -492.5 | TGTGATTTATATCACA | 2.69E-07 | - | Polysialic acid transport permease protein |
| 736066 | 860 | *lipA* | *lipA* | -119.5 | CCTGAAAGCAGCCAAA | 6.96E-03 | N | Lipoyl synthase |
| 3012994 | 857 | EC042_2821 | In K-12 | -175.5 | TCCGCTCTGGCTCATA | 1.91E-03 | - | Conserved hypothetical protein |
| 3645715 | 857 | *garD* | *garD* | -177.5 | TGCGCGCTAAAGCACA | 3.33E-05 | N | D-galactarate dehydratase pseudo |
| 5030357 | 827 | EC042_4698 | *ytfJ* | -109.5 | TGCGGTCAAGCGCACA | 3.81E-05 | N | YtfJ family protein |
| 5116358 | 827 | *msbB2* | Uncommon | -233.5 | TGTACCAGTCACCACA | 1.37E-03 | - | Virulence protein |
| 1381630 | 819 | EC042_1347 | *adhE* | -638.5 | TGTGATGAAAGCCTGT | 8.09E-03 | N | Aldehyde-alcohol dehydrogenase |
| 3878947 | 818 | *pckA* | *pckA* | -69.5 | TATGAGCCTTGTCGCG | 9.07E-04 | Y | Phosphoenolpyruvate carboxykinase [ATP] |
| 2627362 | 785 | *glpT* | *glpT* | -118.5 | TGTGAATTACCGCACA | 1.01E-05 | Y | Glycerol-3-phosphate transporter |
| 5192683 | 765 | EC042_4852 | *btsT* | -88.5 | TGTTGCCAGAGTTACG | 6.96E-03 | Y | Putative carbon starvation protein |
| 3638072 | 758 | *tdcA* | *tdcA* | -71.5 | TGTGCGACCACTCACA | 3.48E-05 | Y | Tdc operon transcriptional activator |
| 2769499 | 755 | *nupC* | *nupC* | -103.5 | TTTGAAGCTGGTCACA | 1.36E-05 | Y | Nucleoside permease NupC |
| 3747290 | 727 | EC042_3523 | *yhcN* | -49.5 | TGTGATATGGGTCACG | 8.60E-07 | N | Conserved hypothetical protein |
| 2622643 | 712 | EC042_2478 | *lysR/pseudo gene* | -71.5 | TGTGATGTACTTTGCA | 1.05E-04 | N | LysR-family transcriptional regulator (partial)/ MFS transporter Pseudo |
| 3335320 | 692 | *serA* | *serA* | -199.5 | CGTGACACATGTCACC | 8.56E-05 | Y | Phosphoglycerate dehydrogenase |
| 3231340 | 646 | EC042_3028 | In K-12 | -126.5 | CGTGACCCAGGTCACA | 1.67E-07 | - | Hypothetical protein |
| 1177277 | 639 | EC042_1101 | *putA/putP* | 6.5 | TGTGTGCTCGATCTCA | 2.44E-04 | Y | Bifunctional protein PutA/sodium proline |

| | | | | | | | | |
|---|---|---|---|---|---|---|---|---|
| | | | | | | | | symporter (proline permease) PutP |
| 1827122 | 631 | *uidA* | *mlc* | -170.5 | CGTGATATAGATCGCA | 2.34E-06 | Y | Protein Mlc (making large colonies protein) |
| 4741151 | 625 | EC042_4418 | *pspG* | 255.5 | TGTGCGGATGATCACA | 3.89E-06 | N | Putative membrane protein |
| 3421201 | 615 | *mchA* | Uncommon | -79.5 | TGTGAGCTTATGCACA | 1.30E-05 | - | Microcin activation protein |
| 3931572 | 607 | *gntK* | *gntK* | -121.5 | TGTGAGCTACTTCAAA | 9.11E-06 | Y | Thermoresistant gluconokinase |
| 228313 | 583 | *gmhB* | *gmhB* | 673.5 | TGTGAATCACTTCACA | 3.28E-06 | N | D-**g**lycero-β-D-manno-heptose-1,7-bisphosphate 7-phosphatase |
| 3832206 | 568 | EC042_3619 | In K-12 | 96.5 | TGCAAAGGACGTCACA | 1.88E-04 | - | Hypothetical protein |
| 4724645 | 552 | *malE* | *malE* | -121.5 | TGTGATCTCTGTTACA | 3.18E-05 | Y | Maltose transport system substrate-binding protein MalE |
| 2416645 | 544 | *mdtA* | *mdtA* | -432.5 | TGCTGGCAGGATCGCA | 6.31E-04 | N | Multidrug resistance protein |
| 1261443 | 508 | *minD* | *bhsA* | -358.5 | TGTGACCGGCGTTGTA | 2.08E-03 | Y | Putative exported protein |
| 1314710 | 502 | *dhaR* | *dadA* | -22.5 | TGCGAGCCGGAACACC | 3.77E-04 | Y | D-amino acid dehydrogenase small subunit |
| 2506198 | 500 | *mglB* | *mglB* | -259.5 | TGTGAAATCACTCACA | 7.43E-06 | Y | Galactose/glucose ABC transporter substrate-binding protein MglB |
| 2205763 | 485 | EC042_2120 | *yedR* | -46.5 | TGTGATAAAGGTCACA | 1.95E-07 | N | Putative membrane protein |
| 3771532 | 442 | *dusB* | *dusB* | -286.5 | TGCGAGCGATGTCACA | 2.78E-06 | Y | tRNA-dihydrouridine synthase B |
| 5027585 | 433 | *cpdB* | *cpdB* | -74.5 | AGTGAAGAATGCCACA | 2.53E-04 | Y | 2',3'-cyclic-nucleotide 2'-phosphodiesterase |
| 3101613 | 429 | *ascG* | *ascF* | -93.5 | GGTGACCGGTTTCACA | 2.91E-05 | Y | Arbutin-, cellobiose-, and salicin-specific PTS system EIIBC component |
| 4388273 | 416 | *atpI* | *atpI* | -252.5 | CGTGCTTCAGATCACA | 4.09E-06 | N | ATP synthase protein I |
| 467174 | 392 | *aroM* | *aroM* | -208.5 | AGGGATCTGCGTCACA | 8.56E-05 | N | AroM family protein |
| 5020611 | 392 | *cycA* | *cycA* | -187.5 | TGTGAGCTGTTTCGCG | 2.78E-05 | N | D-serine, D-alanine, glycine transporter |
| 5212600 | 384 | *osmY* | *osmY* | -36.5 | TGTTGCCAGGCTCAAA | 7.96E-04 | Y | Osmotically inducible protein Y |
| 3676811 | 377 | *deaD* | *deaD* | -22.5 | TGTGAACCGGCTCAAA | 1.12E-05 | N | ATP-dependent RNA helicase (DEAD-box protein) |
| 1711834 | 361 | EC042_1653 | *aslA* | -44.5 | CGTGATTACGATCACA | 1.85E-06 | N | Putative sulfatase |

| | | | | | | | | |
|---|---|---|---|---|---|---|---|---|
| 4753197 | 338 | EC042_4429 | *yjcB* | -39.5 | TGTGAACTATATCACA | 3.14E-07 | N | Putative membrane protein |
| 3517716 | 329 | EC042_3298 | *yghB* | -75.5 | TGCGTCCGGGATCAAG | 5.90E-04 | N | Putative membrane protein |
| 461029 | 312 | EC042_0414 | Uncommon | 72.5 | TGTGCGCAAGATCACA | 9.24E-07 | - | Conserved hypothetical protein |
| 4051417 | 311 | EC042_3818 | *yhjE* | -144.5 | TGTGAAGCATTTCATA | 2.83E-04 | N | MHS family MFS transporter |
| 4062646 | 308 | *dctA* | *dctA* | -132.5 | TTTGAGCTGGCTCGCA | 2.10E-05 | Y | C4-dicarboxylate transport protein |
| 172 | 301 | *thrA* | *thrA* | -218.5 | ATTGACTTAGGTCACT | 5.71E-04 | N | Bifunctional aspartokinase I/homoserine dehydrogenase I |
| 3899333 | 300 | *malT* | *malT* | -131.5 | TGTGACAGAGTGCAAA | 7.56E-05 | Y | Regulatory protein |
| 4046225 | 299 | *treF* | *treF* | -92.5 | CGTGATCTACCGCACG | 1.65E-05 | N | Cytoplasmic trehalase |
| 4272121 | 295 | EC042_4013 | *nepI* | -15.5 | TGTGACGCATTTAACG | 1.68E-04 | N | Purine ribonucleoside efflux pump NepI |
| 4412462 | 285 | *hdfR* | *hdfR* | 1050.5 | CACGCAGGGGGTCGCG | 7.32E-03 | N | LysR-family transcriptional regulator (H-NS-dependent *flhD* regulator) |
| 1475788 | 284 | *insB* | *lapA* | 140.5 | GGTGACAGCAGGCTCA | 2.08E-03 | N | Putative membrane protein |
| 3872334 | 283 | *nudE* | *nudE* | -42.5 | TGCGATATAGGACACG | 1.18E-04 | N | ADP compounds hydrolase |
| 2079650 | 271 | *pphA* | pseudo | -22.5 | CGCGCTAAAGATCACA | 2.78E-05 | - | Conserved hypothetical protein (pseudogene) |
| 4720171 | 269 | *psiE* | *psiE* | -54.5 | ATAGATCTCCGTCACA | 7.70E-04 | Y | Putative phosphate starvation-inducible membrane protein |
| 4522062 | 267 | *engB* | *engB* | 950.5 | TGTGATGGCTATTAGA | 6.98E-04 | N | Probable GTP-binding protein |
| 4747362 | 265 | *aphA* | *aphA* | -119.5 | CCTGCTTTTCATCACA | 1.81E-04 | N | Class B acid phosphatase |
| 3687170 | 262 | *argG* | *argG* | -242.5 | AGTGATCCACGCCACA | 2.10E-05 | Y | Argininosuccinate synthetase |
| 3545591 | 258 | *icc* | *icc* | -174.5 | TGCTGGCTTGAACACA | 2.63E-03 | N | Repressor protein of division inhibition gene |
| 2647958 | 252 | *pmrD* | *pmrD* | -94.5 | TGAGAAGTGAAACGGA | 1.08E-02 | N | Polymyxin B resistance protein |
| 3510478 | 251 | *hyb0* | *hyb0* | -77.5 | TGCGCCATTTACCACA | 1.81E-04 | N | Hydrogenase-2 small chain |
| 4217753 | 250 | EC042_3975 | Uncommon | -33.5 | CGTGTATACGATAACA | 2.48E-03 | - | Hypothetical protein |
| 2844005 | 245 | *hyfA* | *hyfA* | -190.5 | CGTGATCAAGATCACA | 1.79E-07 | Y | Hydrogenase-4 component A |
| 3160614 | 242 | EC042_2967 | *ygcW* | -57.5 | TGTGATCGTAATCACA | 2.51E-07 | N | Putative short chain dehydrogenase |
| 147627 | 240 | *gcd* | *gcd* | -69.5 | TGTGATCGTCATCACA | 7.11E-08 | Y | Quinoprotein glucose dehydrogenase |

| | | | | | | | | |
|---|---|---|---|---|---|---|---|---|
| 4353137 | 229 | *cat* | In K-12 | -103.5 | TGAGACGTTGATCGGC | 2.71E-03 | - | Chloramphenicol acetyltransferase |
| 1344062 | 218 | *adhE* | *ychH* | -230.5 | CGTGATCCAAATCAAA | 1.30E-05 | Y | Putative membrane protein |
| 2833992 | 209 | EC042_2670 | Uncommon | 230.5 | TGTGATACACAGCAAC | 3.15E-04 | - | Conserved hypothetical protein |
| 4006081 | 208 | EC042_3779 | *dtpB* | -62.5 | TGTAAACTTTTTCGCG | 1.55E-03 | N | Putative oligopeptide transporter |
| 3784875 | 200 | EC042_3554 | *yhdZ* | 928.5 | CGCTACTGCCGCCAGA | 8.30E-03 | N | ABC transporter, ATP-binding protein |
| 4752086 | 197 | *ssb* | *ssb* | -89.5 | CGGAACCGAGGTCACA | 7.70E-04 | N | Single-stranded binding protein |
| 2941936 | 195 | *glnB* | *glnB* | -40.5 | TCTGCTAAACGTAACA | 1.10E-03 | N | Nitrogen regulatory protein p-II |
| 167375 | 194 | *sfsA* | *sfsA* | 72.5 | ATCGGGTGTGATCACA | 2.34E-03 | Y | Sugar fermentation stimulation protein |
| 33766 | 193 | *rihC* | *rihC* | -72.5 | CGTGAAGTCGATTAAG | 2.21E-03 | N | Nonspecific ribonucleoside hydrolase (purine/pyrimidine ribonucleoside hydrolase) |
| 3562340 | 187 | EC042_3340 | In K-12 | -140.5 | TTTGAGTTCGCACCCA | 4.96E-03 | - | Conserved hypothetical protein |
| 2310574 | 176 | *amn* | *amn* | 186.5 | TTGGGATGGTAGCACA | 8.30E-03 | N | AMP nucleosidase |
| 4926886 | 174 | *set1A* | Uncommon | 48.5 | ACTGACGGTTTTCCCA | 6.61E-03 | - | Enterotoxin 1 |
| 1444796 | 173 | EC042_1404 | *stfR* | -17.5 | CTCTAACCACATAACG | 1.65E-02 | N | Phage side tail fiber protein |
| 5007735 | 171 | EC042_4670 | *bsmA* | -53.5 | TGTTACCTGGTACGCG | 1.74E-03 | N | Putative lipoprotein |
| 608383 | 168 | EC042_0536 | Uncommon | -120.5 | TGTGATGTAAAGCGCA | 4.09E-06 | - | Putative adhesin (not virulence) |
| 4089215 | 168 | *dppA* | pseudo | -500.5 | AATGGTTTCTGTCACA | 3.30E-03 | - | Conserved hypothetical protein |
| 3739372 | 164 | *sspA* | *sspA* | -174.5 | TGCGACCTTTGTGGTG | 1.54E-02 | N | Stringent starvation protein A |
| 2678707 | 161 | *lrhA* | *lrhA* | -178.5 | CCTGAAGTAGATCACA | 2.65E-05 | N | NADH dehydrogenase operon transcriptional regulator |
| 4448062 | 161 | *aslB* | *aslB* | -277.5 | AGTCCCCCCCCTCGCA | 3.39E-03 | N | Probable arylsulfatase-activating protein |
| 3740611 | 156 | *rplM* | *rplM* | -208.5 | TGTGATTTGTGGCAGG | 2.83E-04 | Y | 50S ribosomal subunit protein L13 |
| 3196847 | 155 | *fucP* | *fucP* | 548.5 | TGCCACATCAATCGCA | 7.21E-04 | Y | L-fucose permease |
| 5081393 | 155 | EC042_4739 | *ahr* | -129.5 | TGCGAGCAAGCTGGCG | 1.25E-03 | N | NADPH-dependent aldehyde reductase Ahr |
| 2272380 | 151 | EC042_2206 | *mtfA* | -181.5 | TGTGATTTTTGTCACT | 2.65E-05 | N | DgsA anti-repressor MtfA |
| 2068782 | 150 | *manX* | *sdaA* | -155.5 | ACGGATCTTCATCACA | 1.06E-03 | N | L-serine ammonia-lyase |

| | | | | | | | | |
|---|---|---|---|---|---|---|---|---|
| 1742072 | 146 | *sotB* | *uxaB* | -83.5 | CGCGATCCAGATCACA | 6.52E-07 | Y | Altronate oxidoreductase |
| 4237814 | 145 | EC042_3993 | In K-12 | -334.5 | CGTGCTACCGGTCACG | 3.98E-05 | - | Phage integrase family (tyrosine-type recombinase/integrase) |
| 1881643 | 144 | EC042_1835 | In K-12 | -123.5 | TTCGAGTCCACTCGGA | 4.00E-03 | - | Conserved hypothetical protein |
| 968344 | 134 | *clpS* | *ybiT* | -124.5 | TGTGACAGATGTCGCT | 8.90E-05 | Y | ABC transporter ATP-binding protein |
| 1038450 | 134 | *focA* | *clpS* | 135.5 | TGCGATTGAGATGAAA | 6.98E-04 | N | ATP-dependent Clp protease adaptor protein |
| 780664 | 122 | *nagE* | *nagE/nagB* | -165.5 | GGTGACAAAACTCACA | 5.87E-05 | Y | N-acetylglucosamine-specific PTS enzyme IIABC component/ Glucosamine-6-phosphate deaminase |
| 4287533 | 122 | *ivbL* | *ivbL* | -98.5 | TGAGGGGTTGATCACG | 3.51E-04 | Y | *ilvBN* operon leader peptide |
| 1072066 | 118 | EC042_1043 | *serC* | 22.5 | TGTGAACTCCGTCAGG | 4.95E-05 | Y | 3-phosphoserine (phosphohydroxythreonine transaminase) |
| 2576006 | 117 | *napF* | *napF* | -124.5 | CCCGATCGGGGTAAAA | 3.89E-03 | N | Ferredoxin-type protein |
| 3417179 | 116 | *mchS4* | Uncommon | 13.5 | TGTGATAATAATCACA | 2.08E-06 | - | Conserved hypothetical protein |
| 2197691 | 115 | EC042_2113 | *yedP* | -110.5 | TGTGACGCGCGTCACC | 6.04E-06 | N | Putative mannosyl-3-phosphoglycerate phosphatase |
| 1825856 | 114 | *mlc* | *mlc* | -97.5 | TGTGATTAACAGCACA | 2.21E-06 | Y | Protein Mlc (making large colonies protein) |
| 3364989 | 113 | *galP* | *galP* | -71.5 | TGTGATTTGCTTCACA | 7.00E-07 | Y | Galactose-proton symporter (galactose transporter) |
| 461937 | 112 | *phoA* | *phoA* | 172.5 | CGTGATTCTCTTAGCG | 1.45E-03 | N | Alkaline phosphatase |
| 263739 | 111 | EC042_0225 | Uncommon | -80.5 | TGTGAGCCGCATCACA | 1.15E-07 | - | Putative type VI secretion system protein |
| 4116976 | 107 | *malS* | *malS* | -61.5 | TGAGAGTTGAATCTCA | 1.03E-03 | Y | Alpha-amylase |
| 4160841 | 104 | *lctP* | *lctP* | -46.5 | GGAGATGAGCATCAGA | 1.74E-03 | N | L-lactate permease |
| 5167710 | 103 | EC042_4823 | pseudo | 14.5 | TGATATATAGATAAGA | 1.28E-02 | - | Conserved hypothetical protein |
| 544222 | 102 | *maa* | *maa* | -86.5 | TGTGATAAAGATCACA | 1.34E-07 | N | Maltose O-acetyltransferase |
| 2164036 | 99 | EC042_2071 | *yecA* | -339.5 | TTTGCAATCCGCTACA | 8.30E-03 | N | Conserved hypothetical protein |
| 3353695 | 98 | EC042_3140 | *yggP* | 472.5 | TGTGATAATTGGCATG | 1.91E-03 | N | Zinc-binding dehydrogenase |
| 263036 | 97 | EC042_0224 | Uncommon | 62.5 | TTTGGTGTATACCGAA | 2.95E-03 | - | Putative type VI secretion system protein |

| | | | | | | | | |
|---|---|---|---|---|---|---|---|---|
| 1868607 | 95 | EC042_1802A | *malX/malI* | 51.5 | CGTGATCAAGATCACG | 1.44E-06 | Y | PTS maltose transporter subunit IIC/ *mal* regulon transcriptional regulator MalI |
| 2214844 | 95 | EC042_2130 | Uncommon | 814.5 | AGTGCTGAATGTCACA | 6.13E-05 | - | Putative prophage protein |
| 4101165 | 95 | *cspA* | *cspA* | -32.5 | ATCGCCGAAAGGCACA | 3.21E-03 | N | RNA chaperone/antiterminator CspA |
| 2544035 | 94 | EC042_2414 | *yejG* | -147.5 | TGCGGGCGTGGTCACG | 5.87E-05 | N | YejG family protein |
| 1740383 | 90 | *uxaB* | *uxaB* | -134.5 | TGTGGCGCGGATCATG | 3.77E-04 | - | Putative membrane protein |
| 2054834 | 90 | *sdaA* | *yeaV* | -142.5 | TGAGACAATCATCGCA | 1.68E-04 | N | BCCT family transporter YeaV |
| 4812398 | 90 | *melR* | *melR* | -63.5 | TGCGAGTGGGAGCACG | 5.63E-05 | Y | Melibiose operon regulatory protein |
| 379869 | 89 | EC042_0340 | In K-12 | -186.5 | TGTGAGCGATGCCGAA | 2.53E-04 | - | Conserved hypothetical protein |
| 4530302 | 86 | *typA* | *glnA* | -114.5 | CGTGAAAGCGATCACA | 6.68E-06 | Y | Glutamine synthetase |
| 4969712 | 85 | EC042_4632 | *yjeM* | -137.5 | TCCGCTAAAGGCCACA | 7.21E-04 | N | Putative permease |
| 1623270 | 82 | EC042_1630 | *ydcH* | -287.5 | TGTGATGAATGTCACT | 1.36E-05 | N | YdcH family protein |
| 2559209 | 81 | EC042_2431 | Uncommon | -285.5 | TGATCCCCCATTAACG | 2.94E-02 | - | Putative prophage protein |
| 2982786 | 81 | *kgtP* | *kgtP* | -1116.5 | TGCTCGCGCCGTCACG | 7.21E-04 | N | Alpha-ketoglutarate permease |
| 2776365 | 79 | EC042_2617 | *fixA*-like family protein | -283.5 | TGTGAAGTACCGAAGT | 7.32E-03 | N | Conserved hypothetical protein |
| 3080457 | 78 | EC042_2887 | *yqaB* | -573.5 | TCCGGGAGGATTCGAA | 1.50E-02 | N | Putative phosphatase |
| 3332837 | 78 | EC042_3122 | *fau* | -118.5 | GGTGAGCATGCTCGGT | 4.83E-03 | N | Conserved hypothetical protein |
| 56171 | 77 | *folA* | *folA* | -178.5 | AGTGACGTAAATCACA | 5.40E-06 | N | Dihydrofolate reductase |
| 937104 | 77 | EC042_0886 | *ybhQ* | -100.5 | TATGCGCTGCGTCACA | 1.33E-04 | N | Putative membrane protein |
| 1970534 | 77 | EC042_1890 | *infC* | -68.5 | TGTGCGTTAGCTCGTG | 1.85E-03 | N | Translation initiation factor IF-3 |
| 4456964 | 77 | *cyaA* | *cyaA* | -161.5 | TGTTAAATTGATCACG | 1.49E-04 | Y | Adenylate cyclase |
| 3346695 | 76 | *mscS* | *mscS* | -102.5 | TGCCAAATAGATCACA | 1.88E-04 | N | Small-conductance mechanosensitive channel |
| 1418735 | 75 | *stfR* | In K-12 | -112.5 | TGTTATGATACGCAGG | 5.37E-03 | - | Putative phage protein |
| 4650149 | 75 | *tufA* | *tufA* | -123.5 | TCTGCCTATCAGCACC | 2.79E-03 | N | Elongation factor Tu |
| 4167598 | 74 | *secB* | *grxC* | -135.5 | TGTGCTGTGCGTCAAT | 1.95E-04 | N | Glutaredoxin 3 |
| 4325818 | 71 | EC042_4064 | *tnaA* | -316.5 | TGTGATTCGATTCACA | 2.47E-06 | Y | Tryptophanase |

| | | | | | | | | |
|---|---|---|---|---|---|---|---|---|
| 855435 | 69 | *moaA* | *aroG* | 2.5 | TGTACATGGCTTCACA | 9.67E-04 | N | Phospho-2-dehydro-3-deoxyheptonate aldolase, Phe-sensitive |
| 3407089 | 69 | EC042_3190 | Uncommon | 62.5 | CTCTCTGTCAGCCACT | 3.81E-02 | - | Conserved hypothetical protein |
| 4734493 | 69 | *plsB* | *plsB/dgkA* | 8.5 | CGTGGCCAGCCGGACA | 2.71E-03 | N | Glycerol-3-phosphate acyltransferase PlsB/ diacylglycerol kinase |
| 150878 | 67 | EC042_0128 | *yadI* | -59.5 | TTTGACGGCTATCACC | 1.81E-04 | N | PTS sugar transporter subunit IIA |
| 4511085 | 67 | *mobB* | *mob* | 2473.5 | AGTGCGAATGCTGACA | 5.66E-03 | N | Molybdopterin-guanine dinucleotide biosynthesis protein B |
| 4664179 | 66 | *htrC* | *yjaZ* | 76.5 | CTTGATGCTCATCAGG | 8.23E-04 | N | Heat shock protein C |
| 3751263 | 64 | *aaeR* | *aaeR* | -26.5 | TGTGATCTAAATCACT | 2.63E-06 | Y | LysR-family transcriptional regulator |
| 4646221 | 64 | *murB* | *murB* | -244.5 | CCTGGCGGCCGTAGCG | 1.19E-02 | N | UDP-N-acetylenolpyruvoylglucosami-ne reductase |
| 1748762 | 63 | EC042_1741 | *ydeA* | -130.5 | TGTTAACCCTGCAACA | 3.30E-03 | N | L-arabinose MFS transporter |
| 816974 | 62 | *sdhC* | *sdhC* | -145.5 | AGTGTTTTGCATGACG | 9.16E-03 | Y | Succinate dehydrogenase cytochrome b-556 subunit |
| 816974 | 62 | *cydA* | *sdhC* | -134.5 | TGTAACTTTTTTATCA | 1.97E-02 | Y | Succinate dehydrogenase cytochrome b-556 subunit |
| 175121 | 61 | *fhuC* | *fhuC* | -586.5 | TGTGGTGACTGGCGCA | 1.38E-04 | N | Ferrichrome transport ATP-binding protein |
| 3209376 | 59 | *mltA* | *mltA* | -153.5 | CGTGATCGGGGTAAAA | 2.10E-04 | N | Murein transglycosylase A |
| 5129030 | 57 | EC042_4783 | *fecI* | -92.5 | TCTGCTATTATTGACA | 5.81E-03 | N | RNA polymerase sigma factor |
| 773400 | 56 | EC042_0701 | In K-12 | -480.5 | CTTGAACTCGCACACC | 4.58E-03 | - | Putative exported protein (pseudo) |
| 2499767 | 56 | EC042_2378 | *preT* | -100.5 | TGTGAATCCTTTCACA | 1.36E-05 | Y | Putative oxidoreductase |
| 2095051 | 55 | *torY* | *pphA* | 470.5 | TTTGATAAACCTTGCA | 1.74E-03 | N | Serine/threonine protein phosphatase 1 |
| 3892798 | 55 | *gntT* | *gntT* | -190.5 | TCTGGTGATTCTCAAA | 2.08E-03 | Y | High-affinity gluconate transporter |
| 3966675 | 53 | EC042_3740 | In K-12 | 134.5 | TGTGGGATACTTCCCG | 1.10E-03 | - | Putative acyltransferase |
| 5101081 | 53 | EC042_4755 | In K-12 | -80.5 | TGTGACTGAGATCGCG | 4.35E-06 | - | Putative sugar kinase |
| 3128509 | 52 | *rpoS* | *nlpD* | -511.5 | TGTGACCGTGGTCGCA | 5.71E-07 | Y | Murein hydrolase activator NlpD |
| 2677252 | 50 | *nuoA* | *nuoA* | -291.5 | TGTGAAGCAATGGAAA | 4.34E-03 | N | NADH-quinone oxidoreductase subunit A |

| | | | | | | | | |
|---|---|---|---|---|---|---|---|---|
| 3932650 | 49 | *gntR* | *gntR* | -42.5 | GTTTAACACGGACGCA | 2.54E-02 | Y | Gluconate utilization operon repressor |
| 242043 | 48 | EC042_0209 | In K-12 | -417.5 | CGTTCCCAACGGAACA | 4.58E-03 | - | Putative lipoprotein |
| 3011421 | 48 | EC042_2820 | In K-12 | -414.5 | CGTAAAAAGCCGCAAA | 1.06E-02 | - | Integrase |
| 5207449 | 48 | EC042_4865 | *yjjZ* | 405.5 | TGCGAGGGGGGGGACT | 3.39E-03 | N | Putative membrane protein |
| 40727 | 46 | *caiF* | *caiF* | -123.5 | CAGGATTTAGCTCACA | 1.80E-03 | Y | Transcriptional activator |
| 4359930 | 46 | *aadA1* | Uncommon | 102.5 | TTTGTACGGCTCCGCA | 4.71E-03 | - | Aminoglycoside adenylyltransferase |
| 3608027 | 45 | EC042_3380 | *ygjR* | 48.5 | TACGAACTGGATCACC | 7.21E-04 | N | Putative oxidoreductase |
| 4374307 | 45 | *tetR* | Uncommon | 78.5 | TTTGCGTGTCGTCAGA | 1.33E-03 | - | Tetracycline repressor |
| 5230955 | 45 | EC042_4888 | ettA/sltY | -90.5 | GTCGATCACCTTCGCA | 1.06E-03 | N | Energy-dependent translational throttle protein EttA/ murein transglycosylase |
| 2917822 | 42 | *EC042_2736* | *trmJ* | -33.5 | CGCGCATCTTATCATA | 5.10E-03 | N | Putative RNA methyltransferase |
| 3204810 | 42 | *gcvA* | *gcvA* | -234.5 | TGCGATTCAGACCATG | 1.03E-03 | N | Glycine cleavage system transcriptional activator |
| 4674584 | 42 | *hupA* | *hupA* | -50.5 | TGGCATTTCCGTCGCA | 2.02E-03 | Y | DNA-binding protein HU-alpha |
| 5121039 | 42 | EC042_4775 | In K-12 | 523.5 | TGTCCGCTCTGGCACA | 9.67E-04 | - | Hypothetical protein |
| 1976942 | 41 | *msrB* | *yniA* | -126.5 | CGTGATGAAAATCACA | 1.74E-06 | N | Fructosamine kinase family protein |
| 4978451 | 41 | *orn* | *orn* | 1081.5 | AGTGCTGTAAAGCACA | 9.67E-05 | N | Oligoribonuclease |
| 5015959 | 41 | EC042_4681 | *rpsF* | -102.5 | TGTCAGTACTATAACG | 7.32E-03 | Y | 30S ribosomal subunit protein S6 |
| 125271 | 40 | *nadC* | *ampD* | 31.5 | GTTGGCGCGCGCCGCG | 7.89E-03 | Y | N-acetyl-anhydromuramyl-L-alanine-amidase |
| 3284051 | 40 | EC042_3081 | *ygeW* | -157.5 | TGTGATCAACCCCACA | 3.11E-06 | N | Putative aspartate/ornithine carbamoyltransferase |
| 841420 | 39 | EC042_0769 | *cydA* | -327.5 | AGCGTGAAGGATAACG | 1.80E-02 | N | Cytochrome d ubiquinol oxidase subunit 1 |
| 851102 | 39 | EC042_0773 | Uncommon | -123.5 | CTTGAAATAATTAACA | 2.71E-03 | - | Hypothetical protein |
| 2051133 | 39 | EC042_1966 | *yeaQ* | -227.5 | TTTGAACCGCGTCACT | 1.28E-04 | N | Putative transglycosylase associated protein |
| 2507357 | 39 | *galS* | *galS* | -84.5 | CGTGAATCGAGTCACA | 1.50E-05 | Y | HTH-type transcriptional regulator GalS |
| 4817898 | 39 | *dcuB* | *dcuB* | -42.5 | TGCTGAATAGATCACA | 3.90E-04 | N | Anaerobic C4-dicarboxylate transporter |
| 3308819 | 37 | EC042_3098 | *ygfT* | -104.5 | TGCGACTGAGTTCAAA | 4.95E-05 | N | Formate-dependent uric acid utilization protein YgfT |

| | | | | | | | | |
|---|---|---|---|---|---|---|---|---|
| 1915871 | 36 | *lpp* | *ydhR* | -45.5 | TGTAATACTTGTAACG | 4.23E-03 | N | Conserved hypothetical protein |
| 4466034 | 35 | EC042_4187 | Uncommon | -7.5 | CGAGGTGTTGATCACG | 5.15E-04 | - | Hypothetical protein |
| 409039 | 34 | EC042_0364 | In K-12 | -30.5 | TGCGAGAGAGATCACA | 2.21E-06 | - | LysE-family translocator |
| 2255919 | 34 | EC042_2180 | pseudo | 556.5 | TGTGAAGTATTTAAAA | 7.21E-04 | - | Putative prophage protein |
| 4202532 | 33 | EC042_3956 | Uncommon | 435.5 | GGCGACATATGACGCG | 5.10E-03 | - | Putative prophage protein |
| 4945596 | 33 | EC042_4606 | Uncommon | 753.5 | CTTGATGGAGCACAGA | 3.39E-03 | - | Hypothetical protein |
| 2766013 | 32 | EC042_2608 | *glk/ yfeO* | -98.5 | ATCGATCTGGGTCACA | 7.25E-05 | N | Glucokinase/Ion channel protein |
| 3985355 | 32 | EC042_3762 | *gntR* | -129.5 | ATCGACTCACGTCACA | 2.93E-04 | Y | GntR-family transcriptional regulator |
| 4451798 | 32 | *aslA* | *aslA* | -415.5 | TGTGGCATAATAAACG | 9.16E-03 | N | Arylsulfatase |
| 4005281 | 31 | *uspA* | *uspA* | -148.5 | GATCATCCGGGTCGCT | 3.06E-02 | N | Universal stress protein A |
| 4290090 | 31 | EC042_4031 | *yidG* | -79.5 | CGCGAGGGAGATCAAA | 6.13E-05 | N | Putative membrane protein |
| 1865552 | 30 | *uidR* | *uidA* | 17.5 | TGTGCTTCAGTCTGCA | 2.15E-03 | Y | Glucuronidase ß |
| 2496355 | 30 | EC042_2373 | *yohj* | -57.5 | TGTGATCGGTAGCACG | 8.25E-06 | N | Putative membrane protein |
| 2782302 | 30 | *zipA* | *zipA* | -70.5 | TTTGCCGATTACCTCA | 4.96E-03 | N | Cell division protein |
| 405459 | 29 | EC042_0361 | *yahK* | -161.5 | TGTGATCTGCCACGAA | 3.51E-04 | N | NADPH-dependent aldehyde reductase YahK |
| 526093 | 29 | *ppiD* | *ppiD* | -77.5 | CCCGTTTCTTGTCACA | 4.11E-03 | N | Peptidyl-prolyl cis-trans isomerase D |
| 1139173 | 29 | *cspG* | *yccA* | -26.5 | TTTGCCGAAAGGCCCA | 2.41E-03 | N | Putative membrane protein |
| 3485206 | 29 | *glcC* | *glcC* | -10.5 | TGTGCACGAGGTCCGG | 1.80E-03 | Y | Glc operon transcriptional activator |
| 540752 | 28 | EC042_0492 | *ybaA* | -170.5 | TGTTGGTTCTCTCGCA | 1.64E-03 | N | Conserved hypothetical protein |
| 3186304 | 27 | EC042_2990 | *yqcC* | -182.5 | TTGCTTCCTGCTCACA | 2.44E-02 | N | Conserved hypothetical protein |
| 943255 | 26 | EC042_0905 | *cecR* | -179.5 | TGCGAAGGGGATTGCA | 3.77E-04 | N | TetR-family transcriptional regulator |
| 3548332 | 26 | EC042_3327 | *ygiB* | -88.5 | CGGGTTATCTGCCGCA | 1.34E-02 | N | Putative lipoprotein |
| 4219192 | 26 | EC042_3978 | *yicG* | -156.5 | TATGCATTTTCTCAGA | 8.93E-03 | N | Putative membrane protein |
| 2315229 | 25 | EC042_2225 | *yeeO* | -68.5 | TTCGAGTCCAGTCAGA | 1.25E-03 | N | Putative membrane protein |
| 2349836 | 25 | *sbcB* | *dacD/sbcB* | -72.5 | TGTGACTACTATCTCA | 8.56E-05 | N | Penicillin-binding protein 6B/ exodeoxyribonuclease I |
| 4968382 | 25 | *frdA* | *frdA* | -76.5 | TGCGAACGCTATTCCA | 3.49E-03 | N | Fumarate reductase flavoprotein subunit |
| 2551890 | 24 | EC042_2422 | *intA* | -239.5 | GGTGTCGGGGGTCGGA | 3.12E-03 | N | Integrase |
| 3276979 | 24 | EC042_3075 | Uncommon | -236.5 | GGCGAAGGGAATCGAA | 1.69E-03 | - | Conserved hypothetical protein |

| | | | | | | | | |
|---|---|---|---|---|---|---|---|---|
| 3363224 | 24 | EC042_3148 | *yqgD* | 133.5 | TGAGACACGATTCAAA | 9.37E-04 | N | Putative membrane protein |
| 4697783 | 24 | *iclR* | *metH* | -96.5 | TGTTGAACAAATCTCA | 4.58E-03 | N | Methionine synthase (5-methyltetrahydrofolate-homocysteine methyltransferase) |
| 2733499 | 23 | EC042_2583 | *yfcZ/fadL* | -113.5 | AGTGACCGAAATCACA | 5.71E-06 | Y | Conserved hypothetical protein/ Long-chain fatty acid transport protein |
| 4243275 | 23 | EC042_3998 | Uncommon | -20.5 | CGCAATCAACGCCACA | 8.78E-04 | - | Putative phage immunity repressor protein |
| 2273560 | 22 | EC042_2207 | In K-12 | -395.5 | TTCGAGTCCAGTCAGA | 1.25E-03 | - | Integrase |
| 2685148 | 22 | EC042_2536 | *yfbV* | -156.5 | TTGGCTGAAAATTACG | 1.84E-02 | N | Putative membrane protein |
| 4301258 | 22 | EC042_4041 | *yidE* | -71.5 | TTTGCCTTATAGCGCA | 4.65E-04 | N | Putative transporter |
| 5010825 | 22 | *ulaA* | *ulaA* | -113.5 | TGCGGGTCGCGTCACA | 3.81E-05 | Y | PTS ascorbate transporter subunit IIC |
| 4236685 | 21 | EC042_3992 | In K-12 | -169.5 | TGTGATCTTCCGCCAA | 5.71E-04 | - | Tyrosine-type recombinase/integrase |
| 2132403 | 20 | *flhD* | *cutC* | -270.5 | TGTGATGCAGATCACA | 7.72E-09 | N | Copper homeostasis protein |
| 3590224 | 20 | *air* | *air* | -93.5 | TTTGATTTAGATCGCA | 8.25E-06 | Y | Aerotaxis receptor protein |
| 3690953 | 20 | EC042_3465 | Uncommon | -68.5 | TTTGGTACCGAGGACG | 8.93E-03 | - | Hypothetical protein |
| 3723245 | 20 | *gltB* | *gltB* | -188.5 | TTTGCGCTAAAGCACA | 1.14E-04 | Y | Glutamate synthase [NADPH] large subunit |
| 3948285 | 20 | EC042_3720 | *panM* | -204.5 | CGTTGTCAGCATAAAA | 1.01E-02 | N | Putative acetyltransferase |
| 965159 | 19 | EC042_0908 | *opgE* | -28.5 | TGCGCGGTTTGTCATA | 1.13E-03 | N | Putative membrane protein |
| 2158975 | 18 | EC042_2067 | *ftnB* | -171.5 | TGTGATGTAAATCACA | 5.37E-08 | N | Ferritin-like protein 2 |
| 3512048 | 18 | EC042_3292 | *gpr* | -76.5 | TGTGAGCCAGACTACG | 3.15E-04 | N | Putative aldo/keto reductase |
| 1926713 | 17 | *infC* | *lpp* | -853.5 | CGTTAACTTCATCGCG | 5.90E-04 | N | Major outer membrane lipoprotein |
| 3640881 | 17 | *garK* | *garK* | 1483.5 | CGCTCACTGGCTCAAG | 7.70E-03 | N | Glycerate kinase 2 |
| 3858617 | 17 | EC042_3644 | In K-12 | -83.5 | GGCGAGGCGCTTCACA | 9.67E-05 | - | Hypothetical protein |
| 4431631 | 17 | *rho* | *rho* | -210.5 | TGTAATTTCCAACGCT | 8.30E-03 | N | Transcription termination factor |
| 336180 | 16 | EC042_0305 | Uncommon | 938.5 | TCCACCGTGATTCACG | 1.14E-02 | - | Conserved hypothetical protein |
| 3327528 | 16 | *gcvT* | *gcvT* | -87.5 | CCCGGTCCCCAACGCA | 7.32E-03 | Y | Aminomethyltransferase (glycine cleavage system protein |
| 4182076 | 16 | *rfaK* | *rfaK* | 658.5 | TGTGAGCAACGGCCAA | 6.53E-04 | N | Lipopolysaccharide N-acetylglucosaminyltransferase |

| | | | | | | | | |
|---|---|---|---|---|---|---|---|---|
| 4318893 | 16 | *dnaA* | *dnaA* | -189.5 | TCTTCTGTTTCTCACA | 1.69E-03 | N | Chromosomal replication initiator protein DnaA |
| 3846061 | 15 | *frlA* | *frlA* | -115.5 | TGTGATCTTCCTCCAC | 1.37E-03 | Y | Fructoselysine (psicoselysine) transporter |
| 2313387 | 14 | EC042_2224 | *yeeN* | 1185.5 | TCTGACTGGACTCGAA | 1.25E-03 | N | Conserved hypothetical protein |
| 3996354 | 14 | EC042_3771 | *ybhG* | -224.5 | TGTGATCTATAAAACT | 1.59E-03 | N | HlyD family secretion protein |
| 4944103 | 14 | EC042_4606 | Uncommon | -645.5 | GGCGGCGTTCAGCCCA | 4.71E-03 | - | Putative helicase (pseudogene) |
| 1302952 | 13 | *dadA* | *minC* | -105.5 | TGTGAGCCAGCTCACC | 7.43E-06 | N | Septum site determining protein MinC |
| 4192231 | 13 | *rpmB* | *rpmB* | -173.5 | TGTGCTCAAGTCCCGA | 2.79E-03 | N | 50S ribosomal subunit protein L28 |
| 4596821 | 13 | *cytR* | *cytR* | -136.5 | TTCGATCCGCCTCGCA | 5.87E-05 | Y | DNA-binding transcriptional regulator |
| 5093367 | 13 | EC042_4746 | Uncommon | -657.5 | GGAGAGCGACTCTGCA | 3.32E-02 | - | Conserved hypothetical protein |
| 27718 | 12 | *ribF* | *rpsT* | -90.5 | TGTGCAAATAAGCGCC | 2.71E-03 | N | 30S ribosomal protein S20 |
| 784980 | 12 | EC042_0709 | *In K-12* | -85.5 | AGCGAGACTTTTCTCA | 4.96E-03 | Y | Putative exported protein |
| 1127654 | 12 | EC042_1056 | In K-12 | -273.5 | GGTGCCTTCAACCGCT | 5.66E-03 | - | Conserved hypothetical protein |
| 4956885 | 12 | EC042_4617 | *groS* | -136.5 | TTTTGTGCTGATCAGA | 3.79E-03 | N | 10 kDa chaperonin |
| 4354065 | 11 | EC042_4087A | *In K-12* | 351.5 | TTTGCTCAGGCTCTCC | 3.39E-03 | - | Putative plasmid-related protein |
| 4852738 | 11 | EC042_4524 | Uncommon | -81.5 | TCCTATAATGATCAAA | 4.71E-03 | - | Putative type VI secretion protein |
| 3337434 | 10 | *sbm* | *scpA* | -136.5 | CTTGAATCACATCACA | 6.13E-05 | N | Methylmalonyl-CoA mutase |
| 4999291 | 10 | *EC042_4657* | *In K-12* | 37.5 | AGTAACGATAATCTCA | 7.70E-03 | - | Putative D-galactarate dehydratase/altronate hydrolase |
| 407967 | 9 | *EC042_0363* | Uncommon | 179.5 | TATGAAAAAAATCATA | 5.37E-03 | - | Hypothetical protein |
| 3187106 | 9 | *syd* | *syd* | 219.5 | TGTGGGTTGTATCACA | 1.57E-05 | N | Putative SecY-interacting protein |
| 3883094 | 9 | *greB* | *greB* | 102.5 | GGTGACCTGGGCCGCA | 5.40E-05 | N | Transcription elongation factor GreB |
| 4177213 | 9 | *rfaD* | *htrL* | -115.5 | TTTGTTGCAATTAGCA | 7.51E-03 | N | Putative lipopolysaccharide biosynthesis protein |
| 927587 | 8 | EC042_0880 | *moaA* | -45.5 | TGTAGTCGGCGTCACA | 1.28E-04 | N | Molybdenum cofactor biosynthesis protein A |
| 1149859 | 8 | *putP* | *cspG* | -196.5 | TGTGAGAGAGTGCAAC | 1.10E-03 | N | Old shock-like protein |
| 4946926 | 8 | *insB* | *insA* | 57.5 | TTTGCCGTTACGCACC | 2.28E-03 | N | Transposase |

| | | | | | | | | |
|---|---|---|---|---|---|---|---|---|
| 1068899 | 7 | *serC* | *focA* | -144.5 | TGTGATGCAAGCCACA | 3.11E-06 | Y | Probable formate transporter 1 |
| 2525169 | 7 | EC042_2399 | *psuK* | -219.5 | TGTGACGGGATGCACA | 1.06E-05 | N | Putative pseudouridine kinase |
| 3419417 | 7 | *mchS1* | Uncommon | 905.5 | GTGGAGTGACAGCACT | 2.39E-02 | - | Putative microcin esterase |
| 4761274 | 7 | EC042_4436 | pseudo | -125.5 | GGTGATGCATTTTGCA | 1.85E-03 | - | Putative type III effector protein (pentapeptide repeat protein) (pseudogene) |
| 20866 | 6 | *insB* | *insB* | 140.5 | GGTGACAGCAGGCTCA | 2.08E-03 | N | IS1 transposase B |
| 516051 | 6 | *cyoA* | *cyoA* | -234.5 | TGCGAAATAAAACAAT | 7.32E-03 | Y | Cytochrome o ubiquinol oxidase subunit 2 |
| 2746629 | 6 | EC042_2591 | *insA* | -11.5 | CGCGCAGACGATGACG | 4.46E-03 | N | Putative transposase (IS1) |
| 2945729 | 6 | EC042_2760 | *glrK* | -163.5 | TTGGACGGCAGGCACC | 5.81E-03 | N | Two-component system sensor kinase |
| 128586 | 5 | *pdhR* | *pdhR* | -128.5 | TGTTAAAATGTGCACA | 6.98E-04 | Y | Pyruvate dehydrogenase complex repressor |
| 4260185 | 5 | EC042_4011 | Uncommon | 2631.5 | GCGGACTCACTTCACC | 1.65E-02 | - | Transcriptional regulator |
| 3565809 | 4 | EC042_3343 | In K-12 | 1818.5 | TCCGGCGTTCATCAAC | 6.28E-03 | - | Putative membrane protein |
| 4213344 | 4 | EC042_3970 | Uncommon | 308.5 | TTTGCTCTCATCCACA | 3.15E-04 | - | Putative prophage protein |
| 4266116 | 4 | EC042_4012 | Uncommon | 4608.5 | CGCGACAAACGTCACG | 4.95E-05 | - | Putative invasion ‘air’ |
| 5141146 | 4 | EC042_4800 | In K-12 | -169.5 | TATGAATGAAAGAACA | 5.81E-03 | - | Conserved hypothetical protein |
| 5240772 | 4 | *arcA* | *arcA* | -88.5 | CATGATCGGCGTAACA | 1.17E-03 | N | Aerobic respiration control protein |
| 1235648 | 3 | *ptsG* | *rluC* | -142.5 | CGTGATAGCCGTCAAA | 5.17E-05 | N | Ribosomal large subunit pseudouridine synthase C |
| 1601165 | 3 | EC042_1553 | *insA1* | -85.5 | CACGATCCCGCTCGCA | 6.75E-04 | N | Transposase |
| 2842820 | 3 | *dapA* | *dapA* | + 0.5 | CGTGAACATGGGCCAT | 1.16E-02 | N | Dihydrodipicolinate synthase |
| 4362407 | 3 | EC042_4096 | In K-12 | 279.5 | TCTGCACAAGCTCGCG | 6.75E-04 | - | Putative acetyltransferase |
| 5118152 | 2 | EC042_4771 | In K-12 | 105.5 | CCTGCAGACAAGCAGA | 1.03E-02 | - | Glycosyl transferase |
| 5133814 | 2 | EC042_4789 | In K-12 | 852.5 | AGCGAACCACACTGCA | 6.44E-03 | - | Transposase |
| 5089682 | 1 | EC042_4744 | Uncommon | -1205.5 | TTTCATATAGGCCGCG | 5.52E-03 | - | Conserved hypothetical protein |

| | | | | | | | |
|---|---|---|---|---|---|---|---|
| *Targets on plasmid pAA* | | | | | | | |
| 27988 | 2975 | *pet* | Uncommon | -117.5 | CGAGAGCATTGTCACA | 2.63E-04 | Serine protease autotransporter toxin Pet |
| 62001 | 907 | EC042_RS26580 | *traS* | -247.5 | TGTGACCATATTATCA | 1.33E-03 | Hypothetical protein |
| 18257 | 387 | *virK* | Uncommon | -235.5 | TGTGGTGACTGGTACA | 1.37E-03 | Virulence protein |
| 112767 | 321 | *repA* | In K-12 | -84.5 | ACAGATCTTCGTCACA | 1.25E-03 | Replication protein A |
| 108769 | 293 | *repA* | In K-12 | -84.5 | TGTGACGAAGATCTGT | 1.25E-03 | Replication protein A |
| 39765 | 81 | *aafA* | Uncommon | 159.5 | CGTTGACAGGAGCGCA | 2.41E-03 | Aggregative adherence fimbria II major subunit AafA |
| 94581 | 23 | EC042_RS26810 | In K-12 | -26.5 | GTTGACTGGAGCCGCA | 2.63E-03 | Hypothetical protein |
| 50078 | 15 | EC042_RS29140 | pseudo | -93.5 | TGTGAAATTAATCAAA | 5.40E-05 | IS66 family insertion sequence element accessory protein TnpB |
| 3987 | 13 | EC042_RS30500 | pseudo | 186.5 | TGAGCCCGCCATCACC | 5.15E-04 | Transposase |
| 86933 | 13 | EC042_RS26755 | *hok* | -115.5 | TCTGCCACACGACACG | 2.56E-03 | Type I toxin-antitoxin system Hok family toxin |

[a] Peak centre indicated by MACS2 (8) and annotated to EAEC 042 chromosome. Locations with more than one CRP site are highlighted in red.

[b] Score (int(-10*log10qvalue)) assigned by MACS2.

[c] Annotated gene to an identified CRP site.

[d] EAEC 042 homologues to *E. coli* K-12 genes. Genes underlined represent matching genes targeted by CRP and previously indicated by Grainger *et a l*(9) . “Uncommon” denotes genes not found in K-12. “In K-12" indicates genes that are present in *E. coli* K-12 but with unknown functions.

[e] Position of identified CRP site to nearest TSS (Translation Start Site).

[f] Matching sequence obtained from MEME SUITE (10).

[g] *p*-value for CRP motif matches from MEME SUITE (10).

[h] Reported to be regulated by CRP, as listed for *E. coli* K-12 in RegulonDB(11).

[i] Product of encoding gene (12).

### Table S3: A comparison of CRP-dependent promoters in *E. coli* K-12 and *E. coli* 042

| Promoter | Strain (gene name is provided if in 042) | Promoter sequence [a] | bp distance CRP to TSS [b] | bp spacing CRP site to -35 [b] | bp spacing CRP to 10[b] | bp spacing -35 to -10[b] | "+1" position (K-12) [a] | % Identity [c] |
|---|---|---|---|---|---|---|---|---|
| *exuTp2* | (K-12)<br>EC042_3387 | ATATTCCCACATTTGTGATGGCTCTCACCTTTTAAAGTTGTATGACAAGTTATCTTTCTGCCGTCGCAAATCATAAGTCGA<br>ATATTCCCACATTTGTGATGGCTCTCACCTTTTAAAGTTGTATGACAAGTTATCTTTCTGCCGTCGCAAATCATAAGTCGA | -39.5 | N/A | 30 | N/A | 3245002 | 100 |
| *lacZp2*<br>*lacZ* | (K-12)<br>EC042_0381 | GCAACGCAATTAATGTGAGTTAGCTCACTCATTAGGCACCCCAGGCTTTACACTTTATGCTTCCGGCTCGTATGTTGTGTG<br>GCAACGCAATTAATGTGAGTCAGCTCACTCATTAGGCACCCCAGGCTTTACACTTTATGCGTCCGGCTCGTATGTTGTGTG | -39.5 | N/A | 30 | N/A | 366365 | 97.5 |
| *acnAp2* | (K-12)<br>EC042_1401 | TCTTTTATCAATTTGGGTTGTTATCAAATCGTTACGCGATGTTTGTGTTATCTTTAATATTCACCCTGAAGAGAATCAGGG<br>TCTTTTATCAATTTGGGTTGTTATCAAATCGTTACGCGATGTTTGTGTTATCTTTAATATTCACCCTGAAGAGAATCAGGG | -40.5 | N/A | 29 | N/A | 1335781 | 100 |
| *deoCp2* | (K-12)<br>EC042_4878 | GATTTCCTTAATTGTGATGTGTATCGAAGTGTGTTGCGGAGTAGATGTTAGAATACTAACAAACTCGCAAGGTGAATTTTA<br>GATTTCCTTAATTGTGATGTATATCGAAGTGTGTTGCGGAGTAGATGTTAGAATACTAACAAACTCGCAAGGTGAATTTTA | -40.5 | N/A | 29 | N/A | 4617278 | 98.8 |
| *focAp1* | (K-12)<br>EC042_0994 | AGCCAGGCGAGATATGATCTATATCAATTTCTCATCTATAATGCTTTGTTAGTATCTCGTCGCCGACTTAATAAAGAGAGA<br>AGCCAGGCGAGATATGATCTATATCAATTTCTCATCTATAATGCTTTGTTAGTATCTCGTCGCCGACTTAATAAAGAGAGA | -40.5 | N/A | 30 | N/A | 954493 | 100 |
| *fucPp* | (K-12)<br>EC042_3000 | CTAGCTAATAAGTGTGACCGCCGTCATATTACAGAGCGTTTTTTATTTGAAAATGAATCCATGAGTTCATTTCAGACAGGC<br>CCAGCTAATAAGTGTGACCGCCGTCATATTACAGAGCGTTTTTTATTTGAAAATGAATCCATGAGTTCATTTCTGACAGGC | -40.5 | N/A | 29 | N/A | 2934119 | 97.5 |
| *gatY* | (K-12)<br>EC042_2328 | ATTGTCGTTTTTGTGATCGTTATCTCGATATTTAAAAACAAATAATTTCATTATATTTTGAAATCGAAAACAAACGACAG<br>ATTGTCGTTTTTGTGATCGTTATCTCGACATTTAAAAACAAATCATTTCATTATGTTTTGAAATCGAAAATAAACGACAG | -40.5 | N/A | 30 | N/A | 2177234 | 95 |
| *glgSp1* | (K-12)<br>EC042_3341 | TTCACTAAAAAGTGTGATCGGGGACAATATATTTACGCACGTTATGTTTAAAGGCACTACACTGATTGGGGAATACTGAAA<br>TTCACTAAAAAGTGTGATCGGGGACAATATATTTACGCACGTTATGTTTAAAGGCACTACACTGATTGGGGAATACTGAAA | -40.5 | N/A | 29 | N/A | 3192012 | 100 |
| *glpA* | (K-12)<br>EC042_2484 | AATGTTCAAAATGACGCATGAAATCACGTTTCACTTTCGAATTATGAGCGAATATGCGCGAAATCAAACAATTCATGTTTT<br>AATGTTCAGAATGACGCATGAAATCACGTTTCACTTTCGAATTATGAGCGAATATGCGCGAAATCAAACAATTCATGTTTT | -40.5 | N/A | 30 | N/A | 2352583 | 98.8 |
| *grcAp1* | (K-12)<br>EC042_2785 | TGTGTGGTTTTTATTGATTTAAATCAAAGATTCAAGGGTGTTTGAGGAGTATATATACACTCAAGCAACAATGGTTTTACC<br>TATGTGTTTTTTATTGATTTAAATCAAAGAATCAAGGGTGATTGAGGAGTATATATACACTCAAGCAACAATGGTTTTACC | -40.5 | N/A | 30 | N/A | 2716523 | 95.1 |
| *manX* | (K-12)<br>EC042_1982 | GAAAGTTAAATTACGGATCTTCATCACATAAAATAATTTTTTCGATATCTAAAATAAATCGCGAAACGCAGGGGTTTTTGG<br>GAAAGTTAAATTACGGATCTTCATCACATAAAATAATTTTTTCGATATCTAAAATAAATCGCGAAACGCAGGGGTTTTTGG | -40.5 | N/A | 30 | N/A | 1901933 | 100 |
| *mhpRp1* | (K-12)<br>EC042_0383 | ACTCGGACAAAATGTCGTTGCGCGCACAGTACAGCGCAACTTATTTTGTTAAAAACATGTAAATGATTTTTTATTGTGCGC<br>ACTCGGACAAAATGCCGTTGCGCGCACAGTACAGCGCAACTTATTTTGTTAAAAACATGTAAATGATTTTTTATTGTGCGC | -40.5 | N/A | 30 | N/A | 368536 | 98.8 |
| *nupC* | (K-12)<br>EC042_2611 | ACGTCATTATAGTGTGTGTCAGATCTCGTTTTCCTTAACCATGTTACATAGAATGTGCACGGAAATTTAACCTGCCTCATA<br>ACGTCATTATAGTGTGTGTCAGATCTCGTTTTCCTTAACCATGTTACATAGAATGTGCACGGAAATTTAACCTGCCTCATA | -40.5 | N/A | 29 | N/A | 2513009 | 100 |
| *nupG* | (K-12)<br>EC042_3171 | TTGCAATTATTTGCCACAGGTAACAAAAAACCAGTCCGCGAAGTTGATAGAATCCCATCATCTCGCACGGTCAAATGTGC<br>TTGCAATTATTTGCCACAGGTAACAAAAAACCAGTCCGCGAAGTTGATAGAATCCCATCATCTCGCACGGTCAAATGTGC | -40.5 | N/A | 29 | N/A | 3105651 | 96.2 |
| *ptsG* | (K-12)<br>EC042_1171 | CTGAAGTTGAAACGTGATAGCCGTCAAACAAATTGGCACTGAATTATTTTACTCTGTGTAATAAATAAAGGGCGCTTAGAT<br>CTGAAGTTGAAACGTGATAGCCGTCAAACAAATCGGCACTGAATTATTTTACTCTGTGTAATAAATAAAGGGCGCTTAGAT | -40.5 | N/A | 30 | N/A | 1157766 | 98.8 |
| *tsx* | (K-12)<br>EC042_0446 | AATGATAGAACTGTGAAACGAAACATATTTTTGTGAGCAATGATTTTTATAATAGGCTCCTCTGTATACGAAATATTTAG<br>AATGATAGAACTGTGAAACGAAACATATTTTTGTGAGCAATGATTTTTATAATAGGCTCCTCTGTATACGAAATATTTAG | -40.5 | N/A | 29 | N/A | 432091 | 100 |
| *bglGp2* | (K-12) | AAGTTAATAACTGCGAGCATGGTCATATTTTTATCAATAGCGCATTGCTATTTTCTCTGCACGCAATTAAATTAATTTCCG | -41.5 | N/A | 30 | N/A | 3906717 | |
| *caiF* | (K-12)<br>EC042_0036 | GATGACATAAGCAGGATTTAGCTCACACTTATCGACGGTGAAGTTGCATACTATCGATATATCCACAATTTTAATATGGCC<br>GATGACATAAGCAGGATTTAGCTCACACTTATCGACGGTGAAGTTGCATACTATCGATATATCCACAATTTTAATATGGCC | -41.5 | N/A | 30 | N/A | 34218 | 100 |
| *caiT* | (K-12)<br>EC042_0042 | CGAAACAAAAATGTGATACCAATCACAGAATACAGCTTATTGAATACCCATTATGAGTTAGCCATTAACGCGTCCACGAGG<br>CGAAACAAAAATGTGACACCAATCACAGAATACAGCTTATTGAATACCCATTATGAGTTAGCCATTAACGCGTCCACGAGG | -41.5 | N/A | 30 | N/A | 42037 | 98.8 |
| *galEp1* | (K-12)<br>EC042_0779 | GATTCCACTAATTTATTCCATGTCACACTTTTCGCATCTTTGTTATGCTATGGTTATTTCATACCATAAGCCTAATGGAGC<br>GATTCCACTAATTTATTCCATGTCACACTTTTCGCATCTTTGTTATGCTATGGTTATTTCATACCATAAGCCTAATGGAGC | -41.5 | N/A | 30 | N/A | 792081 | 100 |
| *glpT*<br>*glpT* | (K-12)<br>EC042_2483 | ATTTAATAATGTGTGCGGCAATTCACATTTAATTTATGAATGTTTTCTTAACATCGCGGCAACTCAAGAAACGGCAGGTTC<br>ATTTAATAATGTGTGCGGTAATTCACATTTAATTTATGAATGTTTTCTTAACATCGCGGCAACTCAAGAAACGGCAGGTTC | -41.5 | N/A | 30 | N/A | 2352451 | 98.8 |
| *gntP* | (K-12)<br>EC042_4826 | TGATCAGAGGATGTGACATTCATCGCAACAATGGTTGACCAATTTACATAACATATCGCCCAAAATAACAACGGTTCAACC<br>TGATCAGAGGATGTGACATTCATCGCAACAATGGTTGACCAATTTACATAACATGTCGCCCAAAATAACAACGGTTCAACC | -41.5 | N/A | 30 | N/A | 4551334 | 98.8 |
| *gntX* | (K-12)<br>EC042_3674 | CTAACCAAATGGCTTTATCAGGTGCCTGTTGCAGCACGGCTTCGGCCATATCAGCAAGTGACAGCGCACCAAATCCCCGGC<br>CTAACCAAATGGCTTTATCAGGTGCCTGTCGCAGCACGGCTTCGGCCATATCAGCAAGCGACAGCGCACCAAATCCCCGGC | -41.5 | N/A | 30 | N/A | 3544667 | 97.5 |
| *idnDp* | (K-12) | CAATTTTCTGACGTGATCTTCATCACAAATAATGACAGTTAAAACCGCTTAAATGCTTCCGAGGTGTCTACCTGACCAGTG | -41.5 | N/A | 31 | N/A | 4494435 | |

| | | | | | | | | |
|---|---|---|---|---|---|---|---|---|
| *malX* | (K-12)<br>EC042_1789 | TCGTTGCGTAATGTGATTTATGCCTCACTAAAATTTGATAAAACGTTTTATCTTCTCGCGCAATTTACTGAATCCAGATTG<br>TCGTTGCGTAATGTGATTTATGCCTCACTATAATTTGATAAAACGTTTTATCTTCTCGCGCAATTTACTGAATCCAGATTG | -41.5 | N/A | 30 | N/A | 1699313 | 98.8 |
| *melR* | (K-12)<br>EC042_4484 | AGGGTGAAAACCGTGCTCCCACTCGCAGTCATCCTCCCTCACTCCTGCCATAATTCTGATATTCCAGGAAAGAGAGCCATC<br>AGGGTGAAAACCGTGCTCCCACTCGCAGTCATCCTCCCTCACTCCTGCCATAATTCTGATATTCCAGGAAAGAGAGCCATC | -41.5 | N/A | 30 | N/A | 4341650 | 100 |
| *mglB* | (K-12)<br>EC042_2383 | CGCTTTCAATCTGTGAGTGATTTCACAGTATCTTAACAATGTGATAGCTATGATTGCACCGTTTTAACGTTGTAACCCGTA<br>CGCTTTCAATCTGTGAGTGATTTCACAGTATCTTAACAATGTGATAGCTATGATTGCACCGTTTTAACGTTGTAACCCGTA | -41.5 | N/A | 30 | N/A | 2240566 | 100 |
| *nirB* | (K-12)<br>EC042_3627 | ATAAAGGTGAATTTGATTTACATCAATAAGCGGGGTTGCTGAATCGTTAAGGTAGGCGGTAATAGAAAAGAAATCGAGGCA<br>ATAAAGGTGAATTTGATTTACATCAATAAGCGGGGTTGCTGAATCGTTAAGGTAGGCGGTAATAGAAAAGAAATCGAGGCA | -41.5 | N/A | 29 | N/A | 3493987 | 100 |
| *pkap* | (K-12)<br>EC042_2790 | AAAACGCAACATCTGGGTAGCATCACAGCAGAACAGTTAGAAAGCGTTTAAAATCATTCGGTCACTTCTGCGGGAGACCGG<br>AAAACGCAACATCTGGGTAGCATCACAGCAGAACAGTTAGAAAGCGTTTAAAATCATTCGGTCACTTCTGCGGGAGACCGG | -41.5 | N/A | 30 | N/A | 2719931 | |
| *rpoHp5*<br>*rpoH* | (K-12)<br>EC042_3722 | GCATTGAACTTGTGGATAAAATCACGGTCTGATAAAACAGTGAATGATAACCTCGTTGCTCTTAAGCTCTGGCACAGTTG<br>GCATTGAACTTGTGGATAAAATCACGGTCTGATAAAACAGTGAATGATAACCTCGTTGCTCTTAAGCTCTGGCACAGTTG | -41.5 | N/A | 30 | N/A | 3600849 | 100 |
| *srlA* | (K-12)<br>EC042_2895 | ATCTTTCATTTTGCGATCAAAATAACACTTTTAAATCTTTCAATCTGATTAGATTAGGTTGCCGTTTGGTAATAAAACAAT<br>ATCTTTCATTTTGCGATCAAAATAACACTTTTAAATCTTTCAATCTGATTAGATTAGGTTGCCGTTTGGTAATAAAACAAT | -41.5 | N/A | 31 | N/A | 2825791 | 100 |
| *udp* | (K-12)<br>EC042_4211 | ATTTGCGTCATGGTGATGAGTATCACGAAAAAATGTTAAACCCTTCGGTAAAGTGTCTTTTTGCTTCTTCTGACTAAACCG<br>ATTTTCGTCATGGTGATGGGTATCACGAAAAAATGTTAAACCCTTCGGTAAAGTGTCTTTTTGCTTCTTCTGACTAAACCG | -41.5 | N/A | 30 | 97.5 | 4016391 | 97.5 |
| *ychH* | (K-12)<br>EC042_1262 | AGGGTTGTAATTGTGATCACGCCCGCACATAACCCACTGGGTGTTGTCTATACTTTACACATAAGGAAGAGGGGTATTCCC<br>AGGGTTGTAATTGTGATCACGCCCGCACATAACCCACAGGGTGTTGTCTATACTTTACACATAAGGAAGAGGGGTATTCCC | -41.5 | N/A | 30 | N/A | 1258738 | 98.8 |
| *ykgR* | (K-12) | CTATTCAAATTTGTGATACATGTCAAAAAATTGGTTGACCAAACTCGTTATTTATATAAGGGCACTTACGAAGTGCACTCT | -41.5 | N/A | 30 | N/A | 313285 | |
| *araF* | (K-12)<br>EC042_2064 | AATTCTGCGATGTGATATTGCTCTCCTATGGAGAATTAATTTCTCGCTAAAACTATGTCAACACAGTCACTTATCTTTTAG<br>AATTCTGCGATGTGATATTGCTCTCCTATGGAGAATTAATTTCTCGCTAAAACTATGTCAACACGGTCACTTATCTTTTAG | -42.5 | N/A | 30 | N/A | 1986238 | 98.8 |
| *cdd* | (K-12)<br>EC042_2375 | GCATAATTAATGAGATTCAGATCACATATAAAGCCACAACGGGTTCGTAAACTGTTATCCCATTACATGATTATGAGGCAA<br>GCATAATTAATGAGATTCAGATCACATATAAAGCCATAACGGGTTCGTAAACTGTTATCCCATTACATGATTATGAGGCAA | -42.5 | N/A | 30 | N/A | 2231819 | 98.8 |
| *csgDp1* | (K-12)<br>EC042_1105 | AGTTACATTTAGTTACATGTTTAACACTTGATTTAAGATTTGTAATGGCTAGATTGAAATCAGATGTAATCCATTAGTTTT<br>AGTTACATTTAGTTACATGTTTAACACTTGATTTAAGATTTGTAATGGCTAGATTGAAATCAGATGTAATCCATTAGTTTT | -42.5 | N/A | 32 | N/A | 1103344 | 100 |
| *csiE* | (K-12)<br>EC042_2739 | CTTGCCAACATTTCTGATGATTAGCATTCCCTTCGCCATTTCCTTGAGCAAACTTTAGCTATTCTTATCAATTATGCTTAT<br>CTTGCCAACATTTCTGATGATTAGCATTCCCTTCGCCATTTCCCTGAGCAAACTTTAGCTATTCTTATCAATTATGCTTAT | -42.5 | N/A | 31 | N/A | 2665401 | 98.8 |
| *cyaR* | (K-12) | TGGAAAATTCTTAGAAACCGATCACATACAGCTGCATTTATTAAGGTTATCATCCGTTTCGCTGAAAAACATAACCCATAA | -42.5 | N/A | 30 | N/A | 2167114 | |
| *fepA* | (K-12)<br>EC042_0621 | CCATGTTTACTGTGCAATTTTTCATTGATTGCAGAAATATATTGATAATATTATTGATAACTATTTGCATTTGCAATAGCG<br>CCATGTTTACTGTGCAATTTTTCATTGATTGCAGAAATATATTGATAATATTATTGATAACTATTTGCATTTGCAATAGCG | -42.5 | N/A | 31 | N/A | 612667 | 100 |
| *galP* | (K-12)<br>EC042_3150 | ATTACACTGATGTGATTTGCTTCACATCTTTTTACGTCGTACTCACCTATCTTAATTCACAATAAAAAATAACCATATTGG<br>ATTACACTGATGTGATTTGCTTCACATCTTTTTACGTCGTACTCACCTATCTTAATTCACAATAAAAAATAACCATATTGG | -42.5 | N/A | 30 | N/A | 3088255 | 100 |
| *galS* | (K-12)<br>EC042_2384 | TTTCCATTGCTGTGACTCGATTCACGAAGTCCTGTATTCAGTGCTGACAAAATAGCCGCCAGCAAGCAGTCATTTACTGCA<br>TTTCCATTGCTGTGACTCGATTCACGAAGTCCTGTATTCAGTGCTGACAAAATAGCCGCCAGCAAGCAGTCATTTACTGCA | -42.5 | N/A | 30 | N/A | 2241710 | 100 |
| *ompW* | (K-12)<br>EC042_1310 | TCATATGTAATGTGATCTATGTAGGATCATTTGTTACTCCAATGTAGGTATATTCGTCACGTTTTTATAACCATAACGACG<br>TCATATGTAATGTGATCTATGTAGGATCATTTGTTACTCCAATGTAGGTATATTCGTCACGTTTTTATAACCATAACGACG | -42.5 | N/A | 31 | N/A | 1313991 | 100 |
| *ppiAp2* | (K-12)<br>EC042_3625 | CATTTTAAGAGGTGATTTTGATCACGGAATAAAAAGTGATCGTCAGGTTACATATATTTCAGATACGTAAAATTAGGTAAA<br>CAATATTTAAAGTGATTCTCATCACGAAATATACGTGATCATCAGGTTACATATATTTCAAGTGCGTAAAATTAGGTAAA | -42.5 | N/A | 31 | N/A | 3492395 | 80.2 |
| *ptsHp1* | (K-12)<br>EC042_2624 | GAATCGATTTTATGATTTGGTTCAATTCTTCCTTTAGCGGCATAATGTTTAATGACGTACGAAACGTCAGCGGTCAACACC<br>GAATCGATTTTATGATTTGGTTCAATTCTTCCTTTAGCGGCATAATGTTTAATGACGTACGAAACGTCAGCGGTCAACACC | -42.5 | N/A | 32 | N/A | 2533608 | 100 |
| *yeiW* | (K-12) | CTTTCGGGCAAGTGGTATTCGCACTTTTGCTGGTGCAAAAAAGGTGGTACTGTGCGCGCTCATCAATCCGGTGGTTAACCT | -42.5 | N/A | 30 | N/A | 2265330 | |
| *yhfA* | (K-12)<br>EC042_3618 | GCACGGTAATGTGACGTCCTTTGCATACATGCAGTACATCAATGTATTACTGTAGCATCCTGACTGTTTTAGCATAGCTTT<br>GCACGGTAATGTGACGTCCTTTGCATACATGCAGTACATCAATGTATTACTGTAGCATCCTGACTGTTTTAGCATAGCTTT | -42.5 | N/A | 30 | N/A | 3485951 | 100 |
| *cpdB* | (K-12)<br>EC042_4694 | TGCGCCAACTGTGATAGTGTCATCATTTTCAAAGCGTAAAATTGTGGCATTCTTCACTGTTCTATAAGTAAGACGTTTATT<br>TGCGCCAACTGTGATAGTGTCATCATTTTCAAAGCGTAAAATTGTGGCATTCTTCACTGTTCTATAAGTAAGACGTTTATT | -43.5 | N/A | 31 | N/A | 4436629 | 100 |
| *dadAp1*<br>*dadA* | (K-12)<br>EC042_1238 | TCAGGGAGATGTGAGCCAGCTCACCATAAAAAAGCCGCATGTTGAATAATATTTTCAACTGAGTTATCAAGATGTGATTAG<br>TCAGGGAGATGTGAGCCAGCTCACCATAAAAAAGCCGTATGTTGAATAATATTTTCAACTGAGTTATCAAGATGTAATTAG | -43.5 | N/A | 30 | N/A | 1237509 | 97.5 |
| *malI* | (K-12)<br>EC042_1788 | AAATTTTAGTGAGGCATAAATCACATTACGCAACGATAATAGCGGGTATAAGATAAATAAAAGGTAAAACGTTTTATCTGT<br>AAATTATAGTGAGGCATAAATCACATTACGCAACGATATTAGCGGGTATAAGATAAATAAAAGGTAAAACGTTTTATCTGT | -43.5 | N/A | 30 | N/A | 1699228 | 97.5 |
| *tdcA* | (K-12)<br>EC042_3410 | AAGTTAATTTGTGAGTGGTCGCACATATCCTGTTCATTTCATTTTGATACACTTCATGCCGTCAATGAGGTAATTAACGTA<br>AAGTTAATTTGTGAGTGGTCGCACATATCCTGTTCATTTCATTTTGATACACTTCATGCCGTCAATGAGGTAATTAACGTA | -43.5 | N/A | 31 | N/A | 3267093 | 100 |
| *yeiP* | (K-12)<br>EC042_2404 | ACCAGCAAAAGTGCGAATACCACTTGCCCGAAAGGCCCGTCGCGAGTACTTTGTCGCGATATTTTTGACATTTTCGACTAC<br>ACCAGCAAAAGTGCGAATACCACTTGCCCGAAAGGCCCGTCGCGAGTACTTTGTCGCGATATTTTTGACATTTTCGACTAC | -44.5 | N/A | 31 | N/A | 2265417 | 100 |
| *aldA* | (K-12)<br>EC042_1546 | TTACACTTGTTTTTATGAAGCCCTTCACAGAATTGTCCTTTCACGATTCCGTCTCTCTGATGATTGATGTTAATTAACAATGTATTCACCGAAAACAAACA<br>TTACACTTGTTTTTATGAAGCCCTTCACAGAATTGTCCTTTCACGATGCCGTCTCTATGATGATTGATGTTAATTAACAATGCATTTACCAATAACAAACA | -59.5 | 26 | 50 | 18 | 1488190 | 94.1 |

| | | | | | | | | |
|---|---|---|---|---|---|---|---|---|
| crp | (K-12)<br>EC042_3620 | CGCGCAACGGAAGGCGACCTGGGTCATGCTGAAGCGAGACACCAGGAGACACAAAGCGAAAGCTATGCTAAAACAGTCAGGATGCTACAGTAATACATTGA<br>CGCGCAACGGAAGGCGACCTGGGTCATGCTGAAGCGAGACACCAGGAGACACAAAGCGAAAGCTATGCTAAAACAGTCAGGATGCTACAGTAATACATTGA | -59.5 | 25 | 48 | 17 | 3485953 | 100 |
| dadAp2 | (K-12) | AGAGTCAGGGAGATGTGAGCCAGCTCACCATAAAAAAGCCGCATGTTGAATAATATTTTCAACTGAGTTATCAAGATGTGATTAGATTATTATTCTTTTAC | -59.5 | 25 | 48 | 17 | 1237525 | |
| fadD | (K-12)<br>EC042_1970 | GATAAAAATAAATAGTGACGCGCTTCGCAACCTTTTCGTTGGGTAATTATCAAGCTGGTATGATGAGTTAATATTATGTTAACGGCATGTATATCATTTGG<br>GATAAAAATAAATAGTGACGCGCTTCGCAACCTTTTCGTTGGGTAATTATCAAGCTGGTATGATGAGTTAATATTATGTTAACGGCATGTATATCATTTGG | -59.5 | 26 | 48 | 16 | 1889807 | 100 |
| tnaC | (K-12) | CTCCCCGAACGATTGTGATTCGATTCACATTTAAACAATTTCAGAATAGACAAAAACTCTGAGTGTAATAATGTAGCCTCGTGTCTTGCGAGGATAAGTGC | -59.5 | 26 | 48 | 16 | 3888411 | |
| aldB | (K-12)<br>EC042_3898 | CTTGCCCGTAAATTCGTGATAGCTGTCGTAAAGCTGTTACCGACTGGCGAAGATTTCGCCAGTCACGTCTACCCTTGTTATACCTCACACCGCAAGGAGACG<br>CTTGCCCGTAAATTCGTGATAGCCGTCGTAAAGCTGTTACCGACTGGCGAAGATTTCGCCAGTCACGTCTACCCTTGTTATACCTCACACCGCAAGGAGACG | -60.5 | 26 | 49 | 17 | 3756535 | 99 |
| glpF | (K-12)<br>EC042_4301 | ATAACGCTCATTTTATGACGAGGCACACACATTTTAAGTTCGATATTTCTCGTTTTTGCTCGTTAACGATAAGTTTACAGCATGCCTACAAGCATCGTGGAG<br>ATAACGCTCATTTTATGACGAAGCACACACATTTTAAGTTCGATATTTCTCGTTTTTGCTCGTTAACGATAAGTTTACAGCATGCCTACAAGCATCGTGGAG | -60.5 | 26 | 49 | 17 | 4118161 | 99 |
| kbaZ | (K-12)<br>EC042_3423 | ATTATGTTTCTTTTGTGAATCAGATCAGAAAACCATTATCTTTCGTTTTATTTTTATCTCACCATGACGCAGTATCAACTGAAACAAAACGAAAGATTAATA<br>ATTATGTTTCTTTTGTGAATCAGATCAGAAAACTATTATCTTTCGTTTTATTTTTATCTCACCATGGCGCAGTATCAACTGAAACAAAACGAAAGATTAATA | -60.5 | 26 | 49 | 17 | 3278866 | 98 |
| lsrR | (K-12)<br>EC042_0152 | CGTAGAGTCAAACTGTGGTTGCCATCACAGATATAAATGAGCAAGAACTGAACAATTGCATTAAAGATTTAAATATGTTCAAAGTGAAGAATGAATTATGAC<br>CGTAGAGTCAAACTGTGGTTGCTATCACAGATATAAATGCGCAAGAACTGAACAATTGCATTAAAGATTTAAATATGTTCAAAGTGAAGAATGAATTATGAC | -60.5 | 27 | 49 | 16 | 1601257 | 98 |
| mdhp1 | (K-12)<br>EC042_3521 | ATAAACGCTAAACTTGCGTGACTACACATTCTTGAGATGTGGTCATTGTAAACGGCAATTTTGTGGATTAAGGTCGCGGCAGCGGAGCAACATATCTTAGTT<br>ATAAACGCTAAACTTGCGTGACTACACATTCTTGAGATGTGGTCATTGTAAACGGCAATTTTGTGGATTAAGGTCGCGGCAGCGGAGCAACATATCTTAGTT | -60.5 | 25 | 48 | 17 | 3384313 | 100 |
| metK | (K-12)<br>EC042_3149 | GTTTTTAACAGAATGAGACACGATTCAAAAAAAAGTGGAAATAGGGTGAAGAATTGACCTAAAATAGCCATCCAGATGTTAATCCATCCATACCGATTAACA<br>GTTTTTAACAGAATGAGACACGATTCAAAAAAAAGTGGAAATAGGGTGAAGAATTGACCTAAAATAGCCATCCAGATGTTAATCCATCCATACCGATTAACA | -60.5 | 26 | 48 | 16 | 3086577 | 100 |
| nanA | (K-12)<br>EC042_3509 | ATAATGCCACTTTAGTGAAGCAGATCGCATTATAAGCTTTCTGTATGGGGTGTTGCTTAATTGATCTGGTATAACAGGTATAAAGGTATATCGTTTATCAGA<br>ATAATGCCACTTTAGTGAAGCAGATCGCATTATAAGCTTTCTGTATGGGGTGTTGCTTAATTGATCTGGTATAACAGGTATAAAGGTATATCGTTTATCAGA | -60.5 | 24 | 49 | 19 | 3373620 | 100 |
| ppiAp3 | (K-12) | TTTTAGGTGATTTTGTGATCTGTTTAAATGTTTTATTGCAATCGGTTGCTAAATTGCATTTTAAGAGGTGATTTTGATCACGGAATAAAAAGTGATCGTCAG | -60.5 | 25 | 49 | 18 | 3492430 | |
| ptsHp5 | (K-12) | TTAGGTGCTTTTTTGTGGCCTGCTTCAAACTTTCGCCCCTCCTGGCATTGATTCAGCCTGTCGGAACTGGTATTTAACCAGACTAATTATTTTGATGCGCGA | -60.5 | 27 | 50 | 17 | 2533498 | |
| relA | (K-12)<br>EC042_2981 | GATCGCGAAAAACTGGAACGCTTTTCGCATTCTGAAGGCCTGGATCTGTATCTCGCCCCCGATAGTGAGATACTCGAAACCGTCTCTGGTGAGATGCCCTGG<br>GATCGCGAAAAACTGGAACGCTTTTCGCATTCTGAAGGCCTGGATCTGTATCTCGCCCCCGATAGTGAGATACTCGAAACCGTCTCTGGTGAGATGCCCTGG | -60.5 | 25 | 50 | 19 | 2914277 | 100 |
| treB | (K-12)<br>EC042_4718 | CTGAACGATAAATTGTGATCTTCGCTGCGTTTCGGGAACGTTCCCGTTTTTAAATTTTTCCGCGCAATATATTCTGCAGCCAACCAAAAATGTCATCTGCCA<br>CTGAACGATAAATTGTGATCTTCGCTGCGTTTCGGGAACGTTCCCGTTTTTAAATTTTTCCGCGCAATATATTCTGCAGCCAACCAAAAATGTCATCTGCCA | -60.5<br>**(-61.5)** | 26<br>**(27)** | 49<br>**(50)** | 17 | 4466210<br>**(+1)** | 99 |
| ulaA | (K-12)<br>EC042_4674 | ATGTGGAATTATTTGCGGGTCGCGTCACATTTAATCATAAATAATCTTGTTGTGATTACTTTTGAAAATTAGAGTGAGTGCACAACATTCCGGGTGTGTGGA<br>ATGTGGAATTATTTGCGGGTCGCGTCACATTTAATCATAAATAATCTTGTTGTGATTACTTTTGAAAATTAGAGTGAGTGCACAACATTCCGGGTGTGTGGA | -60.5 | 26 | 49 | 17 | 4419927 | 100 |
| bglGp | (K-12) | CAAAGTTAATAACTGCGAGCATGGTCATATTTTTATCAATAGCGCATTGCTATTTTCTCTGCACGCAATTAAATTAATTTCCGAACCTGGATGTTCGTTATAA | -61.5 | 26 | 49 | 17 | 3906697 | |
| cspEp2 | (K-12)<br>EC042_0659 | ACATCAAAAATAACGCGACTTTTATCACTTTTTAGTAAAGTTACACTGGACAAAGCGTACCACAATTGGTGTACTGGTAACCGACACAGCATTTGTGTCTATT<br>ACATCAAAAATAACGCGACTTTTATCACTTTTTAGTAAAGTTGCACTGGACAAAGCGTACCACAATTGGTGTACTGGTAACCGACACAGCATTTGTGTCTATT | -61.5 | 26 | 49 | 17 | 657250 | 99 |
| fadH | (K-12)<br>EC042_3374 | CCGGGCGTATTCTTTTTGAATCCCATCACAAACCCCGCACTCCCCTTTTCCCTTTTCTCCGGCGACGGCTAAATTAGAACTCATCCGACCACATAACAATTAT<br>ACAGGCGTATTCTTTTTGAATCCCATCACAAACCCCACACTCCCCTTTTCCCTTTTCTCCGGCGACGGCTAAATTAGAACTCATCCGACCACATAACAATTAT | -61.5 | 26 | 49 | 17 | 3231624 | 97.1 |
| lacZp1 | (K-12) | GCAACGCAATTAATGTGAGTTAGCTCACTCATTAGGCACCCCAGGCTTTACACTTTATGCTTCCGGCTCGTATGTTGTGTGGAATTGTGAGCGGATAACAATT | -61.5 | 26 | 50 | 18 | 366343 | |
| nagE | (K-12)<br>EC042_0707 | AATAACGATATTTGGTGACAAAACTCACAAAAGACACGCGTTTAATTTGCGATACGAATTAAATTTTCACACACTCTGTAGCAGATGATCTAACAATCTGATT<br>GATAACGATATTTGGTGACAAAACTCACAAAAGACACGCGTTTAATTTGCGATACGAATTAAATTTTCACACACTCTGTAGCAGATGATCTAACAATCTGATT | -61.5 | 26 | 49 | 17 | 703840 | 99 |
| nanC | (K-12)<br>EC042_4822 | GATCGACATATTTTGTGACACGAATTGCAAATCTGGTTTTGTTGTATGGATTGCGTGATTTTTGATCTGGTATAACAGGTATAAAGGTGCACCAAGATAGTCA<br>GATCGACATATTTTGTGACACGAATTGCAAATCTGGTTTTGTTGTATGGATTGCGTGATTTTTGATCTGGTATAACAGGTATAAAGGTGCACCGCGATAGACA | -61.5 | 26 | 50 | 18 | 4539928 | 97.1 |
| secB | (K-12)<br>EC042_3919 | TTCCCCAGATTTTTATTGACGCACAGCACATTGGCGGCTGTGATGACTTGTATGCATTGGATGCACGTGGTGGACTGGATCCCCTGCTGAAATAACGTGTGAA<br>TTCCCCAGATTTTTATTGACGCACAGCACATTGGCGGCTGTGATGACTTGTATGCATTGGATGCACGTGGTGGACTGGATCCCCTGCTGAAATAACGTGTGAA | -61.5 | 27 | 50 | 17 | 3784203 | 100 |
| gadA | (K-12)<br>EC042_3812 | CGTTTTTCTGCTTAGGATTTTGTTATTTAAATTAAGCCTGTAATGCCTTGCTTCCATTGCGGATAAATCCTACTTTTTTATTGCCTTCAAATAAATTTAAGGAG<br>CGTTTTTCTGCTTAGGATTTTGTTATTTAAATTAAGCCTGTAATGCCTTGCTTCCATTGCGGATAAATCCTACTTTTTTATTGCCTTCAAATAAATTTAAGGAG | -62.5 | 27 | 50 | 17 | 3667607 | 100 |
| modAp | (K-12)<br>EC042_0782 | TGCCGATATTTTTTCTTATCTACCTCACAAAGGTTAGCAATAACTGCTGGGAAAATTCCGAGTTAGTCGTTATATTGTCGCCTACATAACGTTACATTAAGGGG<br>TGCCGATATTTTTTCTCATCTACCTCACAAAGGTTAGCAATAACTGCTGGGAAAATTCTGAGTTAGCCGTTATATTGACGCCTACATAACGTTACATTAAGGGG | -62.5 | 27 | 50 | 17 | 795062 | 96.2 |
| rbsD | (K-12)<br>EC042_4135 | CCATGTAAAACGTTTCGAGGTTGATCACATTTCCGTAACGTCACGATGGTTTTCCCAACTCAGTCAGGATTAAACTGTGGGTCAGCGAAACGTTTCGCTGATGG<br>CCATGTAAAACGTTTCGAGGTTGATCACATTTCCGTAACGTCACGATGGTTTTCCCAACTCAGTCAGGATTAAACTGCGGGTCAGCGAAACGTTTCGCTGATGG | -62.5 | 29 | 50 | 15 | 3933322 | 99 |
| rpoS | (K-12)<br>EC042_2935 | GCAGAGCAAGGAGTTGTGATCAAGCCTGCACAAAATTCCACCGTTGCTGTTGCGTCGCAACCGACAATTACGTATTCTGAGTCTTCGGGTGAACAGAGTGCTAA<br>GCAGAGCAAGGAGTTGTGATCAAGCCTGCACAAAATTCCACCGTTGCTGTTGCGTCGCAACCGACAATTACGTATTCTGAGTCTTCGGGTGAACAGAGTGCTAA | -62.5 | 29 | 52 | 17 | 2868118 | 100 |
| aaeX | (K-12)<br>EC042_3527 | ACTCTGACTTAAAAGTGATTTAGATCACATAATATAGATAACAGCATAACAGTTACGCTAATATATTAAATATCAATCTACAGCAATGTTGCTCTCGCCCGGCTT<br>ACTCTGACTTAAAAGTGATTTAGATCACATAATATAGATAACAGCATAACAGTTACGCTAATATATTAAATATCAATCTACAGCAATGTTGCTCTCGCCCGGCTT | -63.5 | 32 | 50 | 12 | 3389430 | 100 |
| dusB | (K-12)<br>EC042_3543 | TTTAAATGCAATTCTTTGATCCATCTCAGAGGATTGGTCAAAGTTTGGCCTTTCATCTCGTGCAAAAAATGCGTAATATACGCCGCCTTGCAGTCACAGTATGGT<br>TTTAAATGCAATTCTTTGATCCATCTCAGAGGATTGGTCAAAGTTTGGCCTTTCATCTCGTGCAAAAAATGCGTAATATACGTTGCAGTCACAGTATGGT | -63.5 | 30 | 53 | 17 | 3410246 | 95.2 |
| gadB | (K-12) | ATAAATAACATTAGGATTTTGTTATTTAAACACGAGTCCTTTGCACTTGCTTACTTTATCGATAAATCCTACTTTTTTAATGCGATCCAATCATTTTAAGGAG | -63.5 | 32 | 51 | 13 | 1572072 | |
| glpD | (K-12)<br>EC042_3686 | ATAAACGCCATAATGTTATACATATCACTCTAAAATGTTTTTTCAATGTTACCTAAAGCGCGATTCTTTGCTAATATGTTCGATAACGAACATTTATGAGCTTTA<br>ATAAACGCCATAATGTTATACATATCACTCTAAAATGTTTTTTCAATGTTACCTAAAGCGCGATTCTTTGCTAATATGTTCGATAACGAACATTTGTGAGCTTTA | -63.5 | 32 | 51 | 13 | 3561971 | 99 |

| | | | | | | | | |
|---|---|---|---|---|---|---|---|---|
| ivbL | (K-12)<br>EC042_4026A | ATTCAGTACAAAACGTGATCAACCCCTCAATTTTCCCTTGCTGAAAAATTTTCCATTGTCTCCCCTGTAAAGCTGTGCTTGTATAAATATTGTTAAACACAAAAC<br>GTACAGTACAAAACGTGATCAACCCCTCAATTTTCCCTTGCTGAAAAATTTTCCATTGTCTCCCCTGTAAAGCTGTGCTTGTATAAATATTGTTAAACACAAAAC | -63.5 | 30 | 47 | 11 | 3853023 | 98.1 |
| ldtB | (K-12)<br>EC042_0908 | CAAAGGCAAAAAATGTGATTTCGTACACATCTGATTTCACTGTGAGCTGGAATGAACTTATAATGCGCTTCCAATACTCTAATATTCTCAACCCAATGGCCTGCC<br>CAAAGGCAAAAAATGTGATTTCGTACACATCTGATTTCACTGTGAGCTGGAATGAACTTATAATGCGCTTCCAATACTCTAATATTCTCAACCCAATGGCCTGCC | -63.5 | 32 | 51 | 13 | 855786 | 100 |
| ptsHp4 | (K-12) | TTAGGTGCTTTTTTGTGGCCTGCTTCAAACTTTCGCCCCTCCTGGCATTGATTCAGCCTGTCGGAACTGGTATTTAACCAGACTAATTATTTTGATGCGCGAAAT | -63.5 | 31 | 50 | 13 | 2533501 | |
| cytR | (K-12)<br>EC042_4308 | GCGGATCGAAAAATTCAATATTCATCACACTTTTCATGAAAATTCTGTAACCGTTTTCACGCGCTATCTGCTAAAAATGTTGCCGATGTGAAGTAAACATGGATGT<br>GCGGATCGAAAAATTCAATATTCATCACACTTTTCATGAAAATTCTGTAACCGTTTTCACGCGCTATCTGCTAAAAATGTTGCCGATGTGAAGTAAACATGGAAGT | -64.5 | 34 | 51 | 11 | 4124509 | 99.1 |
| gltB | (K-12)<br>EC042_3502 | TTTAATAAAGAATTTTGCGCTAAAGCACATTTCTGTACCAATAAGCTTGCCATTTGACCTGTATCAGCTTTCCCGATAAGTTGGAAATCCGCTGGAAGCTTTCTGGA<br>TTTAATAAAGAATTTTGCGCTAAAGCACATTTCTGTACCAATAAGCTTGCCATTTGACCTGTATCAGCTTTCCCGATAAGTTGGAAATCCGCTGGAAGCTTTCTGGA | -65.5 | 33 | 54 | 15 | 3354509 | 100 |
| hpt | (K-12)<br>EC042_0124 | AGGTTTTAACATGTGTGATCGTCATCACAATTCGAGCTTTATTAACAGATTCCGCGAATGAATAGTTTTACTGGTATACTGCGTGTCTTGCGCTTTGTTGCGGTGCC<br>AGGTTTTAACATGTGTGATCGTCATCACAATTCGAGCTTTATTAACAGATTCCGCGAATGAATAGTTTTACTGGTATACTGCGTGTCTTGCGCTTTGTTGCGGTGCC | -65.5 | 29 | 54 | 19 | 141360 | 100 |
| sfsA | (K-12)<br>EC042_0146 | ACTGGTCGTATGCGGTGACGGAGTTCACCCTTTACGCCTCCTCGTTTGCCCGTGGACGCACACGCTACACGCCGCTAAAACGCTGGGCGCTAACGCAATAACAAGGA<br>ACTGGTCGTATACGGTGACGGAGTTCACCCTTTACGCCTCCTCGTTTGCCCGTGGACGCACACGCTACACGCCGCTAAAACGCTGGGCGCTAACGCAATAACAAGGA | -65.5 | 32 | 55 | 17 | 161515 | 99.1 |
| csiD | (K-12)<br>EC042_2856 | ATGTTACTAATTTGTTGCTTTTGATCACAATAAGAAAACAATATGTCGCTTTTGTGCGCATTTTTCAGAAATGTAGATATTTTTAGATTATGGCTACGAAATGAGCATCG<br>ATGTTACCAATTTGTTGCTTTTGATCACAATAAGAAAACAATATGGCGCATTTGTGCGCATTTTTCAGAAATGTAGATATTTTTAGATTATGGCTACGAAATGAGCATCG | -68.5 | 31 | 57 | 20 | 2788927 | 97.3 |
| gntK | (K-12)<br>EC042_3703 | TTAAGCCCACAAATTTGAAGTAGCTCACACTTATACACTTAAGGCATGGATGGATATTGCTTCTGATATTGTCCGGCTGGACAATGTTACCGATAACAGTTACCCGTAAC<br>TTAAGCCCACAAATTTGAAGTAGCTCACACTTATACACTTAAGGCACGGATGGATATTGCTTCTGATATTGTCCGGCTGGACAATGTTACCGATAACAGTTACCCGTAAC | -68.5 | 36 | 59 | 17 | 3577645 | 99.1 |
| prpR | (K-12)<br>EC042_0366 | TAATCCGCAAATATGCGTTTCAGTTAACGTTTCAGGCAATGTTTCACCGCGTTTCATTGCAACAATTATGAAACAAGACTAAACCCAATATTCGGTTTCTTAACTTTGCG<br>TAATCCGCAAATATGCGTTTCAGTTAACGTTTCAGGCAATGTTTCACCGCGTTTCATTGCAACAATTATGAAACAAGACTAAACCCAATATTCAGTTTCTTAACTTTGCG | -68.5 | 36 | 59 | 17 | 348471 | 99.1 |
| acs | (K-12)<br>EC042_4440 | AATTCTACTTTTGCGTGATCTGTCGCCCAAATACTAAACAAAACTGCCAATACCCCTACATTTAACGCTTATGCCACATATTATTAACATCCTACAAGGAGAACAAAAGCA<br>AATTCTACTTTTGCGTGATCTGTCGCCCAAATACTAAACAAAACTGCCAATACCCCTACATTTAACGCTTATGCCACATATTATTAACATCCTACAAGGAGAACAAAAGCA | -69.5 | 36 | 58 | 16 | 4287391 | 100 |
| fixA | (K-12)<br>EC042_0043 | TGTTTTCAATATTGGTGAGGAACTTAACAATATTGAAAGTTGGATTTATCTGCGTGTGACATTTTCAATATTGGTGATTAAAGTTTTATTTCAAAATTAAAGGGCGTGATA<br>TGTTTTCAATATTGGTGAGGAACTTAACAATATTGAAAGTTGGATTTATCTGCGTGTGACATTTTCAATATTGGTGATTAAAGTTTTATTTCCAAATTAAAGGGCGTGATA | -69.5 | 35 | 58 | 17 | 42325 | 99.1 |
| chiP | (K-12)<br>EC042_0709 | CATCAAAATATGTGCGGTTGCTCATGTTCTTACATTCTGGTTACAGAAAGAGATTGATAATTCGCGTCGCGAAAAATAGTCTGTTCCTGTAGTCAGCGAGACTTTTCTC<br>CATCAAAATATGTGCGGTTGCTCATGTTCTTACATTCTGGTTACAGAAAGAGATTGATAATTCGCGTCGCGAAAAATAGTCTGTTCCTGTAGTCAGCGAGACTTTTCTC | -70.5 | 36 | 59 | 17 | 708236 | 100 |
| epd | (K-12)<br>EC042_3136 | AAGGGGATAAAAGTGTGATGTGAGTCAGATAAATGTCTTCTTCGGCTGGACAAACATTCCTTTTATTCCACGTTTCGCTTATCCTAGCTGAAGCGTTTCAGTCGATTAAATG<br>AAGGGGATAAAAGTGTGATGTGAGTCAGATAAATGTCTTCTTCGGCTGGACAAACATTCCTTTTATTCCACGTTTCGCTTATCCTAGCTGAAGCGTTTCAGTCGATTAAATG | -70.5 | 36 | 59 | 17 | 3073823 | 100 |
| furpb | (K-12)<br>EC042_0711 | TTGCCGTTGTAAATGTAAGCTGTGCCACGTTTTTATTAACAATATTTGCCAGGGACTTGTGGTTTTCATTTAGGCGTGGCAATTCTATAATGATACGCATTATCTCAAGAGC<br>TTGCCGTTGTAAATGTAAGCAGTGCCACGTTTTTATTAACAATATTTGCCAGGGACTTGTGGTTTTCATTTAGGCGTGGCAATTCTATAATGATACGCATTATCTCAAGAGC | -70.5 | 36 | 59 | 17 | 710723 | 99.1 |
| idnK | (K-12) | ATGTTACGCATAACGTGATGTGCCTTGTAATTCTTATCAGTAGAAATAAAAAAACGTGAAATTATTATGCCGCCAGGCGTAGTATCGCAGCAGGTAAGATGATTCAGGAGAT | -70.5 | 36 | 59 | 17 | 4494597 | |
| malT | (K-12)<br>EC042_3679 | AAAGATTTGGAATTGTGACACAGTGCAAATTCAGACACATAAAAAAACGTCATCGCTTGCATTAGAAAGGTTTCTGGCCGACCTTATAACCATTAATTACGAAGCGCAAAAA<br>AAAGATTTGGAATTGTGACAGAGTGCAAATTCAGACACATAAAAAAACGTCATCGCTTGCATTAGAAAGGTTTCTGGCCGACCTTATAACCATTAATTACGAAGCGCAAAAA | -70.5 | 36 | 59 | 17 | 3553024 | 99.1 |
| mlcp2 | (K-12)<br>EC042_1742 | AAAGCCCGAAAAATGTGCTGTTAATCACATGCCTAAGTAAAAATTTGACGACACGTATTGAAGTGCTTCACCATAGCCTACAGATTATTTCGGAGCGCGAAAATATAGGGAG<br>AAAGCCCGAAAAATGTGCTGTTAATCACATGCCTAAGTAAAAATTTGACGACACGTATTGAAGTGCTTCACCATAGCCTACAGATTATTTCGGAGCGCGAAAATATAGGGAG | -70.5 | 37 | 58 | 15 | 1668591 | 100 |
| nagB | (K-12)<br>EC042_0706 | AACGCGTGTCTTTTGTGAGTTTTGTCACCAAATATCGTTATTATCACTCCCTTTTACTGGCTAAACCAGAAAACTTATTTTATCATTCAAAAAATCAGGTCGGATTGACGCC<br>AACGCGTGTCTTTTGTGAGTTTTGTCACCAAATATCGTTATCATCACTCCCTTTTACTGGCTAAACCAGAAAACTTATTTTATCATTCAAAAAATCAGGTCGGATTGACGCC | -70.5 | 37 | 60 | 17 | 703708 | 99.1 |
| ascF | (K-12)<br>EC042_2908 | AAAATATTCAGGTGACCGGTTTCACAAATATAAAAAATGAACAATTCACTCTCTTGCTTATTTAGTGACAACTATTCATGATTTTGTGAAACCGGTTTCTTAATTCCGTT<br>AAAATATTCAGGTGACCGGTTTCACAAATATAAAAAATGAACAATTAACTCTCTTGCTTATTTAGTGACAACTATTCATGATTTTATGAAACCGGTTTCTTAATTCCGTT | -71.5 | 36 | 59 | 17 | 2839429 | 98.2 |
| feaBp1 | (K-12) | TATTGTCACGATTTGCGGAGCTTGTCACAGCTGACAAAGCGAATGTCACAGCGAAAAAAGTGACTTTTCTTGTCGCTGCGTACACTGAAATCACACTGGGTAAATAATAAGGA | -71.5 | 39 | 60 | 15 | 1447492 | |
| flhD | (K-12)<br>EC042_2058 | ATTGAGTGTTTTGTGTGATCTGCATCACGCATTATTGAAAATCGCAGCCCCCCTCCGTTGTATGTGCGTGTAGTGACGAGTACAGTTGCGTCGATTTAGGAAAAATCTTAGAT<br>ATTGAGTGTTTTGTGTGATCTGCATCACACATTATTGAAAATCGCAGCCCCCCTCCGTTGTATGTGCGTGTAGTGACGAGTACAGTTGCGTCGATTTAGGAAAAATCTTAGAT | -71.5 | 37 | 60 | 17 | 1978395 | 99.1 |
| glnAp1 | (K-12)<br>EC042_4243 | TTGCAGAGTCCCTTTGTGATCGCTTTCACGGAGCATAAAAAGGGTTATCCAAAGGTCATTGCACCAACATGGTGCTTAATGTTTCCATTGAAGCACTATATTGGTGCAACATT<br>TTGCAGAGTCCCTTTGTGATCGCTTTCACGGAGCATAAAAAGGGTTATCCAAAGGTCATTGCACCAATATGGTGCTTAATGTTTCCATTGAAGCACTATATTGGTGCAACATT | -71.5 | 34 | 56 | 16 | 4058222 | 99.1 |
| gntTp | (K-12)<br>EC042_3676 | TTCCCCGCGATAATATGACCAACCTCTCATAATTTAAATTTACCCCGCTCTGGTGATTCTCAAACGCCAGATGTTACCCGTATCATTCACATGGGTACCAAACATACTCCTGA<br>TTCCCCGCGATAATATGACCAACCTCTCATAATTTAAATTTACCCCGCTCTGGTGATTCTCAAACGCCGGATGTTACCCGTATCATTCACGTGGGTACCAAACATACTCCTGA | -71.5 | 36 | 60 | 18 | 3546405 | 98.2 |
| murQp1 | (K-12)<br>EC042_2637 | CCGCTTAAGTTCTTATGACGCTCTTCACACTCTGCGAGTGTAGACTCAATGATTCCTTTAAGCCTGTCAATTCGGGATATATGATTCCATACATCTGCAATTATTTGATCGTA<br>CCGCTTAAGTTCTTATGACGCTCTTCACACTCTGCGAGTGTAGACTCAATGATTCCTTTAAGTCTGTCAATTCGGGATATATGATTCCATACATCTGCAATTATTTGATCGTA | -71.5 | 37 | 59 | 16 | 2545733 | 99.1 |
| dadAp3 | (K-12) | AGAGTCAGGGAGATGTGAGCCAGCTCACCATAAAAAAGCCGCATGTTGAATAATATTTTCAACTGAGTTATCAAGATGTGATTAGATTATTATTCTTTTACTGTATCTACCGTT | -72.5 | 37 | 62 | 19 | 1237538 | |
| rpoHp3 | (K-12) | TCTTCCCGGTATTTCATCTCTATGTCACATTTTGTGCGTAATTTATTCACAAGCTTGCATTGAACTTGTGGATAAAATCACGGTCTGATAAAACAGTGAATGATAACCTCGTTGC | -73.5 | 39 | 62 | 17 | 3600870 | |
| tsx | (K-12) | GAATGTGTGTAAACGTGAACGCAATCGATTACGTAAATGATAGAACTGTGAAACGAAACATATTTTTGTGAGCAATGATTTTTATAATAGGCTCCTCTGTATACGAAATATTTAG | -73.5 | 39 | 62 | 17 | 432091 | |
| yaeQ | (K-12)<br>EC042_0189 | ACTCTGTTTCGCAAGGTGACTATACCACACTCATTTCTGCAATATCAGCGCCGCAACTGCACGTATTCCGTTACAATGGCCTCCTGATTCGAAAGGAGTTTTCTTATGGCGCTTA<br>ACTCTGTTTCGCAAGGTGACTATACCACACTCATTTCTGCAATATCAGCGCCGCAACTGCACGTATTCCGTTACAATGGCCTCCTGATTCGAAAGGAGTTTTCTTATGGCGCTTA | -73.5 | 42 | 65 | 17 | 214280 | 100 |
| ompR | (K-12)<br>EC042_3666 | CGCACATTGGGTATAACGTGATCATATCAACAGAATCAATAATGTTTCGCCGAATAAATTGTATACTTAAGCTGCTGTTTAATATGCTTTGTAACAATTTAGGCTGAAATTCATAC<br>CGCACATTGGGTATAACGTGATCATATCAACAGAATCAATAATGTTTCGCCGAATAAATTGTATACTTAAGCTGCTGTTTAATATGCTTTGTAACAATTTAGGCTGAAATTCATAC | -74.5 | 38 | 62 | 18 | 3536685 | 100 |
| malE | (K-12)<br>EC042_4403 | TTACCGCCAATTCTGTAACAGAGATCACACAAAGCGACGGTGGGGCGTAGGGGCAAGGAGGATGGAAAGAGGTTGCCGTATAAAGAAACTAGAGTCCGTTTAGGTGTTTTCACGAGCA<br>TTACCGCCAATTCTGTAACAGAGATCACACAAAGCAACGGTGGGGCGTAGGGGCAAGGAGGATGGAAAGAGGTTGCTGTATAAAGAAACTAGAGTCCGTTTAGGTGTTTTCACGAGCA | -76.5 | 42 | 64 | 16 | 4246464 | 98.3 |

| | | | | | | | | |
|---|---|---|---|---|---|---|---|---|
| *putP* | (K-12)<br>EC042_1090 | CAACCTGATGAAAAATAGTGTCGCTGAGCACTAAAATTTAATGTAAATGGTGTGTTAAATCGATTGTGAATAACCAGCGCTTCCGGCAGGATACGGTCGCCCTGGTAAAACATAAACT<br>CAACCTGATGAAAAATAGTGTCGCTGAGCACTAAAATTTAATGTAAATGGTGTGTTAAATCGATTGTGAATAACCAGCGCTTCCGGCAGGATACGGTCGCCCTGGTAAAACATAAACT | -76.5 | 43 | 66 | 17 | 1079210 | 100 |
| *furpa* | (K-12) | TTGCCGTTGTAAATGTAAGCTGTGCCACGTTTTATTAACAATATTTGCCAGGGACTTGTGGTTTTCATTTAGGCGTGGCAATTCTATAATGATACGCATTATCTCAAGAGCAAATTCT | -77.5 | 44 | 65 | 17 | 710716 | |
| *murQp2* | (K-12) | CCGCTTAAGTTCTTATGACGCTCTTCACACTCTGCGAGTGTAGACTCAATGATTCCTTTAAGCCTGTCAATTCGGGATATATGATTCCATACATCTGCAATTATTTGATCGTAAATAGTAA | -79.5 | 43 | 67 | 18 | 2545741 | |
| *serA* | (K-12)<br>EC042_3123 | CCCCCGTTAAAAAAATTCTCTTCATTAAATTTGGTGACATGTGTCACGCTTTTACCAGGCAATTGTCGATTGCTCTAAATAAATCCTCTAAACCAGCATATTCATCCAAGAATTACCTTTG<br>CCCCCGTTAAAAAAATTCTCTTCATTAAATTTGGTGACATGTGTCACGCTTTTACCAGGCAATTGTCGATTGCTCTAAATAAATCCTCTAAACCAGCATATTCATCCAAGAATTACCT | -79.5 | 43 | 68 | 19 | 3058549 | 100 |
| *fucP* | (K-12) | TTTCGATTATTAAAGTGATGGTAGTCACATAAAGTCACCTTCTAGCTAATAAGTGTGACCGCCGTCATATTACAGAGCGTTTTTTATTTGAAAATGAATCCATGAGTTCATTTCAGACAGGC | -80.5 | 43 | 69 | 20 | 2934119 | |
| *hofM* | (K-12)<br>EC042_3656 | GGCGCTGTAAATCTGCATCGGAATTTGCAGGCGAACATCTTTTAATGTCGCCACATCCGGCAGTTGTGGCTCGATGTAGCGGTATAGGCCATAAATCGAGCCTGCTCCCAGCAGAATGCAAC<br>GGCGCTGTAAATCTGCATCGGAATTTGCAGGCGAACATCTTTTAATGTCGCCACATCCGGCAGTTGTGGCTCGATGTAGCGGTATAGGCCATAAATCGAGCCTGCTCCCAGCAGAATG | -80.5 | 43 | 69 | 20 | 3522928 | 100 |
| *rpoHp4* | (K-12) | TCTTCCCGGTATTTCATCTCTATGTCACATTTTGTGCGTAATTTATTCACAAGCTTGCATTGAACTTGTGGATAAAATCACGGTCTGATAAAACAGTGAATGATAACCTCGTTGCTCTTAAG | -80.5 | 45 | 68 | 17 | 3600863 | |
| *xylE* | (K-12) | GATAATTATCACAATTAAGATCACAGAAAAGACATTACGTAAACGCATTGTAAAAAATGATAATTGCCTTAACTGCCTGACAATTCCAACATCAATGCACTGATAAAAGATCAGAATGG | -80.5 | 43 | 69 | 20 | 4242291 | |
| *melA* | (K-12)<br>EC042_4485 | TAAACTCAGATTTACTGCTGCTTCACGCAGGATCTGAGTTTATGGGAATGCTCAACCTGGAAGCCGGAGGTTTTCTGCAGATTCGCCTGCCATGATGAAGTTATTCAAGCAAGCCAGGAGATC<br>TAAACTCAGATTTATTGATGCTTCACGTAGGATCTGAGTTTATGGGAATGCTCAACCTGGAAGCCGGAGGTTTTCTGCAGATTCGCCTGCCATGATGAAGTTATTCAAGCAAGCCAGG | -81.5 | 49 | 70 | 15 | 4341887 | 97.6 |
| *pkap* | (K-12) | GGCGAGCATTTTACCCGCTTTAAAACACGCTATCTGGCAGGAAAAACGCAACATCTGGGTAGCATCACAGCAGAACAGTTAGAAAGCGTTTAAAATCATTCGGTCACTTCTGCGGGAGACCGG | -81.5 | 47 | 70 | 17 | 2719931 | |
| *xylA* | (K-12)<br>EC042_3871 | TTCCATTTTATTTTGCGAGCGAGCGCACACTTGTGAATTATCTCAATAGCAGTGTGAAATAACATAATTGAGCAACTGAAAGGGAGTGCCCAATATTACGACATCATCCATCACCCGCGGCAT<br>TTCCATTTTATTTTGCGAGCGAGCGCACACTTGTGAATTATCTCAATAGCAGTGTGAAATAACATAATTGAGCAACTGAAAGGGAGTGCCCAATATTACGACATCATCCATCACCCGC | -81.5 | 47 | 70 | 17 | 3730807 | 100 |
| *sxy* | (K-12)<br>EC042_1045 | TCAACCCTATTTTCGGAAAGAGCCTCGCAAATTTTGTCGTTGGGTGACGGGAAAACATAAATTAATCTTGCCCCTTAAGAATAAGTTGCCTATTTTCGTAGTTAACGGATCCGTTAATGTGAAT<br>TCAACCCTATTTTCGGAAAGAGCCTCGCAAATTTTGTCGTTGGTGACGGGAAAACATAAATTAATCTTGCCCCTTAAGTATAAGTTGCCTATTTTCGTAGTTAACGGATCCGTTAATG | -82.5<br>**(-81.5)** | 47<br>**(46)** | 70<br>**(69)** | 17 | 1021083<br>**(-1)** | 98.4 |
| *chbB* | (K-12)<br>EC042_1903 | TATTTGCCCGAAATGTGAAGAGGGTCATAACCACAGGTCAAGGAGAAACAATTTATAAGGTCAAAGAAATACTATTGCTCAGGTCTATACCGTATACTCCTTTCAGCCACAAAAAAAGTCATGTT<br>TATTTGCCCGAAATGTGAAGAGGGTCATAACCGCAGGTCAAGGAGAGACAATTTATAAGGTCAAAGAAATACTATTGCTCAGGTCTATACCGTATACTCCTTTCAGCCACAAAAAAAG | -83.5 | 49 | 72 | 17 | 1821726 | 98.4 |
| *cspD* | (K-12)<br>EC042_0972 | TTCCTGTCAATAGCGTTAACTGCTTCAAATTTTTGATTCATACCTGCCCCTTGTCCCCTGTCAAATGCTTGACGGCTCGCCCTAATTCTCTAAATTGTATTTCTAGAGTTGGCGAGGTTTTGAAC<br>TTCCTGTCAATAGCGTTAACTGCTTCAAATTTTTGATTCATACCTGCCCCCTGGCCCCTGTCAAATGCTTGACGGCTCGCCCTAATTCTCTAAATTGTATTTCTAGAGTTGGCGAGGT | -83.5 | 48 | 70 | 16 | 922676 | 98.4 |
| *hupBp3* | (K-12)<br>EC042_0478 | GTGACTGCAAAATAGTGACCTCGCGCAAAATGCACTAATAAAAACAGGGCTGGCAGGCTAATTCGGGCTTGCCAGCCTTTTTTTGTCTCGCTAAGTTAGATGGCGGATCGGGCTTGCCCTTATTA<br>GTGACTGCAAAATAGTGACCTCGCGCAAAATGCACTAATAAAAACAGGGCTGGCAGGCTAATTCGGGCTTGCCAGCCTTTTTTTGTCTCGCTAAGTTAGATGGCGGATCGGGCTTGCC | -83.5 | 48 | 71 | 17 | 461332 | 100 |
| *sdhC* | (K-12)<br>EC042_0739 | TAAATGTTGTTATCGTGACCTGGATCACTGTTCAGGATAAAACCCGACAAACTATATGTAGGTTAATTGTAATGATTTTGTGAACAGCCTATACTGCCGCCAGGTCTCCGGAACACCCTGCAATC<br>TAAATGTTGTTATCGTGACCTGGATCACTGTTCAGGATAAAACCCGACAAACTATATGTAGGTTAATTGTAATGATTTTGTGAACGGCCTATACTGCCGCCAGGTCTCCGGAACACCC | -83.5 | 46 | 71 | 19 | 754958 | 99.2 |
| *spf* | (K-12) | AGCTGTCACGTTTTGTGATGGCTATTAGAAATTCCTATGCAACAACTGAAAAAAAATTACAAAAGTGCTTTCTGAACTGAACAAAAAAGAGTAAAGTTAGTCGCGTAGGGTACAGAGGTAAGATG | -84.5 | 50 | 72 | 16 | 4049899 | |
| *serC* | (K-12)<br>EC042_0997 | AGAGATTCTTTTGTGTGATGCAAGCCACATTTTTGCCCTCAACGGTTTACTCATTGCGATGTGTGTCACTGAATGATAAAACCGATAGCCACAGGAATAATGTATTACCTGTGGTCGCAATCGATTG<br>AGAGATTCTTTTGTGTGATGCAAGCCACATTTTTGCCCTCAACGGTTTACTCATTGCGATGTGTGTCACTGAATGATAAAACCGATAGCCACAGGAATAATGTATTACCTGTGGTCG | -85.5<br>**(-86.5)** | 52<br>**(53)** | 74<br>**(75)** | 16 | 957595<br>**(+1)** | 99.2 |
| *dsdX* | (K-12) | TTCACCTTCATTTTTAGATGTAAATCACTCCATTGATGCAATTTACCTCATGTGAAAGGCAAATTTTATCGTTTGTCAGCCTGCGTTGTTTTTTTGTCCAATATCATCAGGTTAATCACAGGGGAAGGT | -87.5 | 52 | 75 | 17 | 2477821 | |
| *casA* | (K-12) | GTATGAATATTTTATGTAATAAAATTCATGGTAATTATTATAACTAAAAGTTTCTTTAATAATAAAACGAATAACTTGCAGATTTGAAATGCATGCATTATTGTCTTTAAACAATTCAACACATCTTAATA | -89.5 | 55 | 78 | 17 | 2884319 | |
| *cstAp1* | (K-12)<br>EC042_0636 | ACTCGGTTAACGGAGTGATCGAGTTAACATTGTTAAGTTAAATATTGGTTTCAACTCCGATTTACATGGTTGCTGTGTTGTTAAATTGTACAAAGATGTTATAGAAACAAAATGTAACATCTCTATGGACA<br>ACTCGGTTAACGGAGTGATCGAGTTAACATTGTTAAGTTAAATATTGGTTTCAACTCCGATTTACATGGTTGCTGTGTTGTTAAATTGCGCAAAGATGTTATAGAAACAAAATGTAAC | -89.5 | 53 | 79 | 20 | 629855 | 98.5 |
| *nupC* | (K-12) | AACATAGCAGAAATGTATGACAGATCACTATTTTTGAAGCCTGTCACAGGACGTCATTATAGTGTGTGTCAGATCTCGTTTTCCTTAACCATGTTACATAGAATGTGCACGGAAATTTAACCTGCCTCATA | -89.5 | 52 | 78 | 20 | 2513009 | |
| *pck* | (K-12)<br>EC042_3664 | CATTCAGGAATGCGATTCCACTCACAATATTCCCGCCATATAACCAAGATTTAACCTTTTGAGAACATTTTCCACACCTAAAATGCTATTTCTGCGATAATAGCAACCGTTTCGTGACAGGAATCACG<br>CATTCAGGAATGCGATTCCACTCACAATATTCCCGCCACATAAACCAAGATTTAACCTTTTGAGAACATTTTCCACACCTAAAATGCTATTTCTGCGATAATAGCAACCGTTTCG | -89.5<br>**(-90.5)** | 52<br>**(53)** | 78<br>**(79)** | 20 | 3532679<br>**(+1)** | 98.4 |
| *uidA* | (K-12)<br>EC042_1785 | TTTTAAAGATTAATGCGATCTATATCACGCTGTGGGTATTGCAGTTTTTGGTTTTTTGATCGCGGTGTCAGTTCTTTTTATTTCCATTTCTCTTCCATGGGTTTCTCACAGATAACTG<br>TTTTAAAGATTAATGCGATCTATATCACGCTGTGGGTATTGCAGTTTTTGGTTTTTTGATCGCGGTGTCAGTTCTTTTTATTTCCATTTCTCTTCCATGGGTTTCTCACAGATAACTG | -89.5 | 51 | 75 | 18 | 1696152 | 100 |
| *araJ* | (K-12)<br>EC042_0428 | CAGCAAACTGGAAAGTACGTTTGCAGTGAAATAACTATTCAGCAGGATAATGAATACAGAGGGGCGAATTATCTCTTGGCCTTGCTGGTCGTTATCCTGCAAGCTATCACTTTATTGGCTACGGTGATTGGT<br>CAGCAAACTGGAAAGTGTGTTTGCAGTGAAATAACTATTCAGCAGGATAATGAATACAGAGGGGCGAATTATCTCTTGGCCTCGCTGGTCGTTATCCTGCAAGCTATCACTTTATTGG | -90.5 | 55 | 79 | 18 | 412529 | 97 |
| *aspA* | (K-12)<br>EC042_4615 | GTAATCCCAAAGCGGTGATCTATTTCACAAATTAATAATTAAGGGGTAAAAACCGACACTTAAAGTGATCCAGATTACGGTAGAAATCCTCAAGCAGCATATGATCTCGGGTATTCGGTCGATGCAGGGGAT<br>GTAATCCCAAAGCGGTGATCTATTTCACAAATTAATAATTAAGGGGTAAAAACCGACACTTAAAGTGATCCAGATTACGGTAGAAATCCTCAAGCAGCATATGATCTCGGGTATTCGG | -90.5 | 54 | 79 | 19 | 4368432 | 100 |
| *glpA* | (K-12) | TTCATAAATTAAATGTGAATTGCCGCACACATTATTAAATAAGATTTACAAAATGTTCAAAATGACGCATGAAATCACGTTTCACTTTCGAATTATGAGCGAATATGCGCGAAATCAAACAATTCATGTTTT | -90.5 | 55 | 80 | 19 | 2352583 | |
| *ansBp2* | (K-12)<br>EC042_3164 | TAATTCTTCGTTTTGTTACCTGCCTCTAACTTTGTAGATCTCCAAAATATATTCACGTTGTAAATTGTTTAACGTCAAATTTCCCATACAGAGCTAAGGGATAATGCGTAGCGTTCACGTAACTGGAGGAATG<br>TAATTCTTCGTTTTGTTACCTGCCTCTAACTTTGTAGATCTCCAAAATATATTCACGTTGTAAATTGTTTAACGTCAAATTTCCCATACAGAGCTAAGGGATAATGCGTAGCGTTCAC | -91.5 | 53 | 79 | 20 | 3100751 | 100 |
| *glpT* | (K-12) | TTCGAAAGTGAAACGTGATTTCATGCGTCATTTTGAACATTTTGTAAATCTTATTTAATAATGTGTGCGGCAATTCACATTTAATTTATGAATGTTTTCTTAACATCGCGGCAACTCAAGAAACGGCAGGTTC | -91.5 | 53 | 80 | 21 | 2352451 | |
| *cdd* | (K-12) | ATGGGCTAAAATTTGCGATGCGTCGCGCATTTTTGATGTATGTTTCACGCGTTGCATAATTAATGAGATTCAGATCACATATAAAGCCACAACGGGTTCGTAAACTGTTATCCCATTACATGATTATGAGGCAA | -92.5 | 53 | 80 | 21 | 2231819 | |
| *manX* | (K-12) | CTTTGCAAACGAATGTGACAAGGATATTTTACCTTTCGAAATTTCTGCTAATCGAAAGTTAAATTACGGATCTTCATCACATAAAATAATTTTTTCGATATCTAAAATAAATCGCGAAACGCAGGGGTTTTTGG | -92.5 | 55 | 82 | 21 | 1901933 | |
| *nupG* | (K-12) | TCAGGGGCAAAAATGTTATCCACATCACAATTTCGTTTTGCAAATTGGGAATCTTTGCAATTATTTGCCACAGGTAACAAAAAACCAGTCCGCGAAGTTGATAGAATCCCATCATCTCGCACGGTCAAATGTGC | -92.5 | 54 | 81 | 21 | 3105651 | |
| *ompF* | (K-12)<br>EC042_1020 | AAGTTCCTTAAATTTTACTTTTGGTTACATATTTTTTCTTTTTGAAACCAAATCTTTATCTTTGTAGCACTTTCACGGTAGCGAAACGTTAGTTTGAATGGAAAGATGCCTGCAGACACATAAAGACACCAAAC<br>AAGTTCCTTAAATTTTACTTTTGGTTACATATTTTTTCTTTTTGAAACCAAATCTTTATCTTTGTAGCACTTTCACGGTAGCGAAACGTTAGTTTGAATGGAAAGATGCCTGCAGACA | -92.5 | 57 | 81 | 18 | 987092 | 100 |
| *rhaB* | (K-12)<br>EC042_4277 | CAATTCAGCAAATTGTGAACATCATCACGTTCATCTTTCCCTGGTTGCCAATGGCCCATTTTCCTGTCAGTAACGAGAAGGTCGCGAATTCAGGCGCTTTTTAGACTGGTCGTAATGAAATTCAGCAGGATCAC<br>CAATTCAGCAAATTGTGAACATCATCACGTTCATCTTTCCCTGGTTGCCAATGGCCCATTTTCCTGTCAGTAACGAGAAGGTCGCGAAGTCAGGCGCTTTTTAGACTGGTCGTAATGA | -92.5 | 60 | 81 | 15 | 4097472 | 99.3 |
| *rhaS* | (K-12)<br>EC042_4278 | GCTCACCGCATTTCCTGAAAATTCACGCTGTATCTTGAAAAATCGACGTTTTTACGTGGTTTTCCGTCGAAAATTTAAGGTAAGAACCTGACCTCGTGATTACTATTTCGCCGTGTTGACGACATCAGGAGGC<br>GCTTACCGCATTTCCTGAAAATTCACGCTGTATCTTGAAAAATCGACGTTTTTACGTGGTTTTCCGTCGAAAATTTAAGGTAAGAACCTGACCTCGTGATTACTATTTCGCCGTGTT | -92.5 | 60 | 81 | 15 | 4097711 | 99.2 |

[a]Promoter sequences are derived from RegulonDB(11) and EcoCyc(13). The annotated CRP site is highlighted yellow, the -35 element is highlighted grey, the -10 element is highlighted cyan, and the transcription start site is highlighted green.

[b]Differences in the promoter sequence found in the 042 genome are in given in parenthesis and are in bold.

[c]The percentage of nucleotide identity across the given sequence.

#### Supplementary material references
